# A high-affinity calmodulin-binding site in the CyaA toxin translocation domain is essential for invasion into eukaryotic cells

**DOI:** 10.1101/2020.09.14.296129

**Authors:** Alexis Voegele, Mirko Sadi, Darragh P O’Brien, Pauline Gehan, Dorothée Raoux-Barbot, Maryline Davi, Sylviane Hoos, Sébastien Brûlé, Bertrand Raynal, Patrick Weber, Ariel Mechaly, Ahmed Haouz, Nicolas Rodriguez, Patrice Vachette, Dominique Durand, Sébastien Brier, Daniel Ladant, Alexandre Chenal

## Abstract

The molecular mechanisms and forces involved in the translocation of bacterial toxins into host cells have thus far remained elusive. The adenylate cyclase (CyaA) toxin from *Bordetella pertussis* displays a unique intoxication pathway in which its catalytic domain is directly translocated across target cell membranes. We have previously identified a translocation region in CyaA that contains a segment, P454 (residues 454–484), exhibiting membrane-active properties related to antimicrobial peptides. Herein, we show that this peptide is able to translocate across membranes and interact with calmodulin. Structural and biophysical analyses have revealed the key residues of P454 involved in membrane destabilization and calmodulin binding. Mutational analysis demonstrated that these residues play a crucial role in CyaA translocation into target cells. We have also shown that calmidazolium, a calmodulin inhibitor, efficiently blocks CyaA internalization. We propose that after CyaA binding to target cells, the P454 segment destabilizes the plasma membrane, translocates across the lipid bilayer and binds calmodulin. Trapping of the CyaA polypeptide chain by the CaM:P454 interaction in the cytosol may assist the entry of the N-terminal catalytic domain by converting the stochastic process of protein translocation into an efficient vectorial chain transfer into host cells.

## Introduction

The adenylate cyclase (CyaA) toxin is a major virulence factor produced by *Bordetella pertussis*, the causative agent of whooping cough, and is involved in the early stages of respiratory tract colonization ^1-5^. CyaA, a 1706-residue long protein (Figure S1), is a Repeat-in-ToXin (RTX) ^6-10^ multi-domain toxin ^11-12^. Once secreted by *B. pertussis*, CyaA invades eukaryotic cells through an original molecular mechanism that involves a direct translocation of its N-terminal adenyl cyclase catalytic (AC) domain across the plasma membrane. The ATP-cyclizing, calmodulin-activated AC domain comprises the first 364 N-terminal residues ^13-16^. The translocation region (TR, residues 365 to 527) is essential for AC translocation into target cells ^17^. The hydrophobic region (HR, residues 528 to 710) inserts into the cell membrane and makes cation-selective pores ^10, 18-19^; the acylation region (AR, residues 711 to 1005) contains two post-translational acylation sites, at lysines K860 and K983 ^20-22^, required for the refolding of the CyaA toxin ^12, 23^ and AC translocation across membranes *in vivo* and *in vitro* ^5, 20-21, 24^. The cell-receptor binding domain of CyaA (RD, residues 1006 to 1706) is made up of approximately 40 copies of calcium-binding RTX motifs ^8-10, 25^.

The dedicated type 1 secretion system (T1SS), made of CyaB, CyaD and CyaE proteins ^26-27^, recognizes a secretion signal located at the C-terminal extremity of CyaA. Once secreted through the T1SS, the toxin binds calcium and folds in the extracellular milieu. A calcium-induced disorder-to-order transition of the RTX motifs occurs upon CyaA secretion from the low-calcium concentration of the bacterial cytosol to the calcium-rich extracellular environment ^7-8, 25, 28-40^. CyaA folding is an acylation-dependent and calcium-driven sequential process ^12, 23, 41^.

Invasion of target cells occurs via a unique process among known bacterial toxins. First, RD binds with high affinity to a cell receptor, the CD11b/CD18 integrin that is expressed by a subset of leukocytes (neutrophils, dendritic cells (DC) and macrophages) ^42-50^. CyaA can also intoxicate cells lacking CD11b/CD18 by directly interacting with the target cell membrane, although with a reduced efficiency ^12, 51-54^. CyaA then inserts into the membrane of target cells via its hydrophobic domains and AC is directly translocated across the plasma membrane into the cytoplasm in a calcium and membrane-potential dependent manner ^17, 24, 46, 55-57^.

Inside the cell, AC binds to calmodulin (CaM) that stabilizes the catalytic site into its active state ^15-16, 58^ to convert ATP into cAMP with a high catalytic turnover (> 1000 /s). Accumulation of supraphysiologic levels of cAMP in target cells alters their phagocytic functions leading to host defense subversion ^5, 11, 59-60^.

Experimental results from the past few decades have provided direct evidence that AC translocation requires CyaA acylation, a calcium gradient, and a membrane potential across the plasma membrane ^21, 24, 61-62^. However, the molecular mechanism and forces involved in the translocation of the CyaA catalytic domain across the plasma membrane have thus far remained elusive. We have previously shown that a peptide, corresponding to the C-terminus of the translocation region TR of CyaA, P454 (residues 454–484, Figure S1), exhibits membrane-active properties related to antimicrobial peptides (AMPs) ^17, 63^: this peptide adopts a helical conformation upon membrane interaction and induces a local destabilization of the lipid bilayer ^63-66^. This property is likely essential for CyaA as deletion of the TR region (residues 384–489) encompassing the P454 segment, selectively abrogates the ability of the modified toxin to intoxicate target cells ^17^.

Here, we show that P454 is able to translocate across a lipid bilayer and binds with high affinity to calcium-loaded calmodulin (holo-CaM). We present structural models and crystal structures of the P454 peptide in complex with holo-CaM, and identified in P454 the amino acid residues that are critical for CaM-binding, membrane interaction and destabilization. Modifications of these residues within the full-length CyaA toxin are sufficient to fully and specifically abrogate the translocation of the catalytic domain across the cell membrane. Finally, we show that calmidazolium, a high-affinity CaM inhibitor, specifically blocks translocation of the AC domain into eukaryotic cells. We propose that once CyaA is inserted into the target cell membrane, the P454 segment can interact with the plasma membrane and destabilize the lipid bilayer, favoring its translocation across the lipid bilayer into the cytosol where it binds CaM. Trapping of the CyaA polypeptide chain by the CaM:P454 interaction may thus assist the irreversible translocation of the N-terminal AC domain. Therefore, CaM is not only a key activator of the catalytic activity of CyaA inside cells, but also acts as an essential cytosolic binder of the CyaA translocation region able to grab the polypeptide chain to favor its entry into target cells.

## Material and methods

Material and methods are described in the supplementary information file.

## Results

### 1 The P454 peptide from the CyaA toxin binds to calmodulin

We have previously shown that the P454 peptide, residues 454-484 from the CyaA translocation region, exhibits membrane-active properties similar to antimicrobial peptides (AMPs) ^67^. As with certain AMPs, the P454 peptide displays biophysical properties that are similar to that of many calmodulin-binding peptides: they can form amphiphilic helices, they are positively charged and contain a few aromatic or hydrophobic residues involved in complex stabilization ^68-70^. Indeed, we found that P454 binds calmodulin (CaM) in solution in a calcium-dependent manner as shown by analytical ultracentrifugation and far-UV circular dichroism spectroscopy (Figure S2 and Table S1). Analysis of the thermodynamic parameters of the P454:CaM complex formation by isothermal titration calorimetry (ITC) revealed a calcium-dependent interaction with a dissociation constant of about 90 nM at 25°C (ΔG_Kd_=- 9.6 kcal/mol) and a P454:CaM stoichiometry of 1:1 (Figure S3A-B, Figure S4 and Table S2). No binding could be detected in the absence of calcium (Figure S3A-B).

The affinity of P454 for CaM is much higher than that for lipid membranes. Indeed, the dissociation constant K_d_ of the P454:membrane equilibrium (calculated from the partition coefficient Kx = 790000) is ≈ 70 µM (ΔG_Kx_ = -8 kcal/mol, Table S3), about three orders of magnitude higher than that for CaM. Therefore, based on the free energy difference, ΔΔG, P454 should preferentially bind to CaM rather than interact with the membrane. This was confirmed by solution-to-membrane partition of P454 measured by fluorescence (Figure S5A and S5B): P454 progressively partitioned from buffer to membranes as lipid concentration increased. A shift of fluorescence polarity was observed upon addition of calcium-loaded calmodulin (holo-CaM), indicating that a P454:CaM complex was formed whatever the lipid concentration. Holo-CaM is converted into apo-CaM upon EDTA addition, leading to its dissociation from P454 that can then interact again with membranes (Figure S5).

The preferential binding of P454 to calmodulin over membrane was confirmed by a lipid vesicle permeabilization assay ^63, 65^. In this set up, addition of P454 peptide to ANTS:DPX loaded vesicles induces membrane permeabilization leading to a dye efflux that is monitored by fluorescence increase as a result of ANTS dequenching. Addition of holo-CaM at an early stage of membrane permeabilization immediately stopped the P454-induced dye efflux (Figure S5C). This suggests that holo-CaM selectively binds P454 leading to a displacement of the peptide from the membrane and an arrest of the P454-induced vesicle permeabilization. Calcium chelation by EDTA triggered dissociation of the P454:CaM complex and release of the P454 peptide that could partition back into membranes to resume permeabilization of the vesicles, leading to ANTS fluorescence recovery (Figure S5C). Taken together, these experiments show that P454 is a calcium-dependent calmodulin-binding peptide and that holo-CaM can efficiently antagonize the P454 interaction with membrane. These results prompted us to evaluate the intrinsic propensity of P454 to translocate across a lipid bilayer, in particular if holo-CaM would be asymmetrically present on the *trans* side of the membrane.

### 2 P454 translocation across lipid bilayers

We investigated the ability of P454 to translocate across membranes using the droplet interface bilayers (DIB) approach ^71^. The *cis* droplet population contains the dye-labeled peptide TAMRA-P454 while the *trans* droplet population is prepared in the presence or absence of holo-CaM. After mixing and random formation of pairs of droplets, a lipid bilayer is formed at the interface between two adhering droplets (Figure S6). We measure the transfer of fluorescence from a *cis* fluorescent droplet to a *trans* non-fluorescent droplet to reveal peptide translocation across the lipid bilayer formed at the droplet interface (Figure S6A). All dye-labeled peptides used in this study interact with membranes, as evidenced by the fluorescent rings staining the *cis* droplets at the beginning of the experiments. In the absence of calmodulin in the *trans* droplet, no increase of fluorescent P454 is measured in the volume of the *trans* compartment after 15 minutes of incubation (Figure 1 and Figure S6B). Conversely, in the presence of 5 µM of holo-CaM in the *trans* droplet, a significant amount of fluorescence is measured in the *trans* compartment (Figure 1 and Figure S6C). These results indicate that P454 is competent to translocate across membrane and to bind holo-CaM if present in the *trans* compartment.

**Figure 1.**
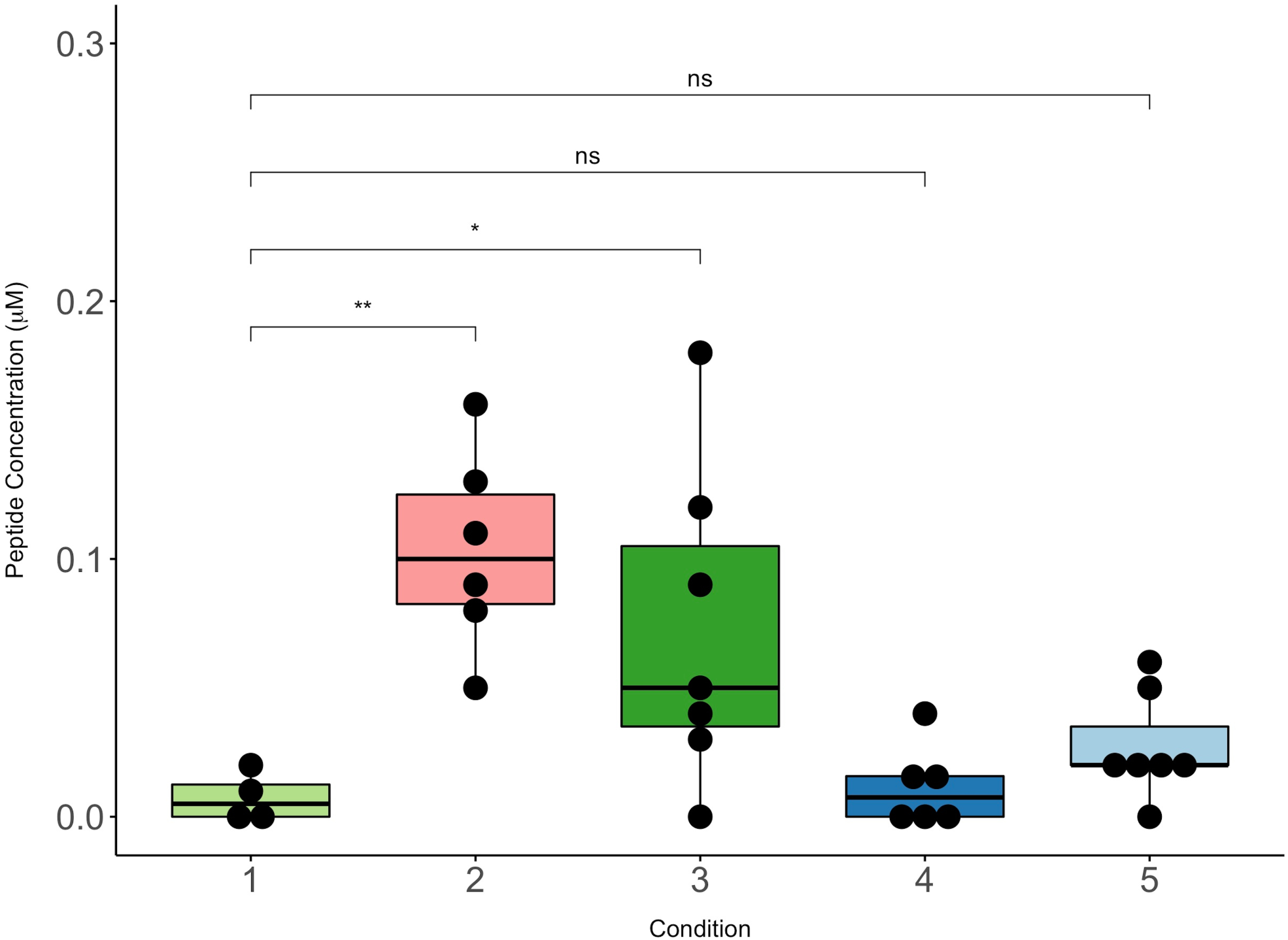
Peptide translocation across droplet interface bilayers. Boxplot representation of the TAMRA-peptide concentration (µM) in the *trans* droplet 15 min after the formation of the droplet interface bilayers (see Material and Methods for details). TAMRA fluorescence was measured in the absence (1) and in the presence of 5 μM CaM (2-5) in the *trans* droplet; in the absence (3) and in the presence (1-2,4-5) of a calcium gradient across the lipid bilayer (CaCl_2_: 2 mM *cis vs* 0.2 mM *trans*). Concentration of TAMRA-P454 WT (1-3), TAMRA-P454 R12E (4) and the TAMRA-H-helix (5) peptides in the *trans* droplets are reported. Five to seven independent trials were conducted for each condition. Mann-Whitney-Wilcoxon test was applied to compare the experiments (ns: p>0.05, *: p<0.05 and **: p<0.01).

We then assayed two other dye-labeled peptides: the first being the H-helix peptide, corresponding to the main binding site of AC (residues 233-254 of CyaA) to CaM and that is involved in adenyl cyclase activation ^16^. The second peptide is a P454-derived peptide in which the two arginine residues R461 (R*1*) and R474 (R*2*) were substituted by glutamate residues, hereafter designated P454_R*12*E_. This peptide exhibits a drastically reduced affinity for CaM 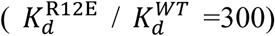 and a slightly decreased partitioning into membrane (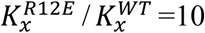, see below). Both P454_R*12*E_ and H-helix peptides interact with the membrane, as evidenced by the fluorescent ring located on the *cis* lipid leaflet observed at the beginning of the experiment. However, these peptides do not accumulate in the *trans* droplets containing 5 µM of holo-CaM even after 15 min of incubation (Figure 1). These results suggest that the H-helix peptide does not translocate across membrane in these experimental conditions, as this peptide should strongly interact with CaM (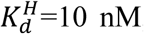 nM, Figure S7 and Table S4) and accumulate as a peptide:CaM complex in the *trans* compartment if translocation had occurred. The DIB experiment was also carried out with P454_R*12*E_ in the presence of 100 µM of CaM in the *trans* compartment (Figure S6D), i.e., at a concentration well above the K_d_ (22.7 µM) of the P454_R*12*E_:CaM complex formation (Table S3). In these conditions, a significant accumulation of P454_R*12*E_ in the *trans* compartment was measured, indicating that this peptide was able to translocate across the lipid bilayer and to bind CaM. Taken together, these data indicate that P454 interacts with the *cis* lipid leaflet of the membrane, translocates across the droplet interface bilayers, and forms a peptide:CaM complex in the *trans* compartment, as summarized in Figure S8.

### 3 Structure and dynamics of P454:CaM complex

We further characterized the interaction between P454 and CaM by an integrative structural biology approach. Initial attempts to crystallize the P454:CaM complex were unsuccessful. However, we obtained two distinct crystal forms of CaM in complex with a slightly shorter peptide, P458, corresponding to the residues 458-481 of CyaA (i.e. shorter than P454 by 4 residues (ASAH) at N-terminus and 3 residues (MTQ) at C-terminus), and that displays an affinity for CaM similar to that of P454 (K_d_^CaM^ = 240 nM, see Table S3). In both cases, several copies of CaM and P458 were present in the asymmetric unit (Table S5A). The superposition of these multiple copies yields an ensemble of P458:CaM conformations (Figures 2A and S9), which illustrates the well-documented conformational plasticity of CaM ^68-69, 72^ due to its central helix flexibility.

**Figure 2.**
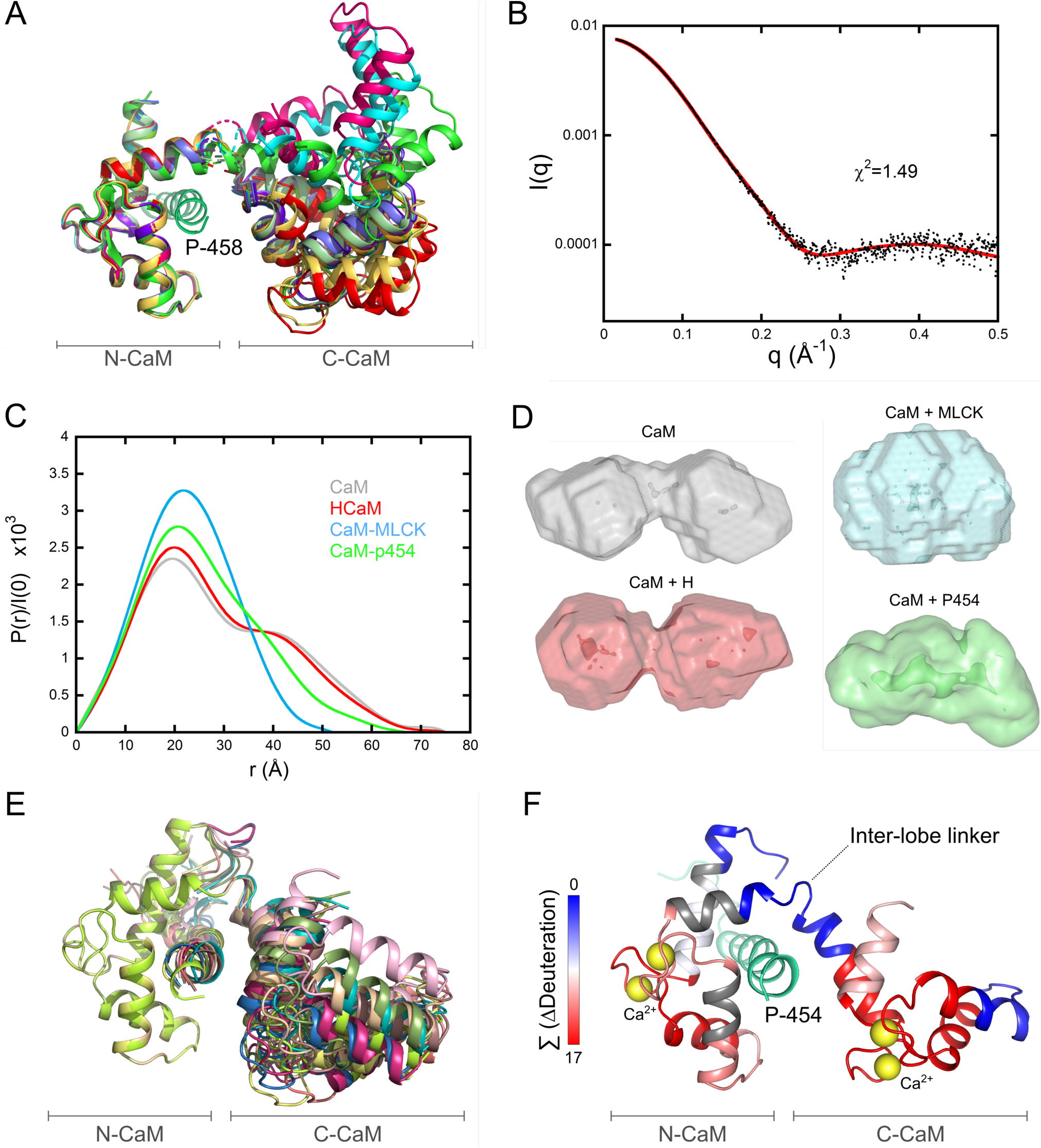
Structure and dynamics of the P454:CaM complex. **(A)** The twelve P458:CaM crystal structures (PDB 6YNS) are displayed after superimposition of C_α_s over the range 10 to 70 included, corresponding to the N-ter lobe of calmodulin. The crystal structure 1CLL {Chattopadhyaya, 1992 #1590} of the extended conformation of CaM is shown in light green. **(B)** Experimental SAXS curve of the P454:CaM complex (black dots) superimposed over the best fit (red curve) obtained from the structural model shown in Fig 2F. **(C)** Comparison of the four distance distribution functions obtained using the program GNOM for CaM alone (grey), H:CaM (red), MLCK:CaM (cyan) and P454:CaM (green) complexes. **(D)** DAMMIN models of CaM alone, H:CaM, MLCK:CaM and P454:CaM complexes, shown with the same color code. **(E)** Ten models fitting the SAXS curve shown on Fig 2B obtained using the program DADIMODO {Evrard, 2011 #1766} are displayed after superimposition of C_α_s over the range 10 to 70. **(F)** Effects of P454 on the HDX behavior of CaM. The uptake differences (ΔDeuteration) measured between the free- and P454-bound CaM were extracted for each peptide at each time point, summed, and plotted on the best-fitting structural model of P454:CaM (red curve in 2B). The summed ΔDeuteration values [Σ (ΔDeuteration)] are colored from blue (no variation of deuterium uptake) to red (major reductions of deuterium uptake). Uncovered regions are in grey.

The dynamics and the overall shape of the P454:CaM complex in solution were confirmed by SEC-SAXS measurements ^73^. The experimental SAXS pattern of the P454:CaM complex is shown in Figure 2B and the derived structural parameters are compared in Table S6 to that of the previously analyzed CaM complexes with either the H-helix peptide, corresponding to the main CaM binding segment from the AC domain or the MLCK peptide, corresponding to the CaM binding site from myosin light chain kinase ^16^. The distance distribution P(r) of the P454:CaM complex appears to be intermediate between that of free CaM and H:CaM complex on the one hand, and MLCK:CaM complex on the other (Figure 2C). *Ab initio* modeling yields a shape intermediate between the globular MLCK:CaM complex and the bi-lobed, extended shape of free CaM and H:CaM complex (Figure 2D), further exemplifying the conformational plasticity of CaM adapted to peptide ligand diversity.

The calculated SAXS curves from ten out of the twelve crystal structures of P458:CaM (PDB 6YNS) obtained are similar to, but slightly different from our experimental SAXS data, while two structures exhibit a different N- and C-domain arrangement and, accordingly, larger amplitude differences with experimental data. Using the model with the closest agreement, we added the six missing N-terminal amino-acids of CaM and the few missing terminal residues of P454 with Modeller to the P458:CaM X-ray structure. We then used this completed complex as a starting model to fit the SAXS data using the modeling program Dadimodo ^74-75^. Each run of the program yielded several models, the scattering pattern of which fitted our experimental SAXS data. After superimposition of the N-CaM domain, all resulting models appeared to exhibit close, but slightly different positions of the C-CaM domain (Figure 2E) that were similar to those observed within the crystal structures (Figure 2A).

Using HDX-MS ^76^, we compared the effect of P454 binding to CaM (Figures 2F, S10 and Table S7) with those observed following H-helix and MLCK peptide binding reported in our previous study ^16^. HDX-MS analysis reveals that the inter-lobe helix (residues 73-84) remains accessible in the presence and absence of all three peptides. The MLCK and H-helix peptides induce similar differences in deuterium uptake when bound to CaM as those observed in Figure S4 and S5 from ^16^. Interestingly, the deuterium uptake difference induced by P454 binding to CaM is significantly distinct from that observed with MLCK or H-helix, further highlighting the high conformational plasticity of CaM that is able to adapt to a wide diversity of peptide ligands. HDX-MS data show that N-CaM is more strongly stabilized by P454 than by the H-helix peptide as suggested from the comparison of the magnitudes in deuterium uptake differences following P454 binding (Figure S10) or H-helix binding (see Figure S4 in ^16^). This is in agreement with the mutational analysis of the P454-derived peptides (see below and Table S3), and the crystal structures of the P458:CaM complex, which show that the C-terminal part of P454 establishes several interactions with the hydrophobic groove of N-CaM (Figure 3A). In summary, these studies establish that P454 is an authentic CaM binder displaying original structural and dynamic features (Figures 2-3, S3 and S7 and Table S3).

**Figure 3.**
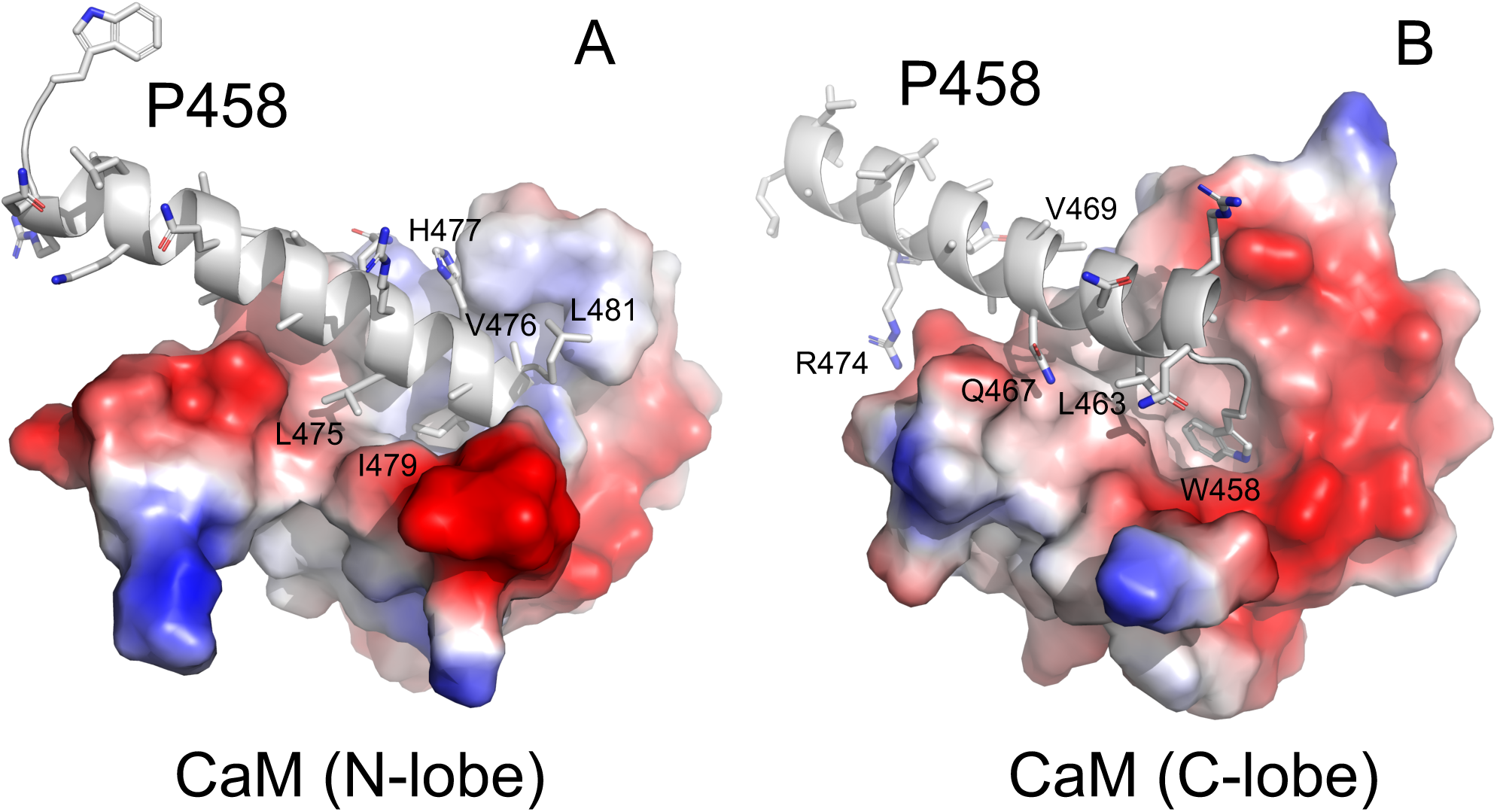
P458:CaM interactions. Close views of the molecular contacts between P458 and the N (panel A) and C (panel B) lobes of CaM (pdb 6YNU). The peptide is shown in cartoon representation and colored in grey. Side chains of key residues interacting with CaM are shown as sticks. These residues establish numerous non-polar interactions, as well a several hydrogen bonds with CaM residues. These contacts are summarized in Table S5B-C.The CaM lobes are represented by their electrostatic surfaces (negative and positive charges are colored in red and blue, respectively).

### 4 Mutations altering P454:CaM and P454:membrane interactions *in vitro*

The structure of the P454:CaM complex presented above reveals several residues of P454 that might be critical for the interaction with CaM, including the aromatic residue W458, the arginine residues R461 (R*1*) and R474 (R*2*), the aliphatic residues L463, L475, I479 and L481, and finally the histidine residue H477. To corroborate the structural data, we designed a series of P454 derivatives in which several of these residues were mutated and characterized both their CaM binding as well as their membrane interaction properties.

The affinity of the P454-derived peptides for CaM was investigated by fluorescence and ITC (Table S3 and Figure S11) through the determination of the dissociation constant K_d_ and the free energy of the peptide:CaM complex (Δ*G*_*Kd*_). Our data shows that the affinity of the P454-derived peptides for CaM is mainly altered by mutations of arginine and aliphatic residues (Table S3 and Figure 4A) with a progressive decrease with the single point mutations L475A, R474Q, H477S, L463A and I479A or with the reduction of the side chain apolarity of the I479 residue (*i*.*e*., I479L, I479V and I479A). These results confirm the contribution of these residues in the complex formation as suggested from the observation of the peptide:CaM crystal structure (Figure 3A).

**Figure 4.**
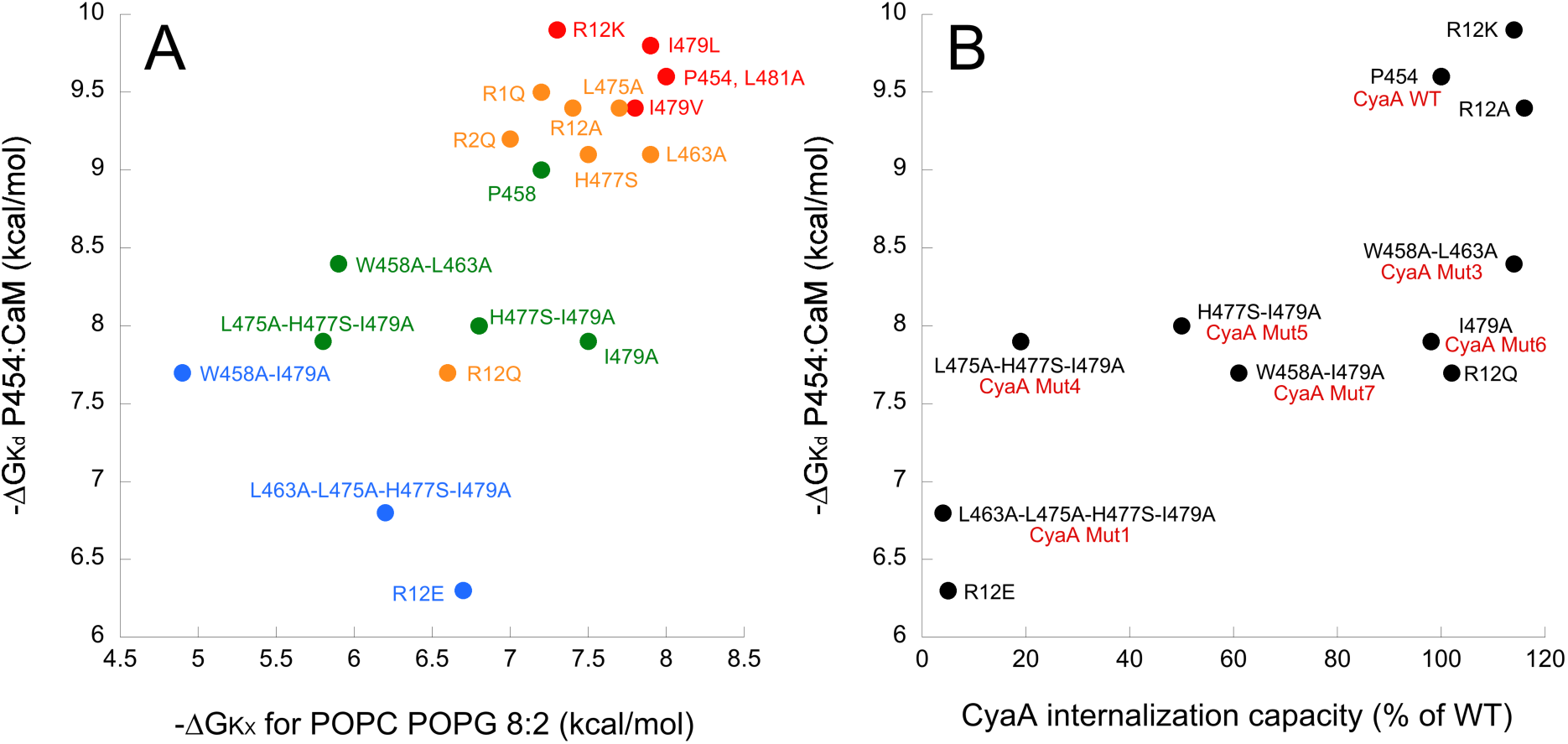
Correlations between *in vitro* properties of P454-derived peptides and the internalization activity of the CyaA recombinant proteins. (**A**) The free energy values of peptide:CaM complex formation (ΔG_Kd_) are plotted as a function of free energy values for peptide solution-to-membrane partitioning (ΔG_Kx_, see Table S3) and permeabilization efficiency (Cp_1/2_ values, see Table S9). The color code refers to the Cp_1/2_ values ranging from red-to-blue (high-to-low permeabilization efficiency, respectively) using a logarithmic scale (red: Cp_1/2_<100 nM, orange: 0.1< Cp_1/2_<1 µM, green: 1< Cp_1/2_<10 µM and blue: 10< Cp_1/2_<100 µM. (**B**) The free energy values of peptide:CaM complex formation (ΔG_Kd_) as a function of relative internalization activity of the CyaA variants (data from Table 1). The peptide names are in black and the names of the recombinant CyaA proteins are in red, if different from the peptide name.

As expected, the affinity of the P454-derived peptides for CaM is affected to a greater extent by multiple mutations. In particular, the double substitution of both arginine residues R461 and R474 (R*12*) into either glutamine or glutamate residues (R*12*Q and R*12*E peptides respectively) resulted in both cases in a significant loss of affinity for CaM. Substitutions of aliphatic and histidine residues at the C-terminus of P454 (H477S-I479A and L475A-H477S-I479A) resulted in a *circa* 20-fold decrease of affinity of P454 for CaM, while the mutation of the N-terminal aromatic and aliphatic residues (W458A-L463A) results in only a 4-fold decrease in P454:CaM affinity (Table S3). These observations are in agreement with the crystal structure of P458:CaM, showing that the residue H477, and more importantly I479, from the C-terminal part of P454 are crucial for P454:CaM interactions (Figure 3A). The W458 substitution also seems to be involved in CaM binding as its combination with I479 (W458A-I479A) induces a significant loss in affinity (Table S3 and Figure 3B). Taken together, the mutational analysis (Table S3), the HDX-MS data (Figure 2F) and the crystal structures of the complex (Figure 3) indicate that the C-terminal part of the peptide strongly interacts with the N-terminal lobe of CaM and, to a lesser extent, the N-terminal part of the peptide interacts with the C-terminal lobe of CaM.

**Table 1.**
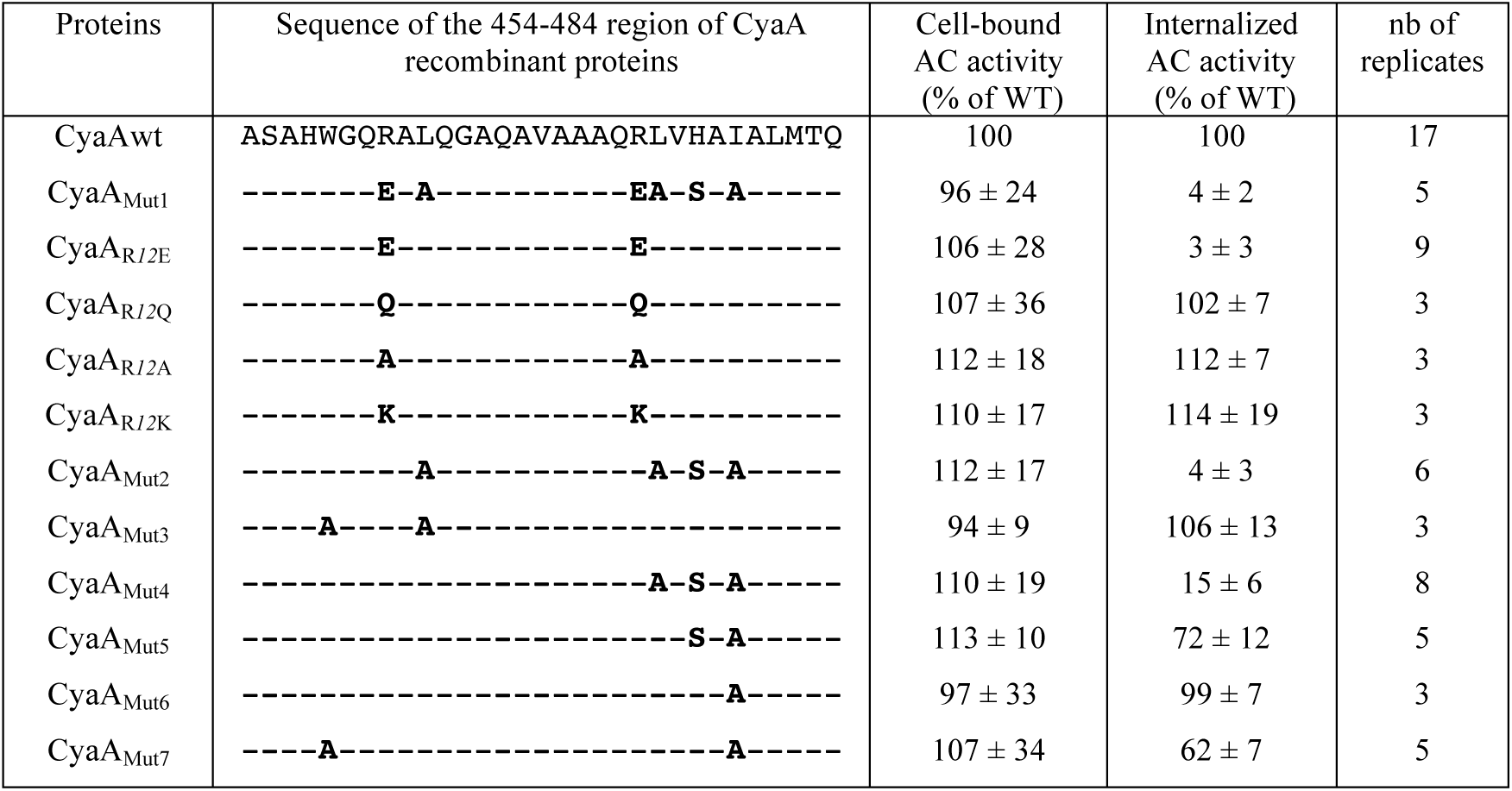
Cytotoxic activities of the recombinant CyaA proteins. The binding and internalization of CyaA and different variants were assayed on sheep erythrocytes as described in the Material and Methods section. The proteins in 6M urea were directly diluted (to a final concentration of 5.6 nM) in sheep erythrocyte suspensions in buffer B. Bound and internalized AC activities for each recombinant proteins is expressed as % of that measured with the wild-type CyaA toxin and represent the average values of at least 3 independent measurements. The substituted residues are indicated in bold letters for each CyaA variants.

We then characterized the membrane binding and permeabilization properties of P454-derived peptides. We analyzed the membrane-induced secondary structure changes of the peptides by circular dichroism (Figure S12 and Table S8) and also determined their partition coefficient, K_X_, and free energy of membrane interaction, Δ*G*_*Kx*_, using fluorescence spectroscopy and ITC (Figure 4A and Table S3). The secondary structure content of the P454-derived peptides is characterized by a disorder-to-helical conformation transition upon membrane binding for most peptides. The Δ*G*_*Kx*_ values indicate that membrane interaction of P454-derived peptides is affected by mutations of arginine, aliphatic and aromatic residues. Notably, a strong correlation is observed between the free energy values of P454-derived peptides for membrane interaction and CaM binding (Figure 4A). We also measured the membrane permeabilization efficiency of the P454-derived peptides, reported as the peptide concentration required for recovering 50% of ANTS fluorescence intensity upon permeabilization (Cp_1/2_ values, see Table S9). We color-coded the Cp_1/2_ values ranging from red-to-blue (high-to-low permeabilization efficiency, respectively) using a logarithmic scale (see legend of Figure 4A for details). As observed for P454 membrane partitioning, the Cp_1/2_ values show a good correlation with the free energy of P454:CaM complex formation (Figure 4A). These results show that the three properties of P454-derived peptides (K_d_, K_X_ and Cp_1/2_) are correlated, *i*.*e*., any mutation affecting one of the parameters will likely affect the other two. Altogether, these data indicate that the biophysical properties of P454 required to interact and destabilize membranes are highly similar to that involved in CaM binding: the peptide must adopt an amphipathic helical conformation with positively charged and apolar residues to exert these three activities *in vitro*.

### 5 Characterization of recombinant CyaA toxins harboring mutations in the P454 region

To explore whether the interaction of the P454 region with membrane and CaM is also involved in CyaA intoxication, we designed several recombinant toxins harboring specific modifications at the key residues identified above (R461 and R474, W458, H477 and aliphatic residues L463, L475 and I479) (Figure S13) and characterized their cytotoxic activities (Table 1). The recombinant toxins were produced in *E. coli* and purified to homogeneity (Figure S14), and their capacities to bind to and translocate their AC domain across the plasma membrane were determined on sheep erythrocytes, a model of target cells (Table 1). In the first recombinant CyaA tested, CyaA_Mut1_, six residues from the P454 region were modified: the arginine residues R461 and R474 were changed to glutamate, the leucine L463 and L475 and the isoleucine I479 to alanine and the histidine residue H477 to serine. These combined modifications did not affect binding of CyaA_Mut1_ to cells but completely abolished translocation of its catalytic domain into the cytosol. This result provides direct evidence that the P454 region is absolutely critical for the invasive activity of the toxin, *i*.*e*., AC translocation into the target cell cytosol.

We then further delineated the respective contribution of these different residues to the cytotoxic properties of CyaA. We first examined the contribution of the arginine residues R461 and R474 to cell intoxication. Charge reversion of these two arginine residues by glutamate residues in CyaA_R*12*E_ was enough to fully abolish the ability of CyaA to invade cells while not affecting toxin binding to plasma membrane (Table 1). However, the substitution of the guanidinium group of the two arginine residues by lysine residues in CyaA_R*12*K_, as well as the neutralization of the arginine residues in CyaA_R*12*Q_ and the deletion of the arginine side chain in CyaA_R*12*A_, did not significantly affect the invasive activities of the corresponding toxins (Table 1). The contribution of the neutral and apolar residues in P454 region to CyaA internalization was evaluated in a second series of recombinant modified toxins, CyaA_Mut2_ to CyaA_Mut7_. The CyaA_Mut2_ variant harboring the 4 mutations L463A, L475A, H477S and I479A, was also completely unable to invade erythrocytes (Table 1). The modified toxin CyaA_Mut4_, which harbors only 3 mutations L475A, H477S and I479A at the C-terminus of P454, was also drastically impaired in translocation capability. The other recombinant toxin that was significantly altered in its invasion activity was CyaA_Mut7_ carrying the double mutation W458A and I479A and showing about 60 % of wild-type translocation activity. Notably, the binding of all these toxins to erythrocytes was not altered, indicating that the specific modifications introduced into the recombinant CyaA proteins do not affect the cell-binding step, but selectively affected the translocation step. Taken together, these results suggest that the ability of the P454 motif to associate with CaM within the target cell is critical for the efficient translocation of the AC domain across the membrane. This is further supported by the correlation between the cytotoxic activity of the CyaA recombinant proteins (data from Table 1) and the affinity of the corresponding P454-derived peptides (data from Table S3) shown in Figure 4B.

### 6 Cytosolic CaM is required for efficient CyaA internalization

The above cell intoxication data (Table 1 and Figure 4B) suggest that the ability of the P454 motif to bind CaM inside target cells might contribute to the translocation of the AC domain across membrane. To test this hypothesis, we analyzed the effects of a CaM inhibitor, calmidazolium (CDZ), which exhibits a high affinity (K_I_ about 10-50 nM) for holo-CaM ^77^. For these experiments, erythrocytes were first incubated in the presence of calcium at 4°C in conditions that allow the toxin to bind to the cell membrane but not to translocate across membranes, as originally described by Rogel and Hanski ^61^. After washing of unbound toxin, a rapid (within minutes) internalization of the catalytic domain was observed upon transfer of the samples to 37 °C (Figure 5). However, when CDZ (10 μM) was added to the cell mixture just prior to the temperature shift from 4 to 37°C, the amount of internalized AC was drastically reduced (Figure 5).

**Figure 5.**
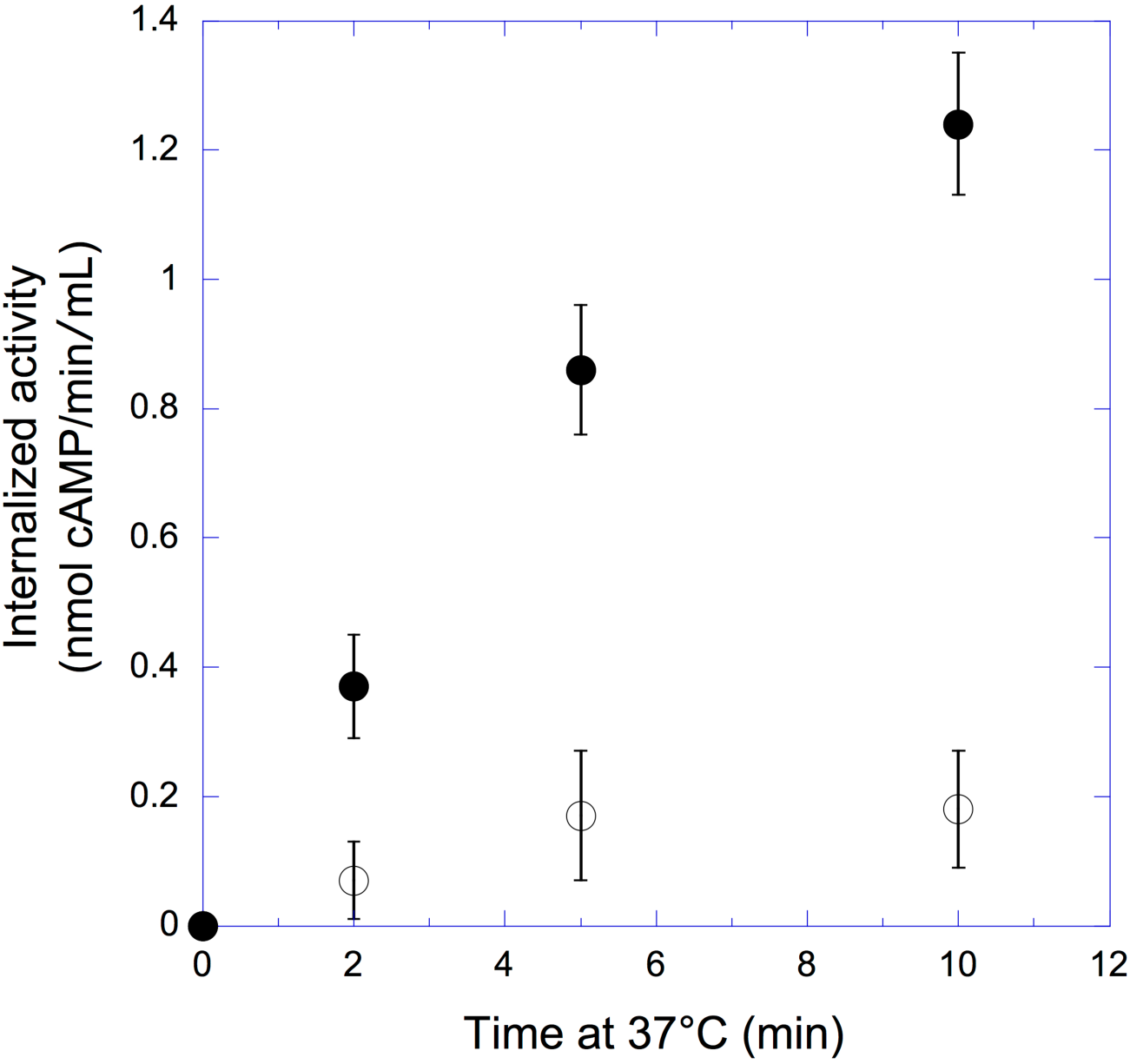
Calmidazolium (CDZ) inhibits CyaA translocation into erythrocytes. Erythrocytes were first incubated with CyaA (5.6 nM) at 4 °C in the presence of CaCl_2_ for 30 min so that the toxin could bind to cells but does not translocate across plasma membrane (see main text). After removal of unbound toxin, 10 μM CDZ was added (open symbols) or not (CyaA, filled symbols) and the cell mixtures were transferred to 37 °C. At the indicated time the cell suspensions were treated with trypsin for 10 min and after addition of soybean trypsin inhibitor, cells were washed and lysed with 0.1% Tween 20 and the internalized AC activity (*i*.*e*., enzyme activity protected from trypsin digestion) was measured as described in Material and Methods.

The inhibitory CDZ concentration (10 μM) is in excess over the total intracellular CaM concentration, estimated to be between 3 to 7 μM ^78-79^ of which up to 10% should be free ^80-81^. In the presence of CDZ, the free CaM in erythrocytes should drop to low nM ranges, *i*.*e*., 1-10 nM (assuming a binding constant for CDZ:CaM of 10-50 nM). This free CaM concentration is enough for binding and activation of AC (Figure S15), given the very high AC:CaM affinity (K_d_ about 0.1 nM) but is well below the affinity constant of P454:CaM (K_d_ about 90 nM). Size exclusion chromatography experiments confirm that CDZ can specifically inhibit CaM association with P454, but not with the AC domain (Figure S16). In agreement, the H:CaM complex (K_d_ about 10 nM, Figure S7 and Table S4) is partially inhibited by CDZ (Figure S16). Notably, it was previously shown that CaM binding to the catalytic domain of CyaA is not required for toxin internalization ^82^. Taken together, we conclude that CDZ blocks CyaA internalization into the cells primarily by preventing CaM binding to the P454 motif. We propose that trapping of the CyaA polypeptide chain by cytosolic CaM *via* the P454 segment may facilitate the entry of the N-terminal AC domain into the cells.

## Discussion

We demonstrate here the critical role of the P454 peptide segment (residues 454 to 484) in the translocation of the catalytic domain of CyaA across the plasma membrane of target cells, and suggest a new mechanism for CyaA invasion. We have previously shown that a deletion of the whole translocation domain TD (residues 373 to 485) hindered the entry of CyaA into target cells ^17^. In the present work, we show that substitutions of a few residues within the P454 segment are sufficient to fully abrogate the delivery of the AC domain into the cell cytoplasm, without impairing toxin binding to target cells. Most importantly, the mutated residues that result in inactivation of CyaA translocation are also involved into two key properties of the P454 motif: firstly, its ability to bind to membranes and destabilize the lipid bilayer, as reported previously ^63, 65^ and corroborated here using various P454-derived peptides; and secondly, its ability to bind with high affinity to holo-CaM, as shown here for the first time. We propose that these two properties are directly implicated in the process of CyaA translocation across the plasma membrane of eukaryotic target cells.

The P454 peptide exhibits characteristics found in many calmodulin-binding peptides that form amphiphilic helices upon association with CaM. We have shown by a combination of biophysical approaches that P454 forms a 1:1 complex with CaM in a calcium-dependent manner, with a K_d_ of about 90 nM at 25°C (Figures S2-S5 and Tables S1-S3). We solved the crystal structures of P454 (actually a slightly shorter peptide, P458, exhibiting similar properties) in complex with holo-CaM, and obtained structural models in solution by SEC-SAXS measurements, which revealed the dynamics and the overall shape of the P454:CaM complex (Figures 2, 3 and S9). Together with HDX-MS data, these studies have revealed the original mode of association of P454 with CaM, which primarily occurs via several interactions between the C-terminal moiety of the peptide and the hydrophobic groove of N-terminal domain of CaM. Comparison of the structural and dynamic characteristics of the P454:CaM complex with those of CaM with MLCK peptide or H peptide (the main CaM-binding site of the AC domain) illustrates the remarkable plasticity of CaM in binding to target proteins. These structural data also indicated key residues of P454 that could be potentially important in CaM-binding (Figure 3). The implication of these amino-acids in CaM was experimentally confirmed by analysis of P454-derived peptides with modified residues (Figure 4). In particular, modification of the two positively charged Arg residues into Glu resulted in a drastic decrease in CaM-binding affinity. Multiple mutations of aliphatic residues L463, L475, I479 to alanine, and histidine residue H477 to serine, also lead to a significant decrease in CaM-binding affinity.

Combining these mutations differently affected binding to CaM, as well as the membrane itself (Figure 4A). In all cases, the affinity values of P454 and P454-derived peptides for CaM were 2-3 orders of magnitude higher than that for membranes, i.e., with dissociation constants in the nM range and µM ranges, respectively (Table S3). Consequently, P454 preferentially interacts with calmodulin over membranes. Moreover, we demonstrate by using a droplet interface bilayers (DIB) technique that P454 can translocate across a lipid bilayer and that holo-CaM favors its accumulation in the *trans* compartment (Figure 1). In contrast, the passage of the fluorescent-P454_R*12*E_ peptide could not be detected unless very high CaM concentrations were loaded in *trans* droplets to overcome the low affinity of this peptide for CaM (K_d_^CaM^ ≈ 20 µM). Taken together, these data suggest that the P454 peptide is capable of translocating across lipid membranes. Moreover, once translocated, the peptide is able to form a complex with CaM. This interaction displaces the P454 peptide from the membrane to form a peptide:CaM complex in solution due to a favorable free energy difference (Figures S4 and S8). The C-terminal part of P454 strongly interacts with the N-CaM lobe via the arginine and apolar residues of the peptide (Figure 3).

The cell intoxication data (Table 1) indicate that the ability of CyaA to translocate its catalytic domain across the membrane of target cells is strongly correlated with both membrane permeabilization efficiency of the P454 motif and its affinity for CaM (Figure 4). We propose that after binding of CyaA to target cells, the translocation region interacts with the plasma membrane and the P454 motif locally destabilizes it, leading to calcium influx ^46^. The negative potential across the membrane may then favor the passage of the positively charged P454 region through the lipid bilayer to reach the cytoplasmic side of the membrane where it can associate with holo-CaM. Binding to a cytosolic partner would strongly increase the entropic pulling force by preventing the backward movement of the polypeptide chain across the plasma membrane. This strong interaction between the P454 motif and CaM in the cytosol may then favor the translocation of the catalytic domain across the plasma membrane. Collectively, we propose that the membrane destabilization caused by P454, followed by its translocation and binding to calmodulin, are essential to convert the stochastic process of protein translocation into an efficient vectorial chain transfer into the cytoplasm of host cells. To evaluate the contribution of P454:CaM complex formation to the AC translocation process into cells, we analyzed the effects of a CaM inhibitor, calmidazolium (CDZ), which exhibits a high affinity for calmodulin ^77^.

We found that CDZ selectively blocks AC internalization into cells and we provide evidence that CDZ primarily interferes with CaM binding to the P454 motif of CyaA, without altering CaM association with the catalytic domain and activation of its enzymatic activity (Figures 5, S15 and S16).

In summary, our results indicate that CaM plays a dual role in CyaA intoxication: firstly, it acts as a cytosolic binder that can grab the polypeptide chain by the P454 motif to favor the translocation of the catalytic domain across the plasma membrane of target cells; secondly, after entry of the catalytic domain into the cell, CaM can associate with the AC domain to activate its enzymatic activity by stabilizing active conformations of the catalytic site ^15-16^. This illustrates a remarkable parsimony in the molecular mechanism of the CyaA toxin which exploits the same ubiquitous and abundant protein, CaM, to enable two essential functions, entry and activation, that are both required for an effective intoxication of eukaryotic cells. Interestingly, we demonstrated in a prior study that CyaA can efficiently translocate across a biomimetic membrane model made of a tethered lipid bilayer (tBLM) assembled over an amine-gold surface derivatized with CaM ^24^. Remarkably, in this synthetic system, CaM is the only eukaryotic component needed for translocation of the CyaA catalytic domain. This observation nicely fits with the present results indicating that CaM may play the dual function of both cytosolic binder and activator of CyaA.

Interestingly, several toxins hijack eukaryotic cytosolic factors to achieve the translocation of their respective catalytic domains: these toxins contain segments able to translocate across target membranes and to interact with host soluble proteins ^83-97^. We propose that the formation of such toxin:target complexes significantly contributes to the energy required to achieve the translocation of bacterial toxin catalytic domains across membranes of eukaryotic cells.

## Supplementary Information

the file contains the materials and methods section, Tables S1 to S9, Figures S1 to S16, and the supplementary references.

## Acknowledgments

A.V. was supported by a DIM MalInf grant from the region Ile-de-France. M.S. was supported by the Pasteur - Paris University (PPU) International PhD Program. D.P.O.B. was supported by Institut Pasteur (grants PasteurInnoV15006-01A and PTR451). P.G. was supported by Sorbonne Université. We acknowledge SOLEIL and ESRF for provision of synchrotron radiation facilities. We thank the staff of the SWING beamline for constant support and help during SAXS data collection, the staff of the PROXIMA-1 (Synchrotron SOLEIL, St Aubin, France) and MASSIF (Synchrotron ESRF, Grenoble, France) beamlines for assistance during the X-ray diffraction data collection.

## Data Availability

All relevant HDX-MS, X-ray and SAXS data are available in supporting information. The crystal structures have been deposited on the PDB with the access codes 6YNU and 6YNS. The molecular model and experimental SAXS data have been deposited on SASBDB (Small Angle Scattering Biological Data Bank, http://www.sasbdb.org/aboutSASBDB/) under the SAS code SASDJ64 (calcium-bound calmodulin) and SASDJ74 (P454 peptide from *B. pertussis* CyaA toxin complexed with calmodulin).

## Abbreviations

AC: adenylate cyclase catalytic domain
CaM: calmodulin
C-CaM,: C-terminal domain of CaM
CyaA: adenylate cyclase toxin
HDX-MS: hydrogen/deuterium exchange mass spectrometry
IDR: intrinsically disordered region
MEMHDX: Mixed-Effects Model for HDX experiments
MLCK: myosin light chain kinase
MS: mass spectrometry
N-CaM: N-terminal domain of CaM
pdb: Protein Data Bank
SASBDB: Small Angle Scattering Biological Data Bank
SAXS: small-angle X-ray scattering
SEC: size exclusion chromatography.

