## Supplemental Information for "A high-affinity calmodulin-binding site in the CyaA toxin translocation domain is essential for invasion into eukaryotic cells"

The Supplementary information file contains the materials and methods section, Tables S1 to S9, Figures S1 to S16, and the supplementary references.

### Materials

1-palmitoyl-2-oleoyl-sn-glycero-3-phosphocholine (POPC, reference 850457C), 1-palmitoyl-2-oleoyl-sn-glycero-3-[phospho-rac-(1-glycerol)] (POPG, reference 840457C) were purchased from Avanti Polar Lipids (Alabaster, AL, USA). ANTS (A-350, 8-aminonapthalene-1,3,6 trisulfonic acid), and DPX (X-1525, p-xylene-bis-pyridinium bromide) were purchased from Molecular Probes (Eugene, OR, USA). Squalene, TPCK-treated trypsin, and soybean trypsin inhibitor were from Sigma-Aldrich. Calmidazolium chloride was purchased from Calbiochem.

### Buffers used in this study

Experiments were performed using these 5 buffers:

Buffer A: 20 mM HEPES, 150 mM NaCl pH 7.4

Buffer B: 20 mM HEPES, 150 mM NaCl, 2 mM $CaCl_2$ pH 7.4

Buffer C: 20 mM HEPES, 150 mM NaCl, 4 mM $CaCl_2$ pH 7.4

Buffer D: 20 mM HEPES, 150 mM NaCl, 8 mM $MgCl_2$, 2 mM $CaCl_2$, pH 7.4

Buffer E: 20 mM HEPES, 150 mM NaCl, 9.8 mM $MgCl_{2,}$ 0.2 mM $CaCl_2$, pH 7.4

### Peptides synthesis

The P454 peptide corresponds to residues 454 to 484 of the CyaA toxin (ASAHWGQRALQGAQAVAAAQRLVHAIALMTQ) and contains a single native tryptophan W458. The P454-derived peptides were produced and purified by Genosphere Biotechnologies (Paris, France). All peptides were capped on the N-terminus with an acetyl group and on the C-terminus with an amide group, except for the P458 peptide (uncapped N-terminus). Their molecular weight was determined by MALDI-TOF and purity analyzed by HPLC.

The sequences of P454-derived peptides are as follows:

- P454 L463A corresponds to the substitution of L463 by an alanine: ASAHWGQRA**A**QGAQAVAAAQRLVHAIALMTQ.
- P454 L475A corresponds to the substitution of L475 by an alanine: ASAHWGQRALQGAQAVAAAQR**A**VHAIALMTQ.
- P454 L481A corresponds to the substitution of L481 by an alanine:

ASAHWGQRALQGAQAVAAAQRLVHAIA**A**MTQ.

- P454 I479L, I479V and I479A correspond to the substitution of I479 by a leucine, a valine or an alanine, respectively:

ASAHWGQRALQGAQAVAAAQRLVHA**L**ALMTQ

ASAHWGQRALQGAQAVAAAQRLVHA**V**ALMTQ

ASAHWGQRALQGAQAVAAAQRLVHA**A**ALMTQ.

- P454 H477S corresponds to the substitution of H477 by a serine:

ASAHWGQRALQGAQAVAAAQRLV**S**AIALMTQ.

- P454 H477S-I479A corresponds to the substitution of H477 by a serine and I479 by an alanine: ASAHWGQRALQGAQAVAAAQRLV**S**A**A**ALMTQ.
- P454 W458A-I479A corresponds to the substitution of both W458 and I479 by an alanine: ASAH**A**GQRALQGAQAVAAAQRLVHA**A**ALMTQ.
- P458 peptide corresponds to a shorter P454 peptide, from residue 458 to 481:

WGQRALQGAQAVAAAQRLVHAIAL.

- P454 L463A-L475A-H477S-I479A corresponds to the substitution of L463, L475 and I479 by an alanine and H477 by a serine:

ASAHWGQRA**A**QGAQAVAAAQR**A**V**S**A**A**ALMTQ.

- P454 W458A-L463A corresponds to the substitution of both W458 and L463 by an alanine:

ASAH**A**GQRA**A**QGAQAVAAAQRLVHAIALMTQ.

- P454 L475A-H477S-I479A corresponds to the substitution of both L475 and I479 by an alanine and H477 by a serine:

ASAHWGQRALQGAQAVAAAQR**A**V**S**A**A**ALMTQ.

The two arginine residues R461 (referred as $R_{1}$) or R474 (referred as $R_{2}$), or both of them (referred as $R_{12}$) were substituted by glutamine ($R_{1}$Q, $R_{2}$Q and $R_{12}$Q), alanine ($R_{12}$A), lysine ($R_{12}$K) or glutamate ($R_{12}$E) residues [1], yielding the following sequences:

P454 $R_{1}$Q: ASAHWGQ**Q**ALQGAQAVAAAQRLVHAIALMTQ

P454 $R_{2}$Q: ASAHWGQRALQGAQAVAAAQ**Q**LVHAIALMTQ

P454 $R_{12}$Q: ASAHWGQ**Q**ALQGAQAVAAAQ**Q**LVHAIALMTQ

P454 $R_{12}$A: ASAHWGQ**A**ALQGAQAVAAAQ**A**LVHAIALMTQ

P454 $R_{12}$K: ASAHWGQ**K**ALQGAQAVAAAQ**K**LVHAIALMTQ

P454 $R_{12}$E: ASAHWGQ**E**ALQGAQAVAAAQ**E**LVHAIALMTQ

The P454, P454 $R_{12}$E and H-helix (corresponding to residues 233-254 of CyaA) peptides were also synthetized with a 5-carboxytetramethylrhodamine (TAMRA) linked to their N-terminus and capped at the C-terminus with an amide group for the droplet interface bilayer experiments.

**Production and purification of proteins used in this study**

Human Calmodulin (CaM)

CaM was produced in *E. coli* and purified as previously described [2-4]. Briefly, CaM was precipitated with ammonium sulfate followed by a glacial acetic acid precipitation. Then, CaM was purified as follows: a first HIC on Phenyl Sepharose (EDTA-CaM), an IEC on Q-Sepharose fast flow, a second HIC on Phenyl Sepharose (calcium-CaM) and a SEC on Sephacryl S100. Protein concentration was determined by spectrophotometry ($\varepsilon_{280nm}$ = 2,980 $M^{-1}$ $cm^{-1}$).

CyaA recombinant proteins

All the CyaA recombinant proteins were produced in *E. coli* BLR. The *E. coli* strain XL1-Blue (Stratagene, California) was used for DNA manipulation and *in vitro* DNA manipulations were performed according to standard protocols [5].

The expression vector used for the wild-type and modified CyaA proteins is derived from plasmid pT7CACT1 ([6], kind gift from Peter Sebo). In a first step, we constructed pCACTw11 in which the CyaA coding region located between the unique BstBI and NcoI sites of pT7CACT1 was replaced, with the Gibson technique [7], using the NEBuilder® HiFi DNA Assembly Cloning Kit from New England Biolabs, USA) by a synthetic DNA fragment (DLTw11, Figure S13, obtained from Twist Bioscience, USA) encoding the native CyaA sequence with *E. coli* optimized codons and harboring several unique restriction sites flanking the P454 region (Figure S13). Plasmid pCACTw11 expresses the native, acylated, and fully functional CyaA toxin. Plasmid pCACTw16, that expresses the modified protein $CyaA_{Mut1}$, was constructed similarly by HiFi subcloning between the BstBI and NcoI sites of a synthetic DNA fragment (DLTw16, Figure S13) encoding a mutated CyaA sequence with six modified residues in the P454 region (R461E, L463A, R474E, L475A, H477S and I479A). Plasmids encoding the CyaA mutants $CyaA_{R12E}$, $CyaA_{R12Q}$, $CyaA_{R12A}$, $CyaA_{R12K}$, $CyaA_{Mut6}$, and $CyaA_{Mut7}$, were constructed by HiFi subcloning between the BstBI and HindIII sites of pCACTw11 (or pCACTw16) of synthetic DNA fragments (obtained from Twist Bioscience) encoding the CyaA sequences with appropriate codon changes. Plasmids encoding the CyaA mutants $CyaA_{Mut2}$, $CyaA_{Mut3}$, $CyaA_{Mut4}$, and $CyaA_{Mut5}$ were constructed by HiFi subcloning between the BstBI and HindIII sites of pCACTw11 (or pCACTw16) of PCR-amplified fragments with synthetic oligonucleotides (obtained from Sigma-Aldrich) designed to introduce the appropriate codon changes. All DNA sequences of recombinant plasmids were verified by DNA sequencing (performed by Eurofins, France). Further details of plasmid construction and sequence can be provided upon request.

The wild-type and modified recombinant CyaA proteins were produced and purified as previously described [8-10]. Briefly, inclusion bodies were resuspended overnight in 8 M urea, 20 mM HEPES. Proteins were purified on two consecutive IEC (Q-Sepharose fast flow and Q-Sepharose Hi-Performance) equilibrated with 6 M urea, 20 mM HEPES, 150 mM NaCl, pH 7.4. Proteins were eluted with a NaCl gradient. Proteins were further purified onto a HIC Phenyl-Sepharose column. Proteins were eluted with 6 M urea, 20 mM HEPES, pH 7.4. Finally, proteins were loaded on a Sephacryl S500 equilibrated with 6 M urea, 20 mM HEPES, 50 mM NaCl, pH 7.4. Proteins were concentrated on Amicon 50 kDa. Protein concentration was determined by spectrophotometry ($\varepsilon_{280}$ = 144,000 $M^{-1}$ $cm^{-1}$). The primary sequence of the WT CyaA is displayed in Figure S1 and the sequences with the substituted residues in the P454 region of the CyaA mutants are listed in Table 1.

**Lipid vesicles preparation**

Small unilamellar vesicles (SUVs) and large unilamellar vesicles (LUVs) were prepared with POPC and POPG at a molar ratio of 8:2, at a lipid concentration of 10 mM in buffer A, as previously described [1]. Briefly, SUVs and LUVs were prepared by reverse phase evaporation and filtered through polycarbonate filters of 1 µm and 0.2 µm of pore size diameter to obtain LUV. SUVs were further sonicated. The hydrodynamic diameters and mean net charge of the vesicles were measured by dynamic light scattering (DLS) and electrophoretic mobility, respectively, using a ZetaSizer NanoZS (Malvern Pananalytical, Orsay, France) [11].

**Peptide titrations by SUVs and CaM followed by tryptophan fluorescence**

Titrations were carried out with a FP-8200 Jasco spectrofluorimeter, equipped with a Peltier-thermostated ETC-272T at 25°C. A bandwidth of 5 nm was used for both excitation and emission beams. Fluorescence experiments were carried out in a 109.004F-QS cuvette (Hellma, France) with constant stirring. Fluorescence emission was recorded between 300 and

400 nm at a scan rate of 100 nm/min, with an excitation wavelength of 280 nm. Fluorescence emission spectra were corrected for SUV or CaM baselines. Fluorescence intensity ratio at 330 nm over 370 nm was used to measure the partition coefficient ($K_X$) of peptides between the solution and membranes [1, 12] and the affinity ($K_d$) of peptides for CaM [13].

#### Determination of partition coefficient $K_X$ and dissociation constant $K_d$ from experimental data

The partition coefficient $K_X$ is defined as the ratio of protein in the lipid ($P_L$) and in buffer ($P_W$) phases [12, 14], expressed by: $K_X = \frac{[P_L]}{[L]} \Big/ \frac{[P_W]}{[W]}$, giving $\frac{[P_W]}{[P_L]} = \frac{[W]}{K_X[L]}$, where [W] the concentration of water (55.5 M) and [L] the lipid concentration.

The fraction of peptide partitioned into the membrane, $f_{PL}$, is equal to:

$f_{P_L} = \frac{[P_L]}{[P_T]} = \frac{[P_L]}{[P_L]+[P_W]} = \frac{1}{1+(\frac{[P_W]}{[P_L]})} = \frac{1}{1+(\frac{[W]}{[L]K_X})}$, where $P_T$ is the total concentration of peptides $[P_T] = [P_L] + [P_W]$.

The dissociation constant, $K_d$, is obtained from titration experiments of the P454-derived peptides by CaM and is expressed as follows: $K_d = \frac{[P]*[CaM]}{[P{:}CaM_n]}$, given $\frac{K_d}{[CaM]} = \frac{[P]}{[P{:}CaM]}$ and the fraction of complex, $f_{P:CaM}$, is: $f_{P:CaM} = \frac{[P{:}CaM]}{[P{:}CaM]+[P]} = \frac{1}{1+\frac{[P]}{[P{:}CaM]}} = \frac{1}{1+\frac{K_d}{[CaM]}}$.

The equations are fitted to the experimental data using the KaleidaGraph software, providing the $K_X$ and $K_d$ values. The partition coefficient is related to the apparent dissociation constant: $K_X \times K_d = [W]$ and to the free energy of partition $\Delta G_{Kx} = -RTln(K_X)$, where ΔG is the free energy in kcal/mol, R the gas constant ($R = 1.98 \times 10^{-3}$ kcal/mol/K) and T the temperature in Kelvin. The free energy of peptide CaM interaction, $\Delta G_{Kd}$, is given by $\Delta G_{Kd} = RTln(K_d)$.

#### Sedimentation velocity-Analytical ultracentrifugation (SV-AUC)

SV-AUC experiments were carried out in a ProteomeLab XL-I analytical ultracentrifuge (Beckman Coulter, Brea, CA, USA). Samples of 400 µL and 410 µL of buffer were loaded in cells equipped with 12-mm thick double-sector epoxy centerpieces and sapphire windows. P454 and CaM were tested alone (at 57 µM and 72 µM, respectively) or mixed in a molar ratio 1:1 (at 70 µM each). Samples were incubated 2h at 20°C in the ultracentrifuge and, then span at 42,000 rpm in an An-50 Ti rotor. Detection of protein concentration as a function of radial position and time was monitored with the ProteomeLab software (Beckman Coulter,

Brea, CA, USA) by both absorbance at 280 nm and interferometry. Four hundred scans were collected at 3-min intervals with a radial step size of 0.003 cm. SV scans were analyzed using the continuous size distribution model c(s) of the software Sedfit [15]. All the c(s) distributions were calculated with a fitted frictional ratio f/f0 and a maximum entropy regularization procedure with a confidence level of 0.68. All partial specific volumes were theoretically calculated with the software Sednterp (Spin Analytical, Berwick, ME, USA). Buffer density and viscosity at 20°C of 1.0310 $g.mL^{-1}$ and 1.0061 cP respectively were also determined with Sednterp for buffer B.

**Isothermal titration calorimetry**

ITC experiments were performed using a VP-ITC calorimeter (Malvern Panalytical, Orsay, France). The ITC experiments were performed in buffer B or buffer A complemented with 2 mM EDTA. For a typical titration, the solution of analyte (ranging from 0.2 to 10 µM) was loaded in the reaction cell. The titrants (ranging from 4 to 430 µM) were loaded into the syringe and then the titrant was injected into the reaction cell, usually at intervals ranging from 200 to 600 seconds. Heats of dilutions were measured by injecting the titrant into the protein-free buffer, and were subtracted from the heat of reaction. To obtain the thermodynamic parameters ΔH, ΔS and ΔG, titration profiles were analyzed using the single- or two-site models with Origin7 software (OriginLab). Changes of heat capacity (ΔCp) for the P454:CaM complex formation was obtained from the slope of ΔH values versus temperature. ITC experiments were performed at 10, 20, 25, 30 and 38°C.

**Synchrotron Radiation Circular Dichroism**

Synchrotron radiation circular dichroism (SRCD) was performed on the DISCO beamline of the synchrotron SOLEIL (Saint-Aubin, France). Spectra were recorded at 25°C with an integration time of 1.2 s and a bandwidth of 1 nm with a resolution of 1 nm. A far-UV spectrum represents the average of four individual scans. QS cells (Hellma, France) with a pathlength of 20, 50, 100 or 200 µm (depending on final protein concentrations) were used to record spectra in the far-UV range (from 190 to 250 nm). CD spectra of peptides and proteins were measured in buffers A and B, 20% of trifluoroethanol (TFE), in the presence of SUVs (POPC:POPG 8:2) or CaM. Secondary structure content was estimated using Bestsel software [16-17].

**Membrane permeabilization**

To monitor membrane permeabilization induced by peptides, ANTS (fluorophore probe) and DPX (quencher) were encapsulated into large unilamellar vesicles (LUVs). The LUVs were prepared at a concentration of 10 mM lipid at a POPC:POPG molar ratio of 8:2 containing 20 mM ANTS and 60 mM DPX. The multilamellar vesicle suspension was extruded through 0.4- and 0.2 μm polycarbonate filters to produce LUVs. The unencapsulated ANTS and DPX were removed by gel filtration with a 5 mL Sephadex G-25 column (GE Healthcare Life Sciences). For permeabilization assays, LUVs were incubated in buffer A at 0.45 mM lipid concentration at 25°C in a 101-QS cuvette (Hellma, France) and under constant stirring. The excitation wavelength was set at 390 nm and the emission of ANTS was continuously measured at 515 nm. The maximum intensity of permeabilization, corresponding to the maximum recovery of ANTS fluorescence, was measured after addition of 0.1% of Triton X100. The concentration of membrane-bound peptides required to induce the recovery of half intensity of the ANTS fluorescence ($C_{P1/2}$) was extracted from the permeabilization intensity as a function of peptide concentration [1].

**Membrane permeabilization with CaM and EDTA**

LUVs were incubated in buffer B at a lipid concentration of 0.45 mM. P454 was added at a concentration inducing vesicle permeabilization. After a few seconds of permeabilization, holo-CaM was added to the solution at a peptide:CaM molar ratio of 1:3. EDTA was subsequently added in excess to chelate calcium and convert holo-CaM to apo-CaM.

**Cell intoxication assays**

Toxin binding and translocation into sheep erythrocytes were assayed essentially as described previously [18]. The CyaA proteins were diluted from the urea stock solutions to a final concentration of 5.6 nM (1 μg/mL) into sheep erythrocytes (5 x $10^8$ cells/mL, from Charles River Laboratories, Wilmington, MA, USA) in buffer A or in buffer B complemented with 2 mM EDTA, and incubated at 30°C for 20 min. An aliquot was taken to measure the total adenylate cyclase (AC) activity that (measured in the presence of 1 μgM CaM at 30°C and pH 8.0, as described in [18]). The cell suspensions were chilled on ice and centrifuged at 4°C. The pelleted cells were resuspended in buffer A and separated into two different batches. One was centrifuged again and pelleted cells were lysed with 0.1% Tween 20. The measured enzymatic activity corresponds to the toxin bound to the cells and was expressed as a percentage of total activity added to the cells. The second batch was digested with 20 μg of

TPCK-treated trypsin (Millipore Sigma, Burlington, MA, USA) for 10 min at 30°C to digest the adenylate cyclase that remained at the external surface of the erythrocytes. A 5-fold excess of soybean trypsin inhibitor was added and cells were washed with buffer A and lysed with 0.1% Tween 20. The activity protected from trypsin digestion corresponds to the internalized toxin. Activities are expressed as percentages of total toxin added to the erythrocytes (taken as 100%) and represent average values from at least three independent measurements. One unit of CyaA activity corresponds to 1 μmol of cAMP formed per min at 30°C and pH 8.0. the bound and internalized AC activities measured with the CyaA recombinant proteins were normalized to that of the WT CyaA toxin.

For the CyaA intoxication experiments in the presence of calmidazolium (CDZ), in order to minimize the cell exposure to this potent CaM inhibitor, we performed a two-step intoxication assay as described originally by Rogel and Hanski [19]. For this, the erythrocytes (5 x $10^8$ cells/mL) were in a first step incubated with CyaA (1 μg/mL, 5.6 nM) in buffer A plus 2 mM $CaCl_2$ at 4 °C for 30 min. In these conditions, the toxin can bind to the cell membrane but cannot translocate across membrane [19]. The unbound toxin was removed by extensive washing in buffer A at 4 °C. An aliquot was kept in ice for measuring bound activity (that amounted to about 0.8 to 1.5% ot total CyaA activity, 5 separate experiments). The cells were then resuspended in cold buffer A plus 2 mM $CaCl_2$ and split in two fractions. CDZ (10 μM) was added to one fraction that was further incubated at 4 ° C for 1-2 min to allow diffusion and cell penetration of the drug, and then both fractions were transferred to 37 °C. At 2.5 and 10 min, aliquots of cell suspension were withdrawn and mixed with trypsin as above for 10 min at 30°C. After addition of soybean trypsin inhibitor, the cells were washed with buffer A then lysed with 0.1% Tween 20 and the internalized AC activity protected from trypsin digestion was measured and expressed as a percentage of bound AC activity. Importantly, CDZ (10 μM) has no inhibitory effect on the enzymatic activity when assayed in the presence of 1 mM CaM (see Figure S15).

**Size exclusion chromatography of P454, H and AC364 with CaM in the presence of CDZ**

The SEC experiments were performed on a TSKgel G3000SW 7.5 mm ID * 30 cm - column equilibrated in buffer B at a flow rate of 0.6 mL/min at RT. The elution profiles were measured by absorbance at 280 nm, conductivity and by tryptophan fluorescence on a FP750 fluorometer (Ex.: 280 nm; Em.: 340 nm; bandwidths 10/10 nm) equipped with a on-line cell. Tryptophan fluorescence is used for the SEC profiles because this allows monitoring the tryptophan-containing CaM ligands, i.e., P454, H and AC364, while CaM provides only a weak fluorescence signal. CDZ alone loaded at 40 uM in the SEC does not contribute to the fluorescence signal. The samples were prepared at the following final concentrations: 10 µM of peptides (P454 and H) and AC364, 15 µM of CaM and 40 µM of CDZ. The initial concentration of CDZ was 14.5 mM in 100% DMSO. Addition of CDZ at a final concentration of 40 µM gives a final concentration of less than 0.3 % of DMSO in the samples containing CDZ.

**Peptide translocation followed by droplet interface bilayer experiments**

An aqueous phase is prepared with buffer D or E and the oil phase is prepared with 15 % (v/v) of chloroform and 0.08 % (w/v) of lipids (POPC:POPG at a 9:1 molar ratio) in squalene. A population of droplets is obtained by adding 2 µL of the aqueous phase to 20 µL of the oil phase before briefly shaking the mixture to create an invert emulsion. Two droplet populations were prepared: one containing the TAMRA-labeled peptide in buffer D, the "source", *cis*-droplets and the other one containing buffer D or E, or in the presence or in the absence of calmodulin, the "capture", *trans*-droplets. The two droplet populations were deposited successively on a glass slide (covered with a thin layer of polydimethylsiloxane, PDMS), which allows their mixing and the random formation of pairs of droplets. A lipid bilayer is thus formed at the interface between the two adhering droplets. We then track the possible translocation of the TAMRA-labeled peptides through this bilayer. The fluorescence signals of the "source" and the "capture" droplets were monitored by epifluorescence microscopy for 15 minutes with an Olympus microscope and a MicroMax camera (Princeton Instrument). Fluorescent probes were excited by illumination with a mercury lamp with an excitation filter (515-555 nm).

Image analysis

Quantification of the translocation process is possible thanks to the analysis of the fluorescence images of the droplet pairs. Even if fluorescence is not directly proportional to peptide concentration due to photo-bleaching and small volume variations, one can hypothesize that the fluorescence intensities of the "source" and the "capture" droplets are proportional to the ratio of their concentrations. Moreover, as the initial concentration in the "source" droplet is known and as the total amount of peptide is conserved during the experience, we can write that: $C_s(t=0)\times V_s = C_s(t)\times V_s + C_c(t)\times V_c$, where $C_i$ is the concentration of the droplet i, $V_i$ its volume, which was determined from microscopy images (i="s" for the source droplet and "c" for the capture droplet). We can thus estimate $C_c$, the concentration of the "capture" droplet at any time as:

$$C_c = \frac{C_s(t=0)\times V_s}{V_c + \frac{V_s \times F_s}{F_c}}$$

where $F_i$ is the fluorescence intensity measured for droplet i.

For each experiment, background fluorescence was subtracted from the fluorescence of the droplet pair. Furthermore, during experiments we observed that a certain amount of peptide was able to exit the droplets, reaching the oil phase and possibly go back inside another droplet. To take into account this amount of peptide that could go inside the "capture" droplet without crossing the bilayer, we performed all experiments with a control droplet, or "dark" droplet, initially peptide-free and we subtracted its fluorescence from the one of the "capture" droplet of the pair so that the remaining fluorescence increase of the capture droplet was from translocation through the bilayer only and not contaminated by the oil-mediated peptide diffusion phenomenon.

**Small angle X-ray scattering (SAXS)**

Data acquisition

X-ray scattering data were collected at the SWING beamline of the SOLEIL Synchrotron (Saint-Aubin, France). All measurements were performed using a size-exclusion HPLC column on-line with the SAXS measuring cell, a 1.5 mm diameter quartz capillary contained in an evacuated vessel [20]. All experimental details are given in Table S5 in accordance with the guidelines given in [21].

We studied solutions of CaM in the presence of a 2-fold excess of P454. All proteins were analyzed in buffer C. Scattering of the elution buffer before void volume was recorded and used as buffer scattering for subtraction from all protein patterns. For each frame, the protein concentration (about 0.6/2 mg/mL at the top of elution peak for CaM-P454 and CaM respectively) was estimated from UV absorption at 280 nm using a spectrometer located immediately upstream of the SAXS measuring cell. We also studied the P458:CaM complex in solution. Due to low sample concentration, the recorded SAXS data were of much lower quality, sufficient however to appear essentially identical to that of the P454:CaM complex.

Data analysis

SAXS data were normalized to the intensity of the incident beam and background subtracted using the programs FoxTrot (courtesy of SWING beamline) and Primus (https://www.embl-hamburg.de/biosaxs/primus.html) [22]. Identical frames under the main elution peak were selected using Cormap [23] and averaged for further analysis. The radius of gyration ($R_g$) was evaluated using the Guinier approximation [24] and also derived from the distance distribution function P(r) calculated by the program GNOM [25]. Two independent molecular masses M estimates were obtained using the SAXSMoW2 and ScÅtter programs available at the URLs http://saxs.ifsc.usp.br/ and https://bl1231.als.lbl.gov/scatter/, respectively. Both mass determinations do not depend on the value of the protein concentration c, a feature of particular interest in the case of CaM containing samples since CaM does not possess any tryptophan.

*Ab initio* shape determination

The shape or envelope of the complex of CaM with P454 peptide was determined using the *ab initio* program DAMMIF [26]. 20 runs yielded as many shapes that were compared using the DAMAVER suite of routines based on the calculation of the Normalized Spatial Discrepancy (NSD) that is lower than 0.7 if all 20 models are similar. The final envelope is obtained using DAMMIN [27] that uses the envelope of all superimposed DAMMIF models (the union of all models) as initial volume.

Rigid-body modeling

SAXS models of the P454:CaM complex were obtained from the crystal structures by using the DADIMODO modeling program [28] that uses an all-atom description of the molecule and explores conformational space by random modifications of the internal degrees of freedom $\phi$ and $\psi$ angles within the helix between the two domains of CaM.

The molecular model and experimental SAXS data have been deposited on SASBDB (Small Angle Scattering Biological Data Bank, http://www.sasbdb.org/aboutSASBDB/) under the SAS code SASDJ64 (calcium-bound calmodulin) and SASDJ74 (P454 peptide from *B. pertussis* CyaA toxin complexed with calmodulin).

**Crystallography**

Crystallization and diffraction data collection

To obtain crystals suitable for X-ray diffraction studies, we used the procedure and the crystallization pipeline implemented in the medium-scale crystallography platform at the Institut Pasteur, Paris [29]. Briefly, crystallization screening trials of calmodulin in complex with different peptides (molar ratio 1:2) were carried out by the sitting drop vapor-diffusion method with a Mosquito automated nanoliter dispensing system (TTP Labtech, Melbourn, UK) at 291K. Sitting drops of 400 nL were set up in Greiner plates for 672 commercially available screening solutions using a 1:1 mixture of protein sample and reservoir well solution (150 µL). The plates were stored in a RockImager (Formulatrix, Bedford, USA) automated imaging system to monitor crystal growth. Manual optimizations of the different obtained hits were performed in Linbro plates with the hanging-drop method by mixing 2 µl of protein with 2 µl of reservoir solution. The best crystals were obtained with the conditions shown in Table S5A. For data collection, the crystals were flash-cooled in liquid nitrogen using a condition of crystallization supplemented by 30% (v/v) of glycerol as a cryoprotectant. The X-ray diffraction data were collected on beamline PROXIMA-1 (Synchrotron SOLEIL, St Aubin, France) or on beamline MASIFF (Synchrotron ESRF, Grenoble, France) and processed with autoPROC (Global Phasing Ltd.) [30].

Structure determination and model refinement

The crystal structures of the calmodulin in complex with the synthetic P458 peptide were solved by the molecular replacement technique using separately, the N-ter and C-ter lobes of calmodulin (from the PDB entry 1CTR) as search models with program Phaser [31]. Peptides were manually traced into Fourier difference electron density maps. Final models of the complexes were obtained through interactive cycles of manual model building with Coot [32] and reciprocal space refinement with Buster [33]. X-ray data collection and model refinement statistics are summarized in Table S5. Figures showing the crystallographic model were generated and rendered with Pymol (Schrodinger, LLC). The atomic coordinates and structure factors of the CaM-P458 complexes have been deposited in the RCSB Protein Data Bank under the accession codes 6YNU (C2221 form) and 6YNS (P212121 form).

**HDX-MS analysis**

A summary of the Hydrogen/Deuterium eXchange – Mass Spectrometry (HDX-MS) experiments is provided in Table S7.

Biological samples

In order to determine the effect of P454 binding on CaM, the labeling was performed on CaM alone, and in the presence of a two-fold molar excess P454. HDX-MS experiments were carried out in buffer B, *i.e.*, in the presence of 2 mM $CaCl_2$. The quality of each protein and peptide was assessed prior to labeling by intact mass measurement (data not shown).

Sample preparation for HDX-MS

The P454:CaM complex was formed by mixing 10 µL of P454 (20 µM in buffer B) with 10 µL of CaM solution (10 µM in buffer B). The labeling control (*i.e.,* CaM alone) was prepared by replacing the P454 solution by 10 µL of buffer B. Prior to addition of the deuterated buffer B (99.98% $D_2O$ in 20 mM HEPES, 150 mM NaCl, 2 mM $CaCl_2$, pD 7.4), all solutions were equilibrated for 1 h at room temperature. Continuous labeling was initiated at 20˚C by adding

80 µL of deuterated buffer (80% final excess deuterium). Considering a $K_d$ of 90 nM and a 1 to 1 binding stoichiometry, 92.3% of CaM remains bound to P454 during labeling. The exchange reaction was quenched after 0.16, 1, 10, 30, 60 and 120 min by mixing 14 µL of the labeling reaction (*i.e.,* 14 pmoles of CaM) with 56 µL of quench buffer (0.6% formic acid) maintained at 4 ˚C to achieve a final pH of 2.5. Quenched samples were immediately snap-frozen in liquid nitrogen and stored at -80 ˚C. Triplicate labelling experiments were performed for each time point and condition for all HDX-MS analyses (independent technical replicates).

Data acquisition

HDX-MS analyses were performed with the aid of an HDX manager (Waters Corporation) maintained at 0 ºC. Prior to mass analysis, samples were rapidly thawed and 50 µL were digested for 2 min at 20 ºC using an in-house prepared column (2.0 X 20 mm, 66 µL bead volume) of immobilized pepsin beads (Thermo Scientific). Peptides were trapped, concentrated and desalted using a VanGuard™ CSH C18 pre-column (1.7 µm, 2.1 x 5 mm; Waters), and separated using an ACQUITY UPLC™ CSH C18 column (1.7 µm, 1 x 100 mm). Labeled peptides were eluted with an 8 min gradient of 5-30% acetonitrile at 40 µL/min and 0°C. After each run, the pepsin column was manually cleaned with two consecutive washes of 1.5% formic acid, 5% acetonitrile, 1.5 M guanidinium chloride, pH 1.6. Blank injections were performed after each sample to confirm the absence of carry-over.

The LC flow was directed to a Synapt™ G2-Si HDMS™ mass spectrometer (Waters Corporation) equipped with a standard electrospray ionization source (ESI). Mass accuracy was ensured by continuously infusing a Glu-1-Fibrinogen solution (100 fmol/µL in 50% acetonitrile, 0.1% formic acid) through the reference probe of the ESI source at a flow rate of 2.5 µL/min. Mass spectra were acquired in positive-ion and resolution mode over the 50–1950 *m/z* range. Peptides were identified in undeuterated samples using a combination of data-independent acquisition scheme ($MS^E$; trap collision energy ramp: 15.0 to 45.0 V) and exact mass measurement (<5.0 ppm mass error) using the same chromatographic settings than for the deuterated samples.

Data processing

Peptide maps were generated by database searching in ProteinLynx Global Server 3.0 (Waters Corporation, Milford, MA) using the following processing and workflow parameters: low and elevated intensity thresholds set to 100.0 and 50.0 counts; intensity threshold sets to 750.0 counts; automatic peptide and fragment tolerance; non-specific primary digest reagent; false discovery rate sets to 4%. Each fragmentation spectrum was manually inspected for assignment confirmation. Peptide map were refined in DynamX 3.0 (Waters Corporation, Milford, MA) using the following Import PLGS results filter: minimum intensity = 3000; minimum products per amino acid value = 0.4; minimum consecutive products = 1; minimum sum intensity for products = 1000; minimum score = 7; maximum MH+ error = 5 ppm. DynamX 3.0 HDX software (Waters Corporation, Milford, MA) was used to extract the centroid masses of all peptides selected for local HDX-analyses; only one charge state was considered per peptide. No adjustment was made for back-exchange and the results are reported as relative deuterium exchange levels expressed in either mass unit or fractional exchange. Fractional exchange data was calculated by dividing the experimentally measured uptake by the theoretically maximum number of exchangeable backbone amide hydrogens that could be replaced into each peptide (taking into account the final excess of deuterium present in the labeling mixture). MEMHDX was used to visualize and statistically validate HDX results (Wald test, false discovery rate sets to 5%) [34].

**Table S1**: Sedimentation coefficient of P454, CaM and P454:CaM complex at a 1:1 molar ratio. P454 and CaM were investigated alone (at 57 µM and 72 µM, respectively) and mixed in a 1:1 molar ratio (at 70 µM each). Experiments were performed at 20°C in buffer B with density at 1.0310 $g.mL^{-1}$ and viscosity at 1.0061 cP.

| | P454 | CaM | P454:CaM |
|---|---|---|---|
| Sedimentation coefficient (S) | 0.65 ± 0.1 | 1.85 ± 0.1 | 2.25 ± 0.1 |

**Table S2:** Thermodynamic parameters of P454:CaM titrations at different temperatures extracted from ITC experiments shown in Figure S3C-D.

| Temperature (K) | $K_A$ ($10^6$ $M^{-1}$) | $K_D$ (nM) | ΔH (kcal/mol) | ΔS (kcal/mol/K) | -TΔS (kcal/mol) | ΔG (kcal/mol) |
|---|---|---|---|---|---|---|
| 283 | 28 | 35 | 2.6 | 0.043 | -12.3 | -9.6 |
| 293 | 3.6 | 270 | -3.9 | 0.016 | -4.8 | -8.8 |
| 298 | 11 | 90 | -5.3 | 0.014 | -4.3 | -9.6 |
| 303 | 8 | 120 | -9.8 | -0.0009 | 0.3 | -9.5 |
| 311 | 6 | 160 | -13.5 | -0.012 | 3.8 | -9.6 |

ΔG, ΔH and TΔS : S.D. < 1 kcal/mol. S.D. for ΔS is below 10%.

S.D for $K_A$ and $K_D$ is below 15%

**Table S3:** Affinity constants and free energy values of the P454-derived peptides for POPC:POPG SUVs and CaM. The free energy $\Delta G_{Kx}$ for POPC:POPG was calculated from the partition coefficient $K_x$. The $K_d$ for CaM was calculated from fluorescent titrations and ITC experiments. Note that the $K_d$ values for CaM and for membranes are expressed in nM and µM, respectively.

| Peptides | $K_d$ for CaM (nM) | $\Delta G_{Kd}$ for CaM (kcal/mol) | Partition coefficient $K_X$ ($\times 10^3$) | $\Delta G_{Kx}$ * for POPC POPG 8:2 (kcal/mol) | $K_d$ ** for POPC:POPG 8:2 (µM) | µH *** |
|---|---|---|---|---|---|---|
| P454 | 90 | -9.6 | 790 | -8 | 70 | 0.422 |
| P454 R1Q | 100 | -9.5 | 190 | -7.2 | 290 | 0.422 |
| P454 R2Q | 180 | -9.2 | 150 | -7 | 370 | 0.366 |
| P454 R12Q | 2090 | -7.7 | 70 | -6.6 | 790 | 0.366 |
| P454 R12A | 130 | -9.4 | 280 | -7.4 | 200 | 0.329 |
| P454 R12K | 50 | -9.9 | 220 | -7.3 | 260 | 0.421 |
| P454 R12E | 22700 | -6.3 | 80 | -6.7 | 690 | 0.396 |
| P454 L463A | 200 | -9.1 | 630 | -7.9 | 90 | 0.422 |
| P454 L475A | 120 | -9.4 | 350 | -7.7 | 160 | 0.363 |
| P454 L481A | 90 | -9.6 | 780 | -8 | 70 | 0.433 |
| P454 I479A | 1430 | -7.9 | 330 | -7.5 | 170 | 0.303 |
| P454 I479V | 110 | -9.4 | 560 | -7.8 | 100 | 0.375 |
| P454 I479L | 60 | -9.8 | 700 | -7.9 | 80 | 0.414 |
| P454 H477S | 190 | -9.1 | 340 | -7.5 | 160 | 0.435 |
| P458 | 240 | -9.0 | 190 | -7.2 | 290 | 0.314 |
| P454 H477S-I479A (HSIA) | 1190 | -8.0 | 105 | -6.8 | 530 | 0.314 |
| P454 W458A-I479A (WAIA) | 2040 | -7.7 | 4.2 | -4.9 | 13200 | 0.303 |
| P454 L463A-L475A-H477S-I479A (EtoR) | 9090 | -6.8 | 35 | -6.2 | 1580 | 0.337 |
| P454 W458A-L463A (HNter) | 640 | -8.4 | 22 | -5.9 | 2550 | 0.422 |
| P454 L475A-H477S-I479A (HCter) | 1540 | -7.9 | 18 | -5.8 | 3160 | 0.314 |

* : the free energy of solution-to-membrane partiting is calculated from: $\Delta G_{Kx} = -RTLnKx$

**: The mole-fraction partition coefficient $K_X$ is converted into the dissociation constant $K_d$ using the following relation: $[W]/K_X = K_d$, where [W] is the water concentration (55.5 M).

***: Hydrophobic moment values are computed from Heliquest (with a window of 12) available at https://heliquest.ipmc.cnrs.fr/

S.D. values are within ± 10%.

**Table S4**: Thermodynamic parameters of the H:CaM ITC experiment performed at 25°C in buffer B shown in figure S7. A two-sites binding model was used to extract the thermodynamic parameters from the ITC data.

| | Stoichiometry (n) | $K_A$ ($10^6$ $M^{-1}$) | $K_D$ (nM) | ΔH (kcal/mol) | ΔS (kcal/mol/K | -TΔS (kcal/mol) | ΔG (kcal/mol) |
|---|---|---|---|---|---|---|---|
| Site 1 | 0.9 ± 0.1 | 114 ± 12 | 9 ± 2 | -5.4 ± 0.6 | 0.02 ± 0.002 | -5.6 ± 0.6 | -10.9 ± 1 |
| Site 2 | 1.03 ± 0.1 | 2.6 ± 0.3 | 380 ± 50 | -17.7 ± 2 | -0.03 ± 0.003 | 8.9 ± 0.9 | -8.7 ± 0.9 |

**Table S5A. Diffraction data collection and refinement statistics of P458:CaM crystal structures.**

| | C2221_form | P212121_form |
|---|---|---|
| **Crystallization conditions** | 0.3 M Ammonium sulfate<br>30% (w/v) PEG 4000 | 0.2M of MgCl2<br>0.1M HEPES-HCl pH 7.5<br>25% (w/v) PEG3350 |
| **Resolution range** | 67.6-3.11 (3.23 -3.11) | 88.86 -3.93 (4.07 - 3.93) |
| **Space group** | C 2 2 21 | P 21 21 21 |
| **Unit cell** | 73.341 174.344 97.843 90 90 90 | 97.001 106.974 221.635 90 90 90 |
| **Total reflections** | 103617 (10499) | 140108 (14505) |
| **Unique reflections** | 11538 (1132) | 21073 (2063) |
| **Multiplicity** | 9.0 (9.3) | 6.6 (7.0) |
| **Completeness (%)** | 99.49 (99.65) | 99.54 (99.76) |
| **Mean I/sigma(I)** | 22.97 (3.18) | 8.15 (3.03) |
| **Wilson B-factor** | 97.46 | 110.61 |
| **R-merge** | 0.055 (0.647) | 0.188 (0.736) |
| **R-meas** | 0.058 (0.684) | 0.204 (0.794) |
| **R-pim** | 0.018 (0.219) | 0.078 (0.297) |
| **CC1/2** | 1 (0.946) | 0.997 (0.875) |
| **CC*** | 1 (0.986) | 0.999 (0.966) |
| **Reflections used in refinement** | 11538 (1129) | 21073 (2061) |
| **Reflections used for R-free** | 592 (52) | 1074 (96) |
| **R-work** | 0.246 (0.385) | 0.221 (0.252) |
| **R-free** | 0.290 (0.488) | 0.284 (0.314) |
| **CC(work)** | 0.893 (0.913) | 0.960 (0.895) |
| **CC(free)** | 0.860 (0.777) | 0.945 (0.860) |
| **Number of non-hydrogen atoms** | 2665 | 17084 |
| **macromolecules** | 2656 | 17055 |
| **ligands** | 9 | 29 |
| **Protein residues** | 340 | 2209 |
| **RMS(bonds)** | 0.010 | 0.016 |
| **RMS(angles)** | 1.16 | 1.74 |
| **Ramachandran favored (%)** | 98.19 | 97.07 |
| **Ramachandran allowed (%)** | 1.51 | 2.51 |
| **Ramachandran outliers (%)** | 0.30 | 0.43 |

| | | |
|---|---|---|
| **Rotamer outliers (%)** | 0.00 | 0.00 |
| **Clashscore** | 8.85 | 19.35 |
| **Average B-factor** | 147.89 | 113.81 |
| **macromolecules** | 147.92 | 113.86 |
| **ligands** | 138.83 | 84.03 |
| **Number of TLS groups** | 10 | |
| **Molecules per ASU (CaM/P458)** | 2/2 | 12/24 |
| **PDB entry** | 6YNU | 6YNS |

Statistics for the highest-resolution shell are shown in parentheses.

Inaccessible residues
Solvent-accessible residues
HSDC Residues making Hydrogen/Disulphide bond, Salt bridge or Covalent link
Interfacing residues
**ASA** Accessible Surface Area, Å²  **BSA** Buried Surface Area, Å²  **Δ<sup>i</sup>G** Solvation energy effect, kcal/mol  |||| Buried area percentage, one bar per 10%

<table>
<tr><th>##</th><th>Structure 1</th><th>HSDC</th><th>ASA</th><th>BSA</th><th>ΔiG</th></tr>
<tr><td>1</td><td>D:TRP 458</td><td></td><td>271.16</td><td>0.00</td><td>0.00</td></tr>
<tr><td>2</td><td>D:GLY 459</td><td></td><td>42.33</td><td>0.00</td><td>0.00</td></tr>
<tr><td>3</td><td>D:GLN 460</td><td></td><td>158.25</td><td>0.00</td><td>0.00</td></tr>
<tr><td>4</td><td>D:ARG 461</td><td></td><td>225.16</td><td>0.00</td><td>0.00</td></tr>
<tr><td>5</td><td>D:ALA 462</td><td></td><td>55.33</td><td>0.00</td><td>0.00</td></tr>
<tr><td>6</td><td>D:LEU 463</td><td></td><td>81.81</td><td>0.00</td><td>0.00</td></tr>
<tr><td>7</td><td>D:GLN 464</td><td></td><td>125.76</td><td>30.14 |||</td><td>-0.22</td></tr>
<tr><td>8</td><td>D:GLY 465</td><td></td><td>36.08</td><td>0.00</td><td>0.00</td></tr>
<tr><td>9</td><td>D:ALA 466</td><td></td><td>58.78</td><td>0.00</td><td>0.00</td></tr>
<tr><td>10</td><td>D:GLN 467</td><td></td><td>121.94</td><td>0.00</td><td>0.00</td></tr>
<tr><td>11</td><td>D:ALA 468</td><td></td><td>58.38</td><td>40.38 |||||||</td><td>0.56</td></tr>
<tr><td>12</td><td>D:VAL 469</td><td></td><td>97.90</td><td>14.59 ||</td><td>0.21</td></tr>
<tr><td>13</td><td>D:ALA 470</td><td></td><td>57.59</td><td>0.00</td><td>0.00</td></tr>
<tr><td>14</td><td>D:ALA 471</td><td></td><td>53.67</td><td>27.62 ||||||</td><td>0.44</td></tr>
<tr><td>15</td><td>D:ALA 472</td><td></td><td>52.18</td><td>52.02 ||||||||||</td><td>0.81</td></tr>
<tr><td>16</td><td>D:GLN 473</td><td></td><td>92.30</td><td>15.70 ||</td><td>-0.11</td></tr>
<tr><td>17</td><td>D:ARG 474</td><td></td><td>190.61</td><td>0.00</td><td>0.00</td></tr>
<tr><td>18</td><td>D:LEU 475</td><td></td><td>118.41</td><td>94.55 ||||||||</td><td>1.51</td></tr>
<tr><td>19</td><td>D:VAL 476</td><td></td><td>82.31</td><td>77.99 ||||||||||</td><td>1.11</td></tr>
<tr><td>20</td><td>D:HIS 477</td><td></td><td>95.24</td><td>31.60 ||||</td><td>0.62</td></tr>
<tr><td>21</td><td>D:ALA 478</td><td></td><td>55.07</td><td>29.39 ||||||</td><td>0.09</td></tr>
<tr><td>22</td><td>D:ILE 479</td><td></td><td>146.52</td><td>141.87 ||||||||||</td><td>1.82</td></tr>
<tr><td>23</td><td>D:ALA 480</td><td></td><td>93.41</td><td>75.23 |||||||||</td><td>0.65</td></tr>
<tr><td>24</td><td>D:LEU 481</td><td></td><td>177.28</td><td>27.72 ||</td><td>0.04</td></tr>
</table>

Table S5B. P454 residues involved in the interaction with the N-lobe of CaM.

<table>
<tr><th>##</th><th>Structure 1</th><th>HSDC</th><th>ASA</th><th>BSA</th><th>ΔiG</th></tr>
<tr><td>1</td><td>B:TRP 458</td><td>H</td><td>280.59</td><td>241.34 |||||||||</td><td>2.17</td></tr>
<tr><td>2</td><td>B:GLY 459</td><td></td><td>41.61</td><td>0.00</td><td>0.00</td></tr>
<tr><td>3</td><td>B:GLN 460</td><td></td><td>142.80</td><td>16.89 ||</td><td>-0.19</td></tr>
<tr><td>4</td><td>B:ARG 461</td><td></td><td>245.37</td><td>5.53 |</td><td>-0.05</td></tr>
<tr><td>5</td><td>B:ALA 462</td><td></td><td>64.72</td><td>54.60 |||||||||</td><td>0.63</td></tr>
<tr><td>6</td><td>B:LEU 463</td><td></td><td>65.71</td><td>53.88 |||||||||</td><td>0.80</td></tr>
<tr><td>7</td><td>B:GLN 464</td><td></td><td>111.46</td><td>0.00</td><td>0.00</td></tr>
<tr><td>8</td><td>B:GLY 465</td><td></td><td>33.89</td><td>12.00 ||||</td><td>0.18</td></tr>
<tr><td>9</td><td>B:ALA 466</td><td></td><td>62.78</td><td>62.78 ||||||||||</td><td>0.86</td></tr>
<tr><td>10</td><td>B:GLN 467</td><td></td><td>115.46</td><td>75.30 |||||||</td><td>-0.09</td></tr>
<tr><td>11</td><td>B:ALA 468</td><td></td><td>62.88</td><td>0.00</td><td>0.00</td></tr>
<tr><td>12</td><td>B:VAL 469</td><td></td><td>97.01</td><td>42.40 |||||</td><td>0.68</td></tr>
<tr><td>13</td><td>B:ALA 470</td><td></td><td>57.70</td><td>47.98 |||||||||</td><td>0.76</td></tr>
<tr><td>14</td><td>B:ALA 471</td><td></td><td>54.54</td><td>0.17 |</td><td>0.00</td></tr>
<tr><td>15</td><td>B:ALA 472</td><td></td><td>50.81</td><td>0.00</td><td>0.00</td></tr>
<tr><td>16</td><td>B:GLN 473</td><td></td><td>91.56</td><td>0.00</td><td>0.00</td></tr>
<tr><td>17</td><td>B:ARG 474</td><td>HS</td><td>192.35</td><td>49.69 |||</td><td>-0.54</td></tr>
<tr><td>18</td><td>B:LEU 475</td><td></td><td>117.88</td><td>0.00</td><td>0.00</td></tr>
<tr><td>19</td><td>B:VAL 476</td><td></td><td>84.08</td><td>0.00</td><td>0.00</td></tr>
<tr><td>20</td><td>B:HIS 477</td><td></td><td>99.60</td><td>0.00</td><td>0.00</td></tr>
<tr><td>21</td><td>B:ALA 478</td><td></td><td>67.87</td><td>0.00</td><td>0.00</td></tr>
<tr><td>22</td><td>B:ILE 479</td><td></td><td>146.78</td><td>0.00</td><td>0.00</td></tr>
<tr><td>23</td><td>B:ALA 480</td><td></td><td>80.63</td><td>0.00</td><td>0.00</td></tr>
<tr><td>24</td><td>B:LEU 481</td><td></td><td>181.92</td><td>0.00</td><td>0.00</td></tr>
</table>

Table S5C. P454 residues involved in the interaction with the C-lobe of CaM.

**Table S6**: Data collection and scattering derived parameters from SAXS.

| | CaM | P454-CaM | *H-CaM*[a] | *MLCK-CaM*[a] |
|---|---|---|---|---|
| **Data collection parameters** | | | | |
| Instrument | SWING (SOLEIL) | | | |
| Detector | CCD-based AVIEX | | | |
| Beam geometry | 0.8 mm x 0.15 mm | | | |
| Wavelength [Å] | 1.0 | | | |
| q-range [$Å^{-1}$] | $0.006 < q < 0.50$ | | | |
| Absolute scaling | Comparison with scattering from pure $H_2O$ | | | |
| Exposure (Dead) time [s] | 1/0.5 | | | |
| Temperature [K] | 288 | | | |
| SEC-SAXS column | Biosec 3 Agilent | | | |
| Loading concentration[μM] | 409 | 136 (CaM)-272(P454) | | |
| Injection volume [μL] | 50 | | | |
| Buffer | 20 mM Hepes, 150 mM NaCl, 4 mM CaCl2, pH 7.4 | | | |
| Flow rate [$mL.min^{-1}$] | 0.2 | | | |
| **Software employed for SAXS data reduction and analysis** | | | | |
| Foxtrot | SWING in-house software | | | |
| PRIMUS | ATSAS 2.8 suite, Franke et al. J. Appl. Cryst. 2017 | | | |
| DADIMODO | Roudenko O., Thureau A. & Pérez J., 2018 | | | |
| DAMMIF/DAMMIN | D. Franke and D. I. Svergun, J. Appl. Cryst. 2009 | | | |
| **Structural parameters** | | | | |
| I(0) Guinier [$cm^{-1}$] | 0.02576 ± 0.00004 | 0.00781 ± 0.00001 | *0.0159* | *0.0134* |
| $R_g$ Guinier [Å] | 22.16 ± 0.1 | 19.74 ± 0.1 | *21.9* | *17.4* |
| qRg-range | 0.250 - 1.03 | 0.256 - 1.30 | | |
| I(0) p(r) [$cm^{-1}$] | 0.02574 ± 0.00004 | 0.00783 ± 0.00001 | *0.0159* | *0.0134* |
| $R_g$ p(r) [Å] | 22.25 ± 0.1 | 19.96 ± 0.1 | *22.1* | *17.3* |
| q-range [$Å^{-1}$] | 0.0113 - 0.35 | 0.013 - 0.35 | | |
| $D_{Max}$ [Å] | 73 | 70 | *75* | *52* |
| **Molecular mass determination** | | | | |
| $MM_{sequence}$ [kDa] [b] | 16.8 | 20.1 | | |
| $MM_{SAXS\ MoW}$ [kDa] [c] | 17.2 | 19.9 | | |
| $MM_{SAXS\ QR}$ [kDa] [d] | 16.4 | 19.1 | | |
| **DAMMIF/DAMMIN analysis** | | | | |
| Model number | 20 | 20 | | |
| NSD | 0.56 | 0.74 | | |
| $\chi^2$ (Dammin) | 1.56 | 1.34 | | |
| **DADIMODO analysis** | | | | |
| Model number | | 20 | | |
| q-range [$Å^{-1}$] | | 0.016-0.50 | | |
| $\chi^2$ (Crysol) | | 1.49-1.70 | | |

[a] Values extracted from Table S1 in O'Brien *et al.*, *PLoS Biol* **2017,** *15* (12), e2004486 [4].

[b] The calculated masses were derived from the sequences.

[c] Molecular mass M obtained from the whole I(q) curve ($q_{max}$ = 0.3$Å^{-1}$) using the SAXS-MoW2 program, available at http://saxs.ifsc.usp.br/.

[d] Molecular mass M obtained from the whole I(q) curve ($q_{max}$ = 0.25 $Å^{-1}$) using the ScÅtter3 program, available at https://bl1231.als.lbl.gov/scatter/.

**Table S7**: HDX Data Summary

| **EXPERIMENT** | **P454 BINDING TO CaM** | |
|---|---|---|
| Dat set : | CaM_Ca | CaM_Ca_P454 |
| HDX reaction details | | |
| ○ *pD* | 7.4 | 7.4 |
| ○ *T°C* | 20°C | 20°C |
| ○ *Excess deuterium* | 80% | 80% |
| HDX time course analyzed (min) | 0.16, 1, 10, 30, 60, 120 | 0.16, 1, 10, 30, 60, 120 |
| Number of peptides | 46 | 46 |
| Sequence coverage | 89.3 | 89.3 |
| Average peptide length | 11.7 | 11.7 |
| Redundancy | 4.01 | 4.01 |
| Average peptide length/Redundancy ratio | 2.91 | 2.91 |
| Replicates | 3 (technical) | 3 (technical) |
| Repeatability (pooled standard deviation)[#] | 0.07 Da | 0.07 Da |
| Significant difference between state[$] | Wald test, $p < 0.05$ | |

[#] One unique charge state was used per peptide
[$] MEMHDX (www. memhdx.c3bi.pasteur.fr)

**Table S8**: Estimation of the α-helical content of P454-derived peptides. The secondary structure content is estimated using the on-line Bestsel software [17] available at http://bestsel.elte.hu/index.php. The alpha-helical content (%) is reported in the table for each peptide in buffer A, in 20% TFE and in the presence of POPC POPG (molar ratio 8:2) at 2 mM lipid concentration. The far-UV CD spectra of several P454-derived peptides are shown in figure S12.

| P454 derived peptide | Solution (helix, %) | TFE 20% (helix, %) | POPC:POPG 8:2 (helix, %) |
|---|---|---|---|
| P454 | 17 ± 2 | 60 ± 2 | 50 ± 1 |
| P454 R12Q | 8 ± 4 | 50 ± 2 | 23 ± 1 |
| P454 R12K | 13 ±2 | 45 ± 4 | 26 ± 1 |
| P454 R12E | 6 ± 1 | 55 ± 3 | 10 ± 1 |
| P454 L463A | 14 ± 2 | 48 ± 3 | 19 ± 2 |
| P454 L475A | 5 ± 2 | 45 ± 1 | 15 ± 1 |
| P454 L481A | 5 ± 1 | 49 ± 1 | 44 ± 3 |
| P454 I479A | 8 ± 2 | 29 ± 1 | 12 ± 3 |
| P454 I479V | 2 ± 1 | 47 ± 6 | 37 ± 2 |
| P454 I479L | 13 ± 1 | 51 ± 1 | 51 ± 5 |
| P454 H477S | 4 ± 2 | 40 ± 1 | 20 ± 2 |
| P458 | 10 ± 4 | 50 | 23 ± 2 |
| P454 W458A-I479A | 10 ± 3 | 55 ± 1 | 7 ± 3 |
| P454 L463A-L475A-H477S-I479A (EtoR) | 4 ± 1 | 38 ± 2 | 5 ± 2 |
| P454 W458A- L463A (HNt) | 7 ± 1 | 56 | 8 ± 2 |
| P454 H477S-I479A | 2 ± 2 | 51 ± 4 | 27 ± 3 |
| P454 L475A-H477S-I479A (HCt) | 6 ± 2 | 51 ± 4 | 6 ± 2 |

**Table S9**: Concentrations of P454-derived peptides required to induce the recovery of 50% of ANTS fluorescence intensity upon LUVs permeabilization ($C_{P1/2}$) (see Material and Methods for details). Vesicle permeabilization induced by P454-derived peptides is blocked by the addition of calmodulin in the presence of calcium (see Material and Methods for details), except for the peptides (i.e., R12E, EtoR and W458A-I479A) that cannot be assayed as they do not permeabilize vesicles. Substituted residues are bold in the peptide sequences. S.D.: ± 5%, excepted for R12E, W458A-I479A and EtoR for which the $Cp_{1/2}$ values cannot be determined.

| Peptide name (nickname) | $C_{P1/2}$ (µM) | Peptide sequences |
|---|---|---|
| P454 WT | 0.07 | ASAHWGQRALQGAQAVAAAQRLVHAIALMTQ |
| P454 R461Q (R*1*Q) | 0.22 | ASAHWGQ**Q**ALQGAQAVAAAQRLVHAIALMTQ |
| P454 R474Q (R*2*Q) | 0.11 | ASAHWGQRALQGAQAVAAAQ**Q**LVHAIALMTQ |
| P454 R461Q R474Q (R*12*Q) | 0.23 | ASAHWGQ**Q**ALQGAQAVAAAQ**Q**LVHAIALMTQ |
| P454 R461A R474A (R*12*A) | 0.14 | ASAHWGQ**A**ALQGAQAVAAAQ**A**LVHAIALMTQ |
| P454 R461K R474K (R*12*K) | 0.09 | ASAHWGQ**K**ALQGAQAVAAAQ**K**LVHAIALMTQ |
| P454 R461E R474E (R*12*E) | > 50 | ASAHWGQ**E**ALQGAQAVAAAQ**E**LVHAIALMTQ |
| P454 L463A (L463A) | 0.41 | ASAHWGQRA**A**QGAQAVAAAQRLVHAIALMTQ |
| P454 L475A (L475A) | 0.64 | ASAHWGQRALQGAQAVAAAQR**A**VHAIALMTQ |
| P454 L481A (L481A) | 0.06 | ASAHWGQRALQGAQAVAAAQRLVHAIA**A**MTQ |
| P454 I479A (I479A) | 7.86 | ASAHWGQRALQGAQAVAAAQRLVHA**A**ALMTQ |
| P454 I479V (I479V) | 0.09 | ASAHWGQRALQGAQAVAAAQRLVHA**V**ALMTQ |
| P454 I479L (I479L) | 0.04 | ASAHWGQRALQGAQAVAAAQRLVHA**L**ALMTQ |
| P454 H477S (H477S) | 0.25 | ASAHWGQRALQGAQAVAAAQRLV**S**AIALMTQ |
| P458 | 1.26 | WGQRALQGAQAVAAAQRLVHAIAL |
| P454 H477S-I479A | 1.88 | ASAHWGQRALQGAQAVAAAQRLV**S**A**A**ALMTQ |
| P454 W458A-I479A | > 80 | ASAH**A**GQRALQGAQAVAAAQRLVHA**A**ALMTQ |
| P454 L463A-L475A-H477S-I479A | > 50 | ASAHWGQRA**A**QGAQAVAAAQR**A**V**S**A**A**ALMTQ |
| P454 W458A-L463A | 6.59 | ASAH**A**GQRA**A**QGAQAVAAAQRLVHAIALMTQ |
| P454 L475A-H477S-I479A | 9.03 | ASAHWGQRALQGAQAVAAAQR**A**V**S**A**A**ALMTQ |

**A**

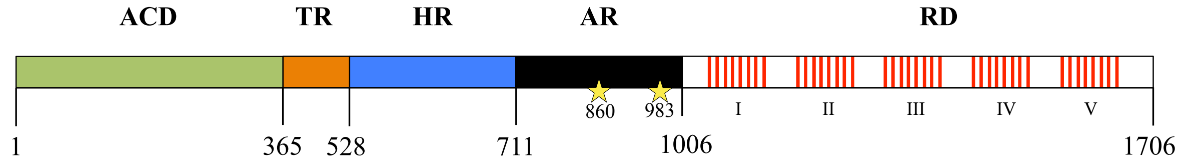


**B**

MQQSHQAGYANAADRESGIPAAVLDGIKAVAKEKNATLMFRLVNPHSTSLIAEGVATKGLGVHAKSSD
WGLQAGYIPVNPNLSKLFGRAPEVIARADNDVNSSLAHGHTAVDLTLSKERLDYLRQAGLVTGMADGV
VASNHAGYEQFEFRVKETSDGRYAVQYRRKGGDDFEAVKVIGNAAGIPLTADIDMFAIMPHLSNFRDS
ARSSVTSGDSVTDYLARTRRAASEATGGLDRERIDLLWKIARAGARSAVGTEARRQFRYDGDMNIGVI
TDFELEVRNALNRRAHAVGAQDVVQHGTEQNNPFPEADEKIFVVSATGESQMLTRGQLKEYIGQQRGE
GYVFYENRAYGVAGKSLFDDGLGAAPGVPSGRSKFSPDVLETVPASPGLRRPSLGAVERQDSGYDSLD
GVGSRSFSLGEVSDMAAVEAAELEMTRQVLHAGARQDDAEPGVSG**ASAHWGQRALQGAQAVAAAQRLV**
**HAIALMTQ**FGRAGSTNTPQEAASLSAAVFGLGEASSAVAETVSGFFRGSSRWAGGFGVAGGAMALGGG
IAAAVGAGMSLTDDAPAGQKAAAGAEIALQLTGGTVELASSIALALAAARGVTSGLQVAGASAGAAAG
ALAAALSPMEIYGLVQQSHYADQLDKLAQESSAYGYEGDALLAQLYRDKTAAEGAVAGVSAVLSTVGA
AVSIAAAASVVGAPVAVVTSLLTGALNGILRGVQQPIIEKLANDYARKIDELGGPQAYFEKNLQARHE
QLANSDGLRKMLADLQAGWNASSVIGVQTTEISKSALELAAITGNADNLKSVDVFVDRFVQGERVAGQ
PVVLDVAAGGIDIASRKGERPALTFITPLAAPGEEQRRRTKTGKSEFTTFVEIVGKQDRWRIRDGAAD
TTIDLAKVVSQLVDANGVLKHSIKLDVIGGDGDDVVLANASRIHYDGGAGTNTVSYAALGRQDSITVS
ADGERFNVRKQLNNANVYREGVATQTTAYGKRTENVQYRHVELARVGQLVEVDTLEHVQHIIGGAGND
SITGNAHDNFLAGGSGDDRLDGGAGNDTLVGGEGQNTVIGGAGDDVFLQDLGVWSNQLDGGAGVDTVK
YNVHQPSEERLERMGDTGIHADLQKGTVEKWPALNLFSVDHVKNIENLHGSRLNDRIAGDDQDNELWG
HDGNDTIRGRGGDDILRGGLGLDTLYGEDGNDIFLQDDETVSDDIDGGAGLDTVDYSAMIHPGRIVAP
HEYGFGIEADLSREWVRKASALGVDYYDNVRNVENVIGTSMKDVLIGDAQANTLMGQGGDDTVRGGDG
DDLLFGGDGNDMLYGDAGNDTLYGGLGDDTLEGGAGNDWFGQTQAREHDVLRGGDGVDTVDYSQTGAH
AGIAAGRIGLGILADLGAGRVDKLGEAGSSAYDTVSGIENVVGTELADRITGDAQANVLRGAGGADVL
AGGEGDDVLLGGDGDDQLSGDAGRDRLYGEAGDDWFFQDAANAGNLLDGGDGRDTVDFSGPGRGLDAG
AKGVFLSLGKGFASLMDEPETSNVLRNIENAVGSARDDVLIGDAGANVLNGLAGNDVLSGGAGDDVLL
GDEGSDLLSGDAGNDDLFGGQGDDTYLFGVGYGHDTIYESGGGHDTIRINAGADQLWFARQGNDLEIR
ILGTDDALTVHDWYRDADHRVEIIHAANQAVDQAGIEKLVEAMAQYPDPGAAAAAPPAARVPDTLMQS
LAVNWR

**Figure S1**: (**A**) Schematic representation of the CyaA toxin. The toxin is composed of the following domains: a N-terminal catalytic domain (ACD, residues 1-364), a translocation region (TR, 365-527), a hydrophobic region (HR, 528-710), an acylation region (AR, 711-1005) with acylated K860 and K983 labeled with yellow stars, and a C-terminal receptor-binding domain (RD, 1006-1706) containing the Repeat-in-ToXin (RTX) motifs (red bars). (**B**) Sequence of the full-length CyaA toxin from *Bordetella pertussis*. The region corresponding to the P454 peptide is bold and underlined. The same color code is used in panels A and B.

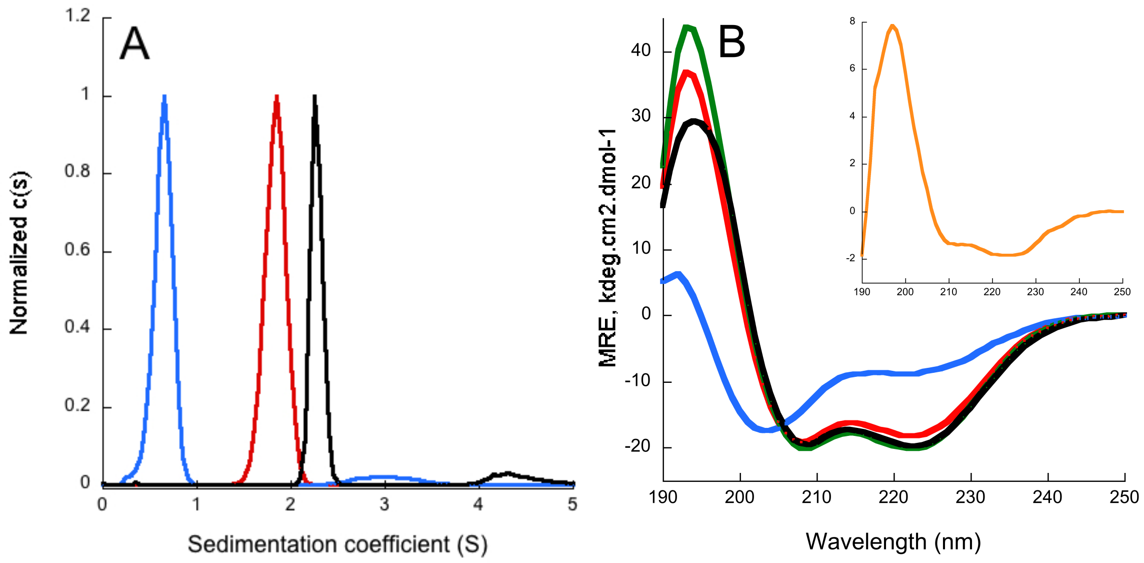


**Figure S2**: (**A**) Sedimentation coefficient distributions of P454 (blue), CaM (red) and the P454:CaM complex (black). Sedimentation coefficients are listed in Table S1. (**B**) Synchrotron radiation circular dichroism in the far-UV region of P454 and CaM. Spectra were recorded in a 100 µm cell. Far-UV spectra of P454 (blue), CaM (green) or P454:CaM at a 1:1 molar ratio (black) in buffer B. The red spectrum corresponds to the combination of the isolated CaM and P454 spectra. The inset corresponds to the structural changes induced by the P454:CaM complex formation (subtraction of the black spectrum by the red spectrum). The subtraction spectrum shows that the P454:CaM complex formation induces an increase of secondary structure content. Experimental conditions: buffer B, temperature: 25°C.

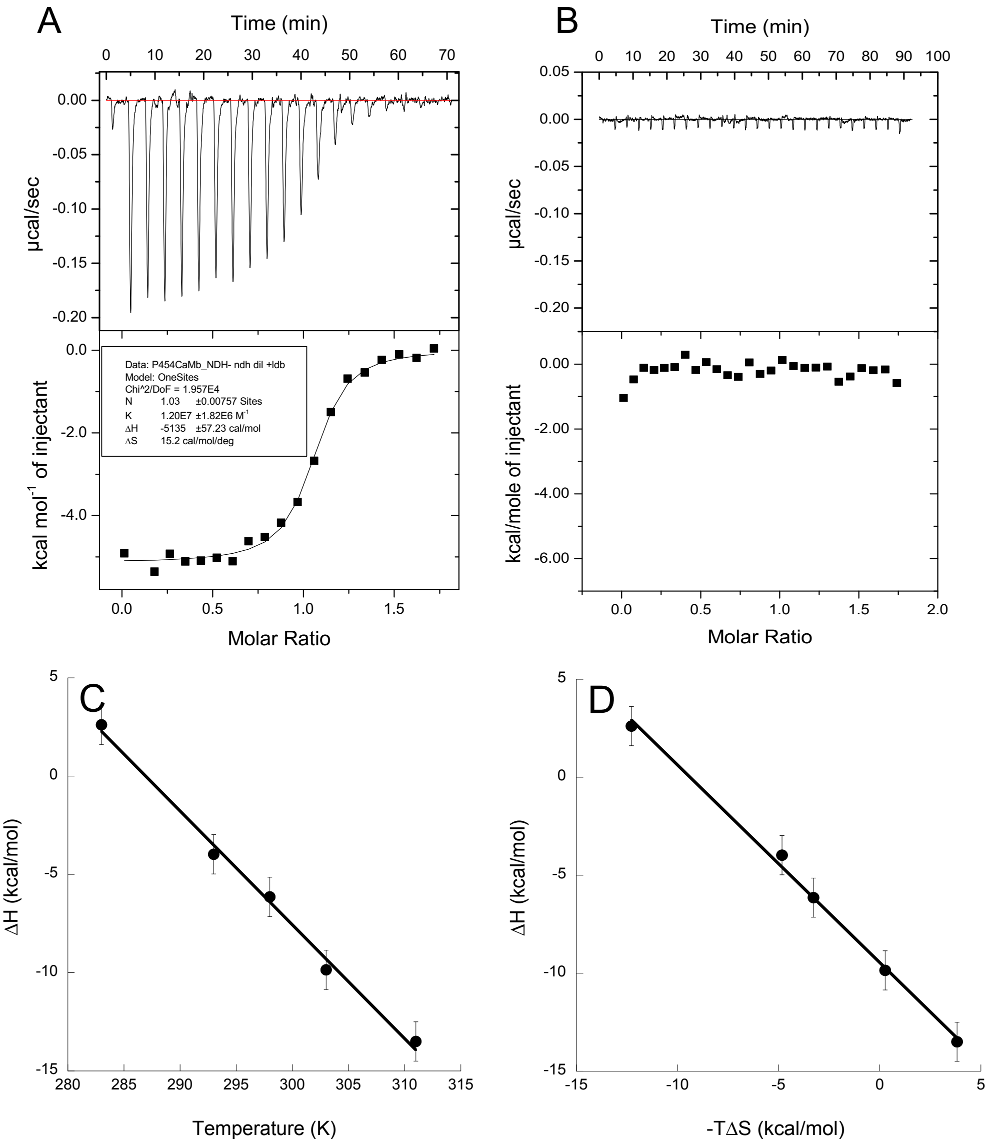


**Figure S3**: (**A**) Isothermal titration calorimetry of CaM by P454 in buffer B at 25°C using a VP-ITC instrument. Titration was performed by injections of 13 µL of P454 (loaded in the syringe at 90 µM) into the cell containing CaM at 10 µM (see Table S3 for ΔG values). (**B**) Isothermal titration calorimetry of CaM by P454 in buffer A complemented with 0.2 mM EDTA at 25°C. Titration was performed by consecutive injections of 10 µL of P454 (loaded in the syringe at 90 µM) into the cell containing CaM at 10 µM. (**C**) ITC experiments were performed at various temperatures to infer the heat capacity of the reaction. The slope of ΔH=f(T) provides us with the heat capacity ΔCp of the P454:CaM complex. The ΔCp value is negative, as measured for canonical CaM-binding peptides [35], however, lower than those measured for globular peptide:CaM complexes, suggesting that the P454:CaM complex does not adopt a globular conformation. This is confirmed by X-ray and SAXS data reported in this article. (**D**) Representation of the enthalpy-entropy compensation (EEC) plot derived from ITC experiments of P454:CaM titrations at different temperatures. The P454:CaM complex formation at various temperatures follows a typical entropy-enthalpy compensation. The enthalpy and temperature values are listed in Table S2. S.D for ΔH values is ± 1 kcal/mol.

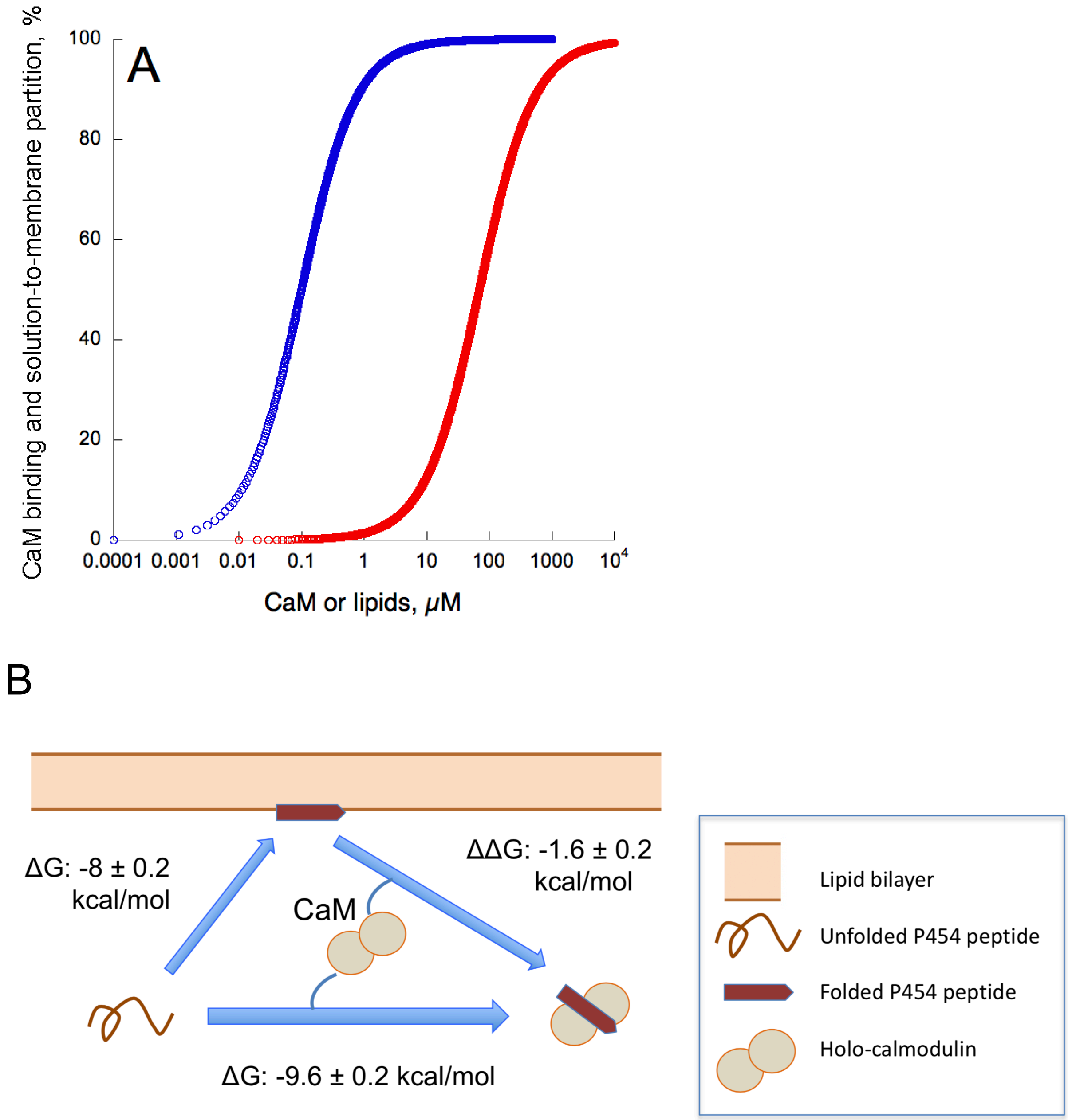


**Figure S4**: (**A**) Fraction of P454 bound to CaM and membrane, calculated from the affinity constants for CaM ($K_d$=90 nM, blue trace) and POPC POPG 8:2 membranes ($K_d$=70 µM, red trace). The mole-fraction partition coefficient $K_X$ is converted into $K_d$ using the following relation: $[W]/K_X = K_d$, where [W] is the water concentration (55.5 M). (**B**) Scheme of P454 interactions with membranes and CaM. The P454 peptide is represented in red and CaM by two grey lobes. The free energy of interaction of P454 with membrane and with CaM was calculated from its $K_X$ and $K_d$ values, as described in the materials and methods section. The $K_X$ and $K_d$ values are listed in Table S3.

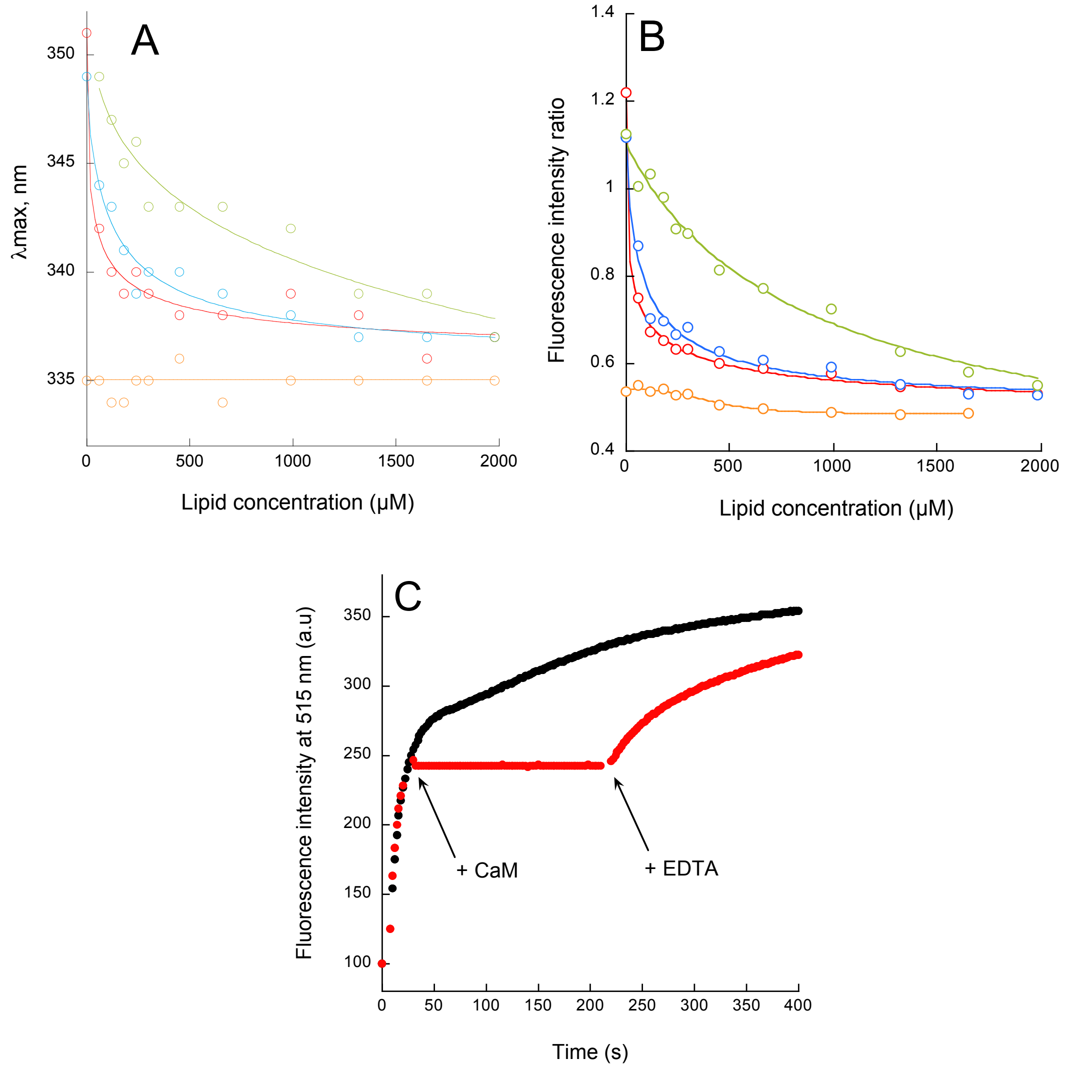


**Figure S5**: P454 membrane interaction and permeabilization followed by fluorescence. Maximum wavelength of emission (**A**) and ratio of fluorescence intensity (**B**) of P454 followed by tryptophan fluorescence as a function of lipid concentration. A sample of P454 is prepared for each lipid concentration. Tryptophan fluorescence of P454 was first measured for each concentration of lipids with SUVs composed of POPC POPG 8:2 (red circles) in buffer A. Then, an excess of apo-CaM is added to each sample (blue circles, molar ratio P454:CaM 1:3), followed by 1 mM of calcium (orange circles) and finally 2 mM EDTA (green circles). (**C**) Membrane permeabilization assay. Time course of ANTS and DPX efflux from LUVs made of POPC POPG 8:2 (0.45 mM lipid concentration) in buffer B, *i.e.*, including 2 mM $CaCl_2$. Experiment was performed in the presence of P454 at 100 nM (black curve). In red, the same experiment is performed but holo-CaM is added (molar ratio P454:CaM 1:3) at 40 sec, followed by addition of 5 mM EDTA (red curve).

**Figure S6:** Droplet Interfacial Bilayers (DIB) experiments.

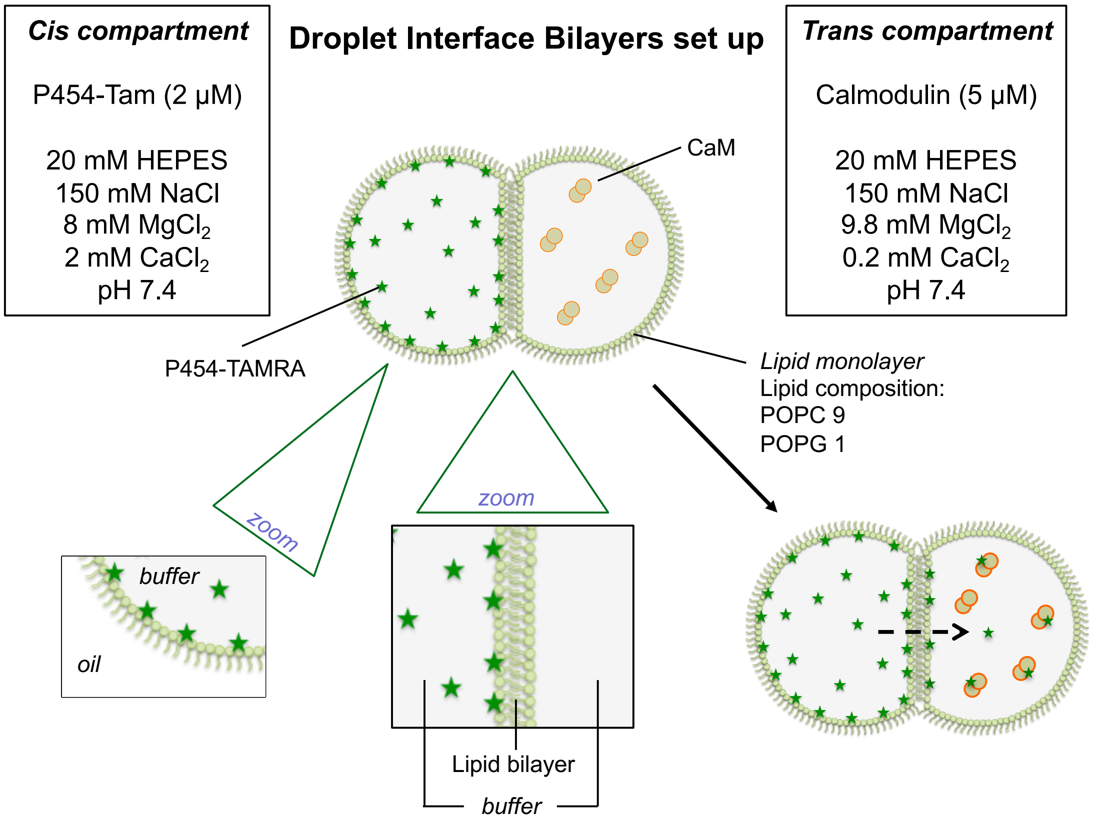


Figure S6A: Scheme of the Droplet Interface Bilayer (DIB) experimental set-up.

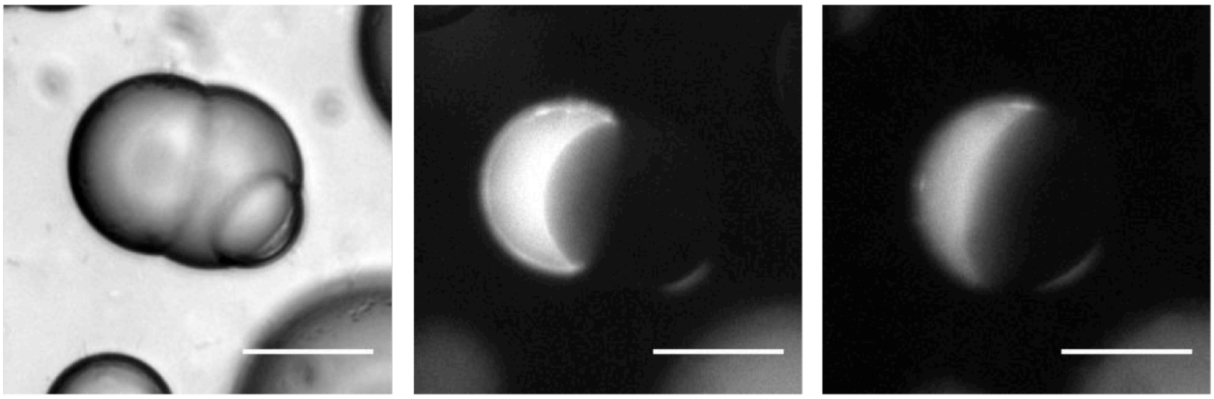

Figure S6B: Pictures of a DIB made of 10% POPG 90% POPC with P454 (2 μM) in the left droplet and without CaM in the right droplet. On the left: brightfield microscopy image. In the middle: fluorescence microscopy image at the beginning of the experiment (t=0min). On the right: fluorescence microscopy image after 15 minutes of experiment. No fluorescence was observed in the *trans* compartment in this experiment. Scale bar represents 30 μm.

*Figure S6 continued on next page*

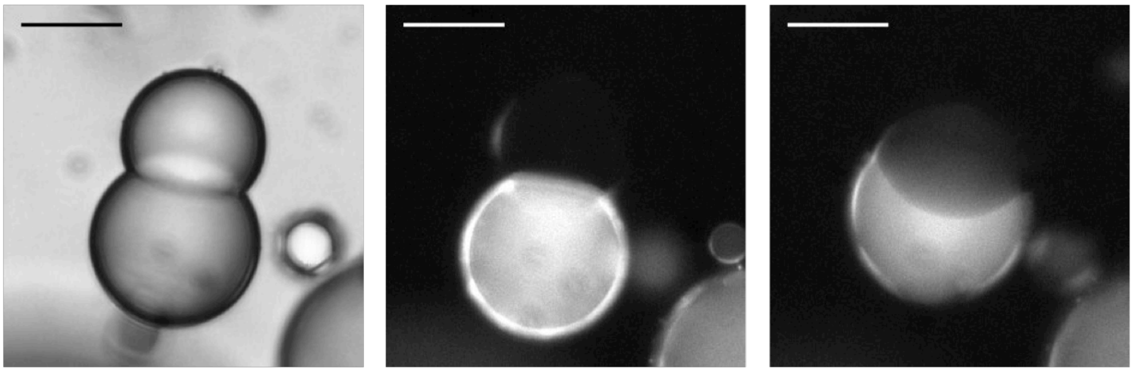

Figure S6C: Images of a DIB made of 10% POPG 90% POPC with P454 (2 µM) in the bottom droplet and CaM (5 µM) in the upper droplet. On the left: brightfield microscopy image. In the middle: fluorescence microscopy image at the beginning of the experiment (t=0min). On the right: fluorescence microscopy image after 15 minutes of experiment. Translocation was observed in this experiment. Scale bar represents 30 µm.

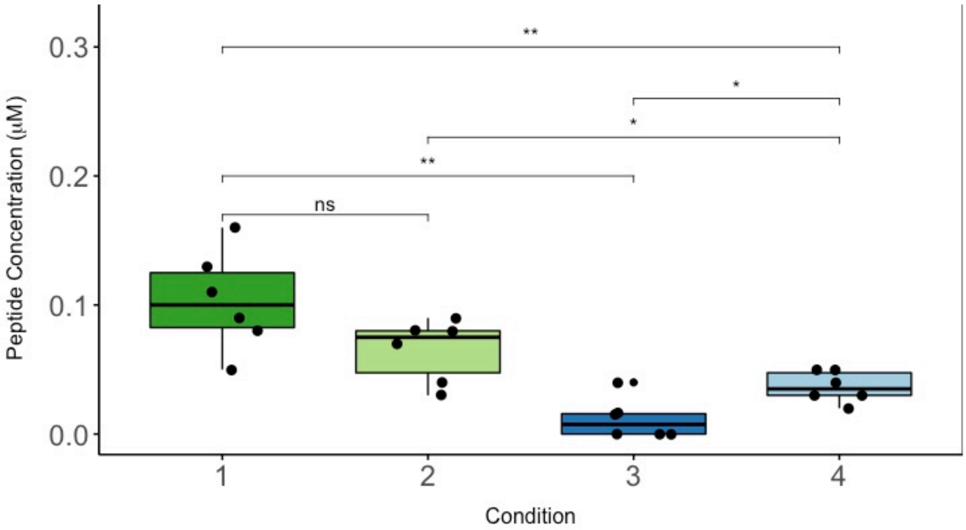


Figure S6D: Peptide translocation across droplet interface bilayers. Boxplot representation of the TAMRA-P454 concentration (µM) in the *trans* droplet 15 min after the formation of the droplet interface bilayers (see Material and Methods for details). All experiments were performed in the presence of a calcium gradient across the lipid bilayer ($CaCl_2$: 2 mM *cis vs* 0.2 mM *trans*). Peptide concentration in the *trans* droplet of TAMRA-P454 WT (green conditions, 1-2) and TAMRA-P454 R12E (blue conditions, 3-4) was measured in the presence of 5 µM of CaM (dark colors, 1 and 3) and 100 µM of CaM (light colors, 2 and 4) in the *trans* droplet. Five to seven independent trials were conducted for each condition. Mann-Whitney-Wilcoxon test was applied to compare experiments.

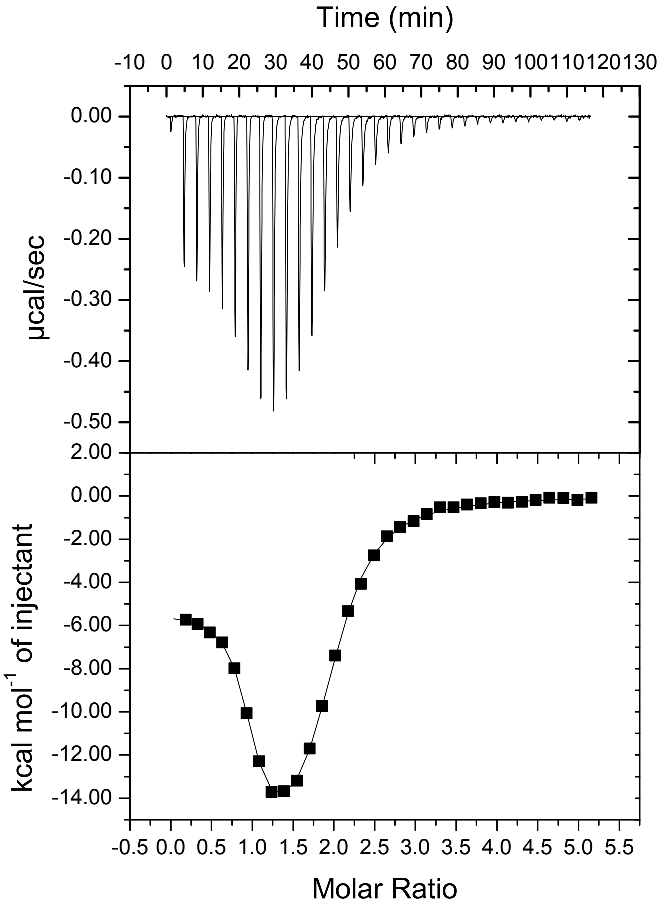


**Figure S7**: Isothermal titration calorimetry of holo-CaM by the H-helix peptide in buffer B at 25°C. The H-helix peptide is loaded in the syringe at 130 µM and CaM in the cell at 5 µM. Titration was performed on VP-ITC with consecutive injections of 8 µL. A two-sites binding model was used to fit to the ITC data and to extract the thermodynamic parameters reported in Table S4.

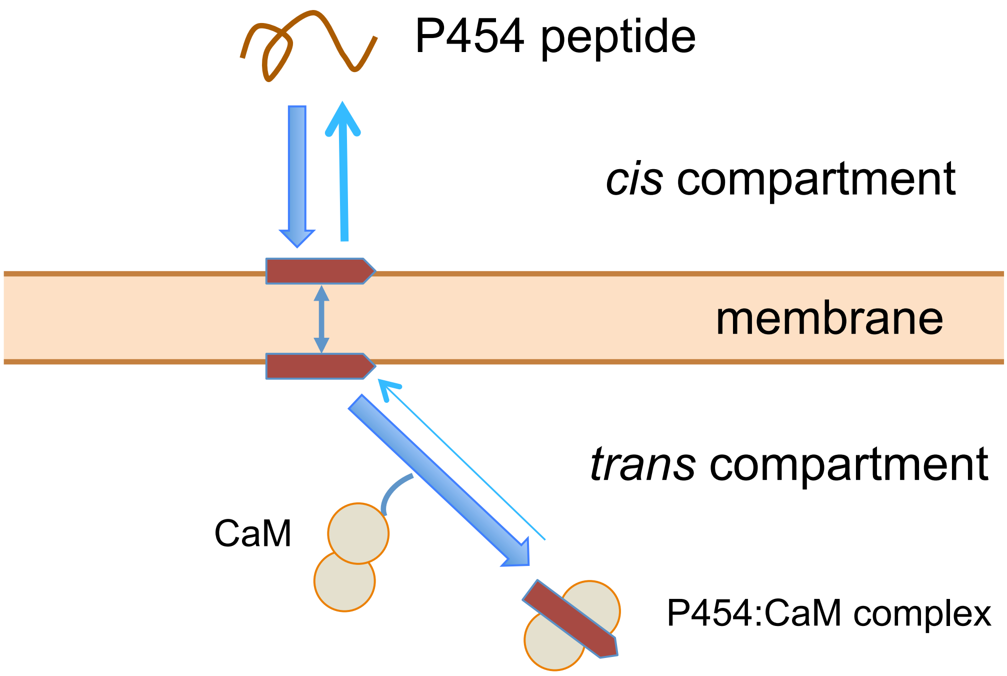


**Figure S8**: Scheme of the P454 translocation and CaM binding inferred from the *in vitro* experiments based on the ITC, fluorescence, SRCD and DIB experiments described in this study.

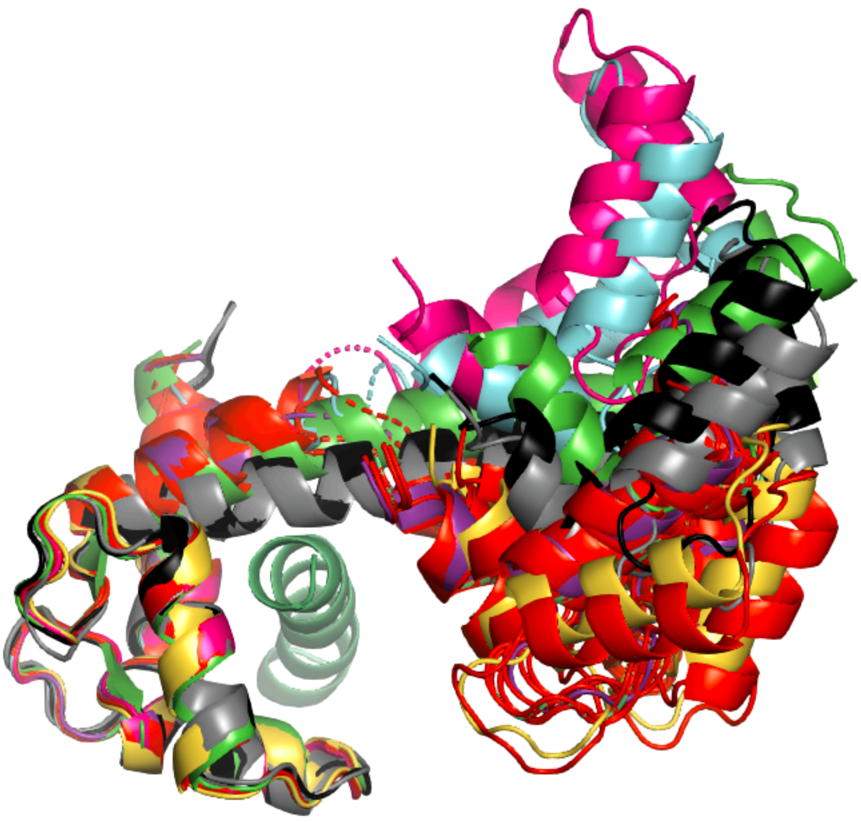

**Figure S9:** Superimposition of the fourteen P458:CaM crystal structures. The crystal structure 1CLL [36] of the extended conformation of CaM is shown in light green. The 2 crystal structures 6YNU (black and grey) and the 12 crystal structures 6YNS are displayed with the N-CaM lobe superimposed. The P458 peptide is shown in cartoon representation and colored in green. Diffraction data collection and refinement statistics of P458:CaM crystal structures 6YNS and 6YNU are given in Table S5.

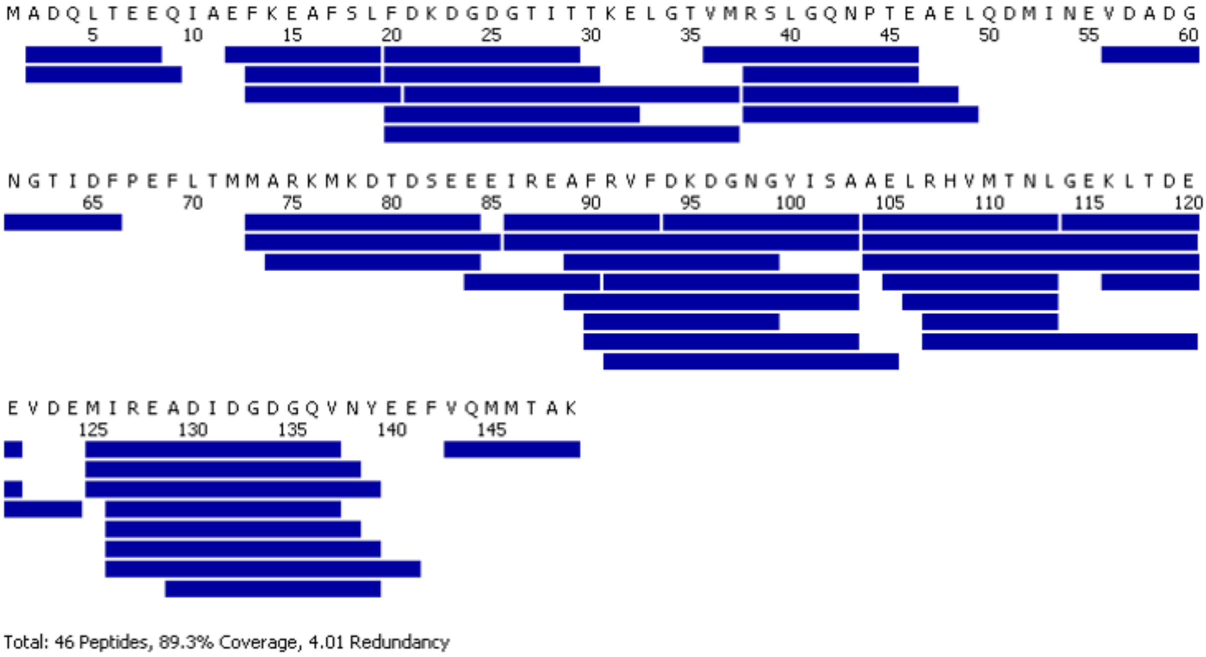


**Figure S10A:** Peptide map used to analyze the deuterium content in CaM. CaM was digested with immobilized pig pepsin at 20°C and pH 2.5 for 2 min. Each blue bar corresponds to a unique peptide selected for HDX-MS analysis. A final linear sequence coverage 89.3% (4.01 redundancy) was achieved.

*Figure S10 continued next page*

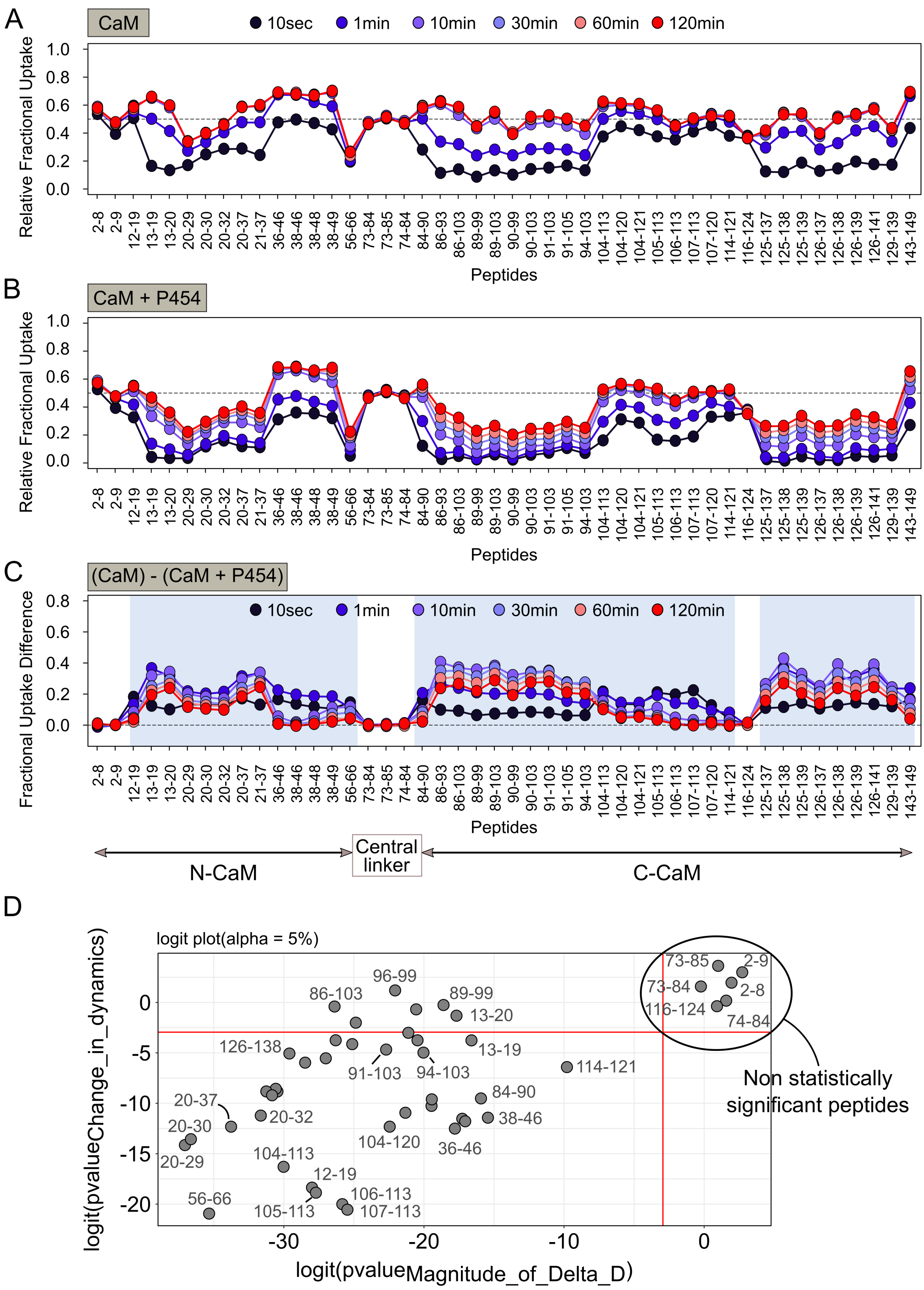


**Figure S10 continued.** Effect of P454 binding on the HDX-MS behavior of full-length human CaM. Fractional uptake profiles of CaM alone (**A**) and in the presence of a 4-fold molar excess P454 (**B**). The deuterium content calculated for each peptide and condition and at each time point is plotted as a function of peptide position. Each dot corresponds to the average value of three independent replicates. (**C**) Differential fractional uptake plot showing the uptake difference between full-length human CaM alone and in the presence of a 2-fold molar excess P454 for all peptides and labeling time points. The binding of P454 induces significant reduction of deuterium uptake throughout both N- and C-terminal of CaM.

Regions displaying statistically significant variations of deuterium uptake in the presence of P454 (Wald test, $p < 0.05$) are highlighted in light blue. (**D**) Logit representation showing the statistical results obtained with MEMHDX for each peptide (gray dots) of CaM in the presence of P454. The FDR value was set to 5% (red lines). Peptides displaying no statistically significant difference of deuterium uptake between states are clustered on the right hand corner of the plot.

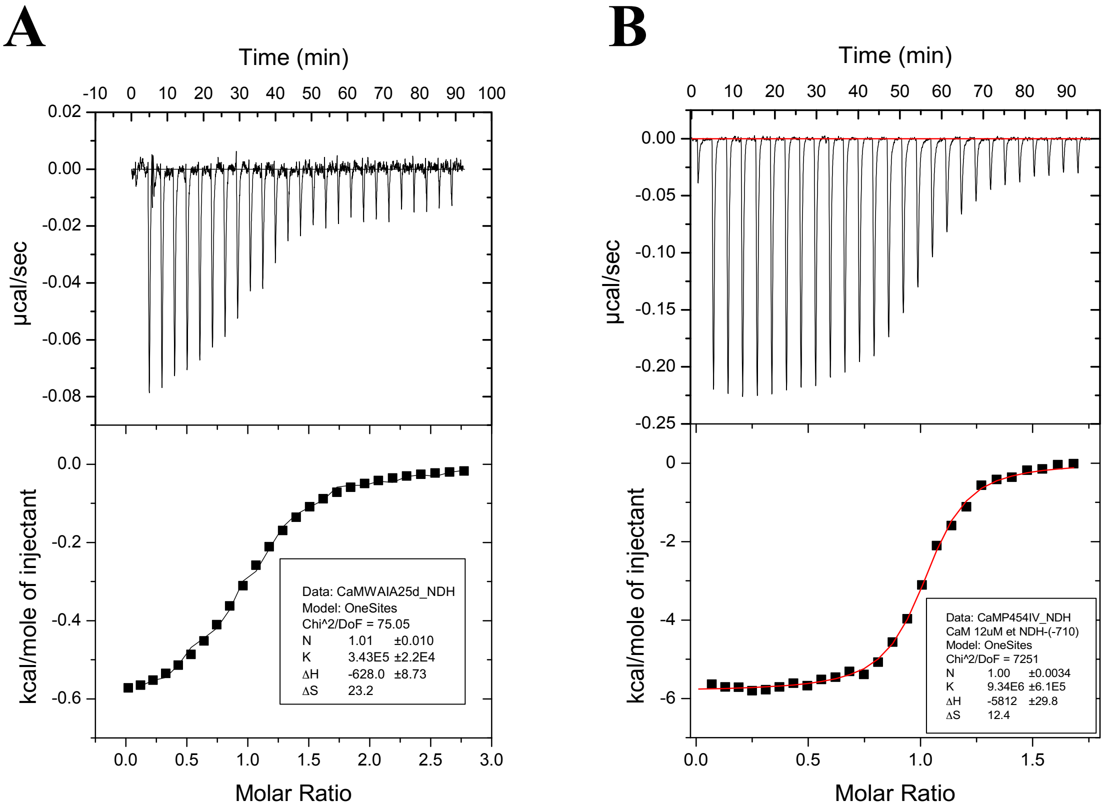


**Figure S11:** Isothermal titration calorimetry of holo-CaM by P454-derived peptides performed on a VP-ITC instrument. The P454$_{W458A-I479A}$ (**A**) and P454$_{I479V}$ (**B**) peptides are loaded in the syringe at 130 µM and injected in the cell containing holo-CaM at 10 µM. The free energy values of peptide:CaM interaction are listed in Table S3.

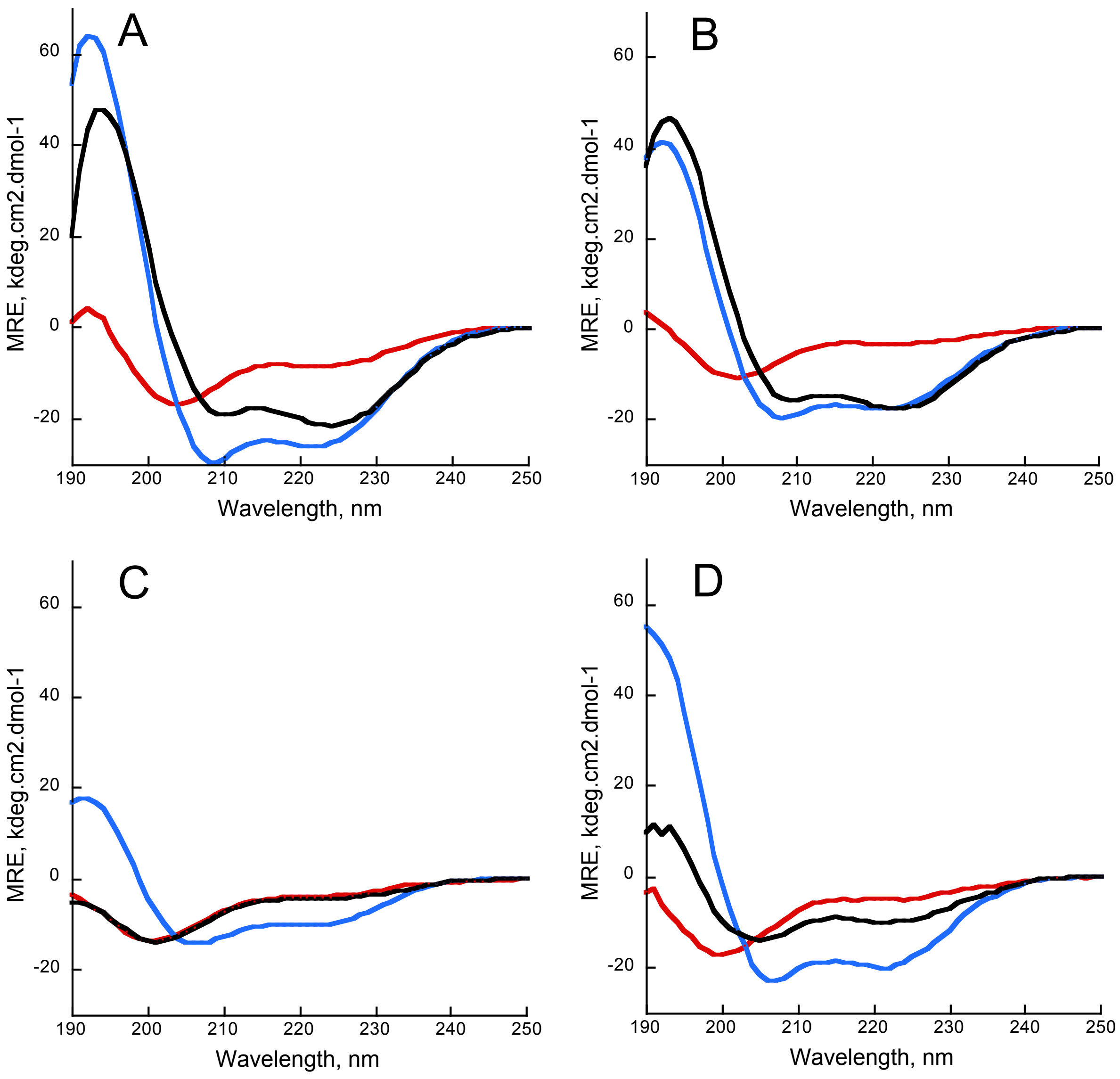


**Figure S12**: Synchrotron radiation circular dichroism in the far-UV region of P454-derived peptides. The far-UV CD spectra of the peptides have been acquired at 25°C in buffer A (red), in the presence of 20% TFE (blue) or in the presence of SUVs composed of POPC:POPG 8:2 at 2 mM lipids (black). Far-UV CD spectra of the following peptides are shown: P454 (**A**), P454 L481A (**B**), P454 I479A (**C**) and P454 R12Q (**D**). CD units are expressed as mean residue ellipticity (MRE)/1000, i.e., in Kdeg.cm$^2$.dmol$^{-1}$. The secondary structure content of the peptides is obtained from the deconvolution of the far-UV CD spectra (see Table S8) using the BestSel software [17] available at http://bestsel.elte.hu/index.php.

**Figure S13: Map of pCACTw11 plasmid and DNA sequence of encoded CyaA gene**

**A) pCACTw11 plasmid map**

Schematic map of pCACTw11 plasmid highlighting the regions

The open reading frame coding for the wild type CyaA is in orange with the segment located between the BstBI and NcoI sites replaced by a synthetic DNA fragment highlighted in magenta. The P454 coding region is in blue.

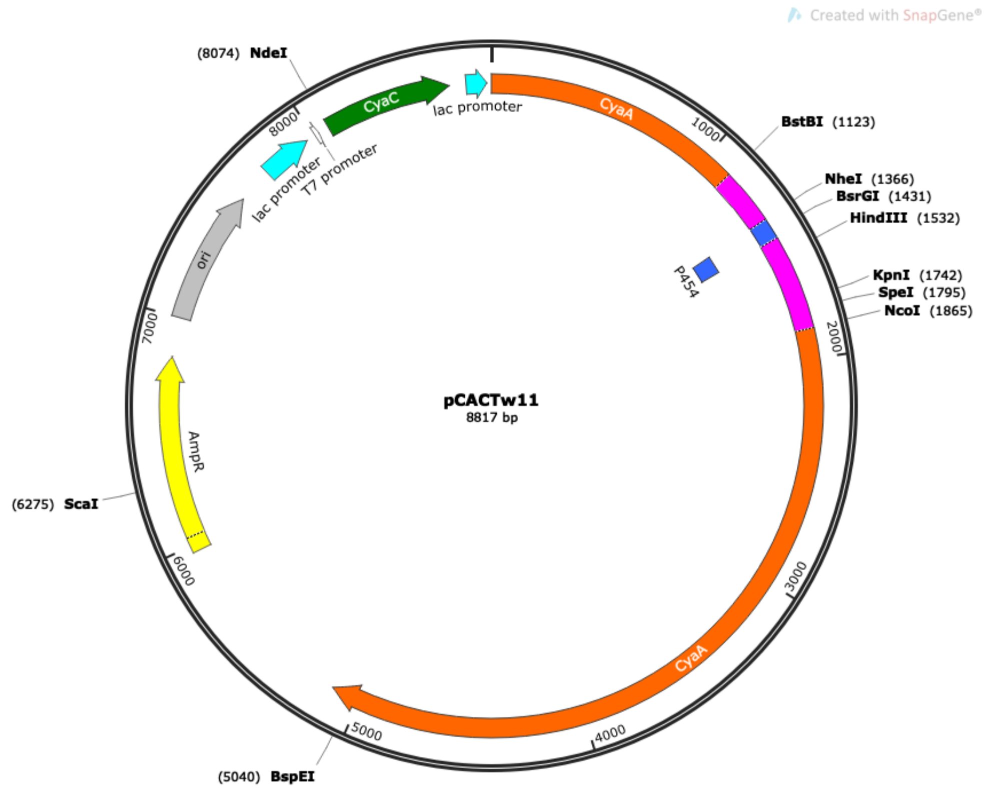


**B) CyaA WT open reading frame in pCACTw11 plasmid**

The full CyaA open reading frame is shown below with the synthetic DNA fragment located between the BstBI and NcoI sites in magenta. The P454 coding region is in blue. The bold underlined sequences correspond to the BstBI, NheI, KpnI and NcoI restriction sites that were used for subcloning the various synthetic DNA fragment encoding the modified CyaA toxins produced in this work.

ATGACCATGCAGCAATCGCATCAGGCTGGTTACGCAAACGCCGCCGACCGGGAG
TCTGGCATCCCCGCAGCCGTACTCGATGGCATCAAGGCCGTGGCGAAGGAAAAA
AACGCCACATTGATGTTCCGCCTGGTCAACCCCCATTCCACCAGCCTGATTGCCG
AAGGGGTGGCCACCAAAGGATTGGGCGTGCACGCCAAGTCGTCCGATTGGGGGT
TGCAGGCGGGCTACATTCCCGTCAACCCGAATCTTTCCAAACTGTTCGGCCGTGC
GCCCGAGGTGATCGCGCGGGCCGACAACGACGTCAACAGCAGCCTGGCGCATGG
CCATACCGCGGTCGACCTGACGCTGTCGAAAGAGCGGCTTGACTATCTGCGGCAA
GCGGGCCTGGTCACCGGCATGGCCGATGGCGTGGTCGCGAGCAACCACGCAGGC
TACGAGCAGTTCGAGTTTCGCGTGAAGGAAACCTCGGACGGGCGCTATGCCGTGC
AGTATCGCCGCAAGGGCGGCGACGATTTCGAGGCGGTCAAGGTGATCGGCAATG

CCGCCGGTATTCCACTGACGGCGGATATCGACATGTTCGCCATTATGCCGCATCT
GTCCAACTTCCGCGACTCGGCGCGCAGTTCGGTGACCAGCGGCGATTCGGTGACC
GATTACCTGGCGCGCACGCGGCGGGCCGCCAGCGAGGCCACGGGCGGCCTGGAT
CGCGAACGCATCGACTTGTTGTGGAAAATCGCTCGCGCCGGCGCCCGTTCCGCAG
TGGGCACCGAGGCGCGTCGCCAGTTCCGCTACGACGGCGACATGAATATCGGCG
TGATCACCGATTTCGAGCTGGAAGTGCGCAATGCGCTGAACAGGCGGGCGCACG
CCGTCGGCGCGCAGGACGTGGTCCAGCATGGCACTGAGCAGAACAATCCTTTCCC
GGAGGCAGATGAGAAGATTTTCGTCGTATCGGCCACCGGTGAAAGCCAGATGCT
CACGCGCGGGCAACTGAAGGAATACATTGGCCAGCAGCGCGGCGAGGGCTATGT
CTTCTACGAGAACCGTGCATACGGCGTGGCGGGGAAAAGCCTGTTCGACGATGG
GCTGGGAGCCGCGCCCGGCGTGCCGAGCGGACG**TTCGAA**GTTCTCGCCGGATGT
ACTGGAAACCGTTCCGGCGTCACCAGGTTTGCGTCGTCCTTCGCTGGGTGCAGTT
GAACGTCAGGATTCTGGTTATGACTCTCTTGATGGTGTTGGTTCGCGTTCTTTCTC
ATTAGGTGAGGTTTCTGACATGGCTGCAGTTGAAGCAGCTGAACTGGAAATGACT
CGTCAAGTTTTACACGCTGGTGCACGTCAGGACGATGCTGAGCCAGGTGTTTCTG
GT**GCTAGC**GCTCATTGGGGTCAGCGTGCTCTGCAAGGTGCTCAGGCTGTTGCAGC
TGCTCAGCGTCTTGTACATGCTATTGCACTGATGACTCAATTCGGGCGCGCCGGTT
CTACTAACACTCCACAGGAAGCTGCTTCTCTTTCTGCTGCAGTTTTCGGTCTGGGT
GAAGCTTCTAGCGCAGTAGCTGAAACTGTTTCTGGTTTTTCCGTGGTTCTTCACG
CTGGGCTGGCGGTTTCGGCGTGGCTGGTGGCGCAATGGCTCTGGGAGGTGGTATC
GCTGCAGCTGTTGGTGCTGGTATGTCTTTGACTGATGACGCTCCAGCTGGTCAAA
AAGCTGCAGCTGGTGCTGAAATTGCTCTGCAGTTAACAGGT**GGTACC**GTTGAGCT
GGCTTCTTCCATTGCTTTGGCCTTAGCTGCAGCTCGTGGTGTAACTAGTGGCTTGC
AGGTTGCCGGTGCTTCTGCAGGTGCGGCTGCTGGTGCATTGGCCGCGGCGCTCAG
TC**CCATGG**AGATCTACGGCCTGGTGCAGCAATCGCACTATGCGGATCAGCTGGA
CAAGCTGGCGCAGGAATCGAGCGCATACGGTTACGAGGGCGACGCCTTGCTGGC
CCAGCTGTATCGCGACAAGACGGCCGCCGAGGGCGCCGTCGCCGGCGTCTCCGC
CGTCCTGAGCACGGTGGGGGCGGCGGTGTCGATCGCCGCGGCGGCCAGCGTGGT
AGGGGCCCCGGTGGCGGTGGTCACTTCCTTGCTGACCGGGGCTCTCAACGGCATC
CTGCGCGGCGTGCAGCAGCCCATCATCGAAAAGCTGGCCAACGATTACGCTCGC
AAGATCGACGAGCTGGGCGGGCCGCAAGCGTACTTCGAGAAAAACCTGCAGGCG
CGTCACGAACAACTGGCCAATTCGGACGGCCTACGGAAAATGCTGGCCGACCTG
CAGGCCGGTTGGAACGCCAGCAGCGTGATCGGGGTGCAGACGACAGAGATCTCC
AAGTCGGCGCTCGAACTGGCCGCCATTACCGGCAACGCGGACAACCTGAAATCC
GTCGACGTGTTCGTGGACCGCTTCGTCCAGGGCGAGCGGGTGGCCGGCCAGCCG
GTGGTCCTCGACGTCGCCGCCGGCGGCATCGATATCGCCAGCCGCAAGGGCGAG
CGGCCGGCGCTGACGTTCATCACGCCGCTGGCCGCGCCAGGAGAAGAGCAGCGC
CGGCGCACGAAAACGGGCAAGAGCGAATTCACCACATTCGTCGAGATCGTGGGC
AAGCAGGACCGCTGGCGCATCCGGGACGGCGCGGCCGACACCACCATCGATCTG
GCCAAGGTGGTGTCGCAACTGGTCGACGCCAATGGCGTGCTCAAGCACAGCATC
AAACTGGATGTGATCGGCGGAGATGGCGATGACGTCGTGCTTGCCAATGCTTCGC
GCATCCATTATGACGGCGGCGCGGGCACCAACACGGTCAGCTATGCCGCCCTGG
GTCGACAGGATTCCATTACCGTGTCCGCCGACGGGGAACGTTTCAACGTGCGCAA
GCAGTTGAACAACGCCAACGTGTATCGCGAAGGCGTGGCTACCCAGACAACCGC
CTACGGCAAGCGCACGGAGAATGTCCAATACCGCCATGTCGAGCTGGCCCGTGTC
GGGCAAGTGGTGGAGGTCGACACGCTCGAGCATGTGCAGCACATCATCGGCGGG
GCCGGCAACGATTCGATCACCGGCAATGCGCACGACAACTTCCTAGCCGGCGGG
TCGGGCGACGACAGGCTGGATGGCGGCGCCGGCAACGACACCCTGGTTGGCGGC
GAGGGCCAAAACACGGTCATCGGCGGCGCCGGCGACGACGTATTCCTGCAGGAC
CTGGGGGTATGGAGCAACCAGCTCGATGGCGGCGCGGGCGTCGATACCGTGAAG

TACAACGTGCACCAGCCTTCCGAGGAGCGCCTCGAACGCATGGGCGACACGGGC
ATCCATGCCGATCTTCAAAAGGGCACGGTCGAGAAGTGGCCGGCCCTGAACCTGT
TCAGCGTCGACCATGTCAAGAATATCGAGAATCTGCACGGCTCCCGCCTAAACGA
CCGCATCGCCGGCGACGACCAGGACAACGAGCTCTGGGGCCACGATGGCAACGA
CACGATACGCGGCCGGGGCGGCGACGACATCCTGCGCGGCGGCCTGGGCCTGGA
CACGCTGTATGGCGAGGACGGCAACGACATCTTCCTGCAGGACGACGAGACCGT
CAGCGATGACATCGACGGCGGCGCGGGGCTGGACACCGTCGACTACTCCGCCAT
GATCCATCCAGGCAGGATCGTTGCGCCGCATGAATACGGCTTCGGGATCGAGGC
GGACCTGTCCAGGGAATGGGTGCGCAAGGCGTCCGCGCTGGGCGTGGACTATTA
CGATAATGTCCGCAATGTCGAAAACGTCATCGGTACGAGCATGAAGGATGTGCTC
ATCGGCGACGCGCAAGCCAATACCCTGATGGGCCAGGGCGGCGACGATACCGTG
CGCGGCGGCGACGGCGATGATCTGCTGTTCGGCGGCGACGGCAACGACATGCTG
TATGGCGACGCCGGCAACGACACCCTCTACGGGGGGCTGGGCGACGATACCCTT
GAAGGCGGCGCGGGCAACGATTGGTTCGGCCAGACGCAGGCGCGCGAGCATGAC
GTGCTGCGCGGCGGAGATGGGGTGGATACCGTCGATTACAGCCAGACCGGCGCG
CATGCCGGCATTGCCGCGGGTCGCATCGGGCTGGGCATCCTGGCTGACCTGGGCG
CCGGCCGCGTCGACAAGCTGGGCGAGGCCGGCAGCAGCGCCTACGATACGGTTT
CCGGTATCGAGAACGTGGTGGGCACGGAACTGGCCGACCGCATCACGGGCGATG
CGCAGGCCAACGTGCTGCGCGGCGCGGGTGGCGCCGACGTGCTTGCGGGCGGCG
AGGGCGACGATGTGCTGCTGGGCGGCGACGGCGACGACCAGCTGTCGGGCGACG
CCGGACGCGATCGCTTGTACGGCGAAGCCGGTGACGACTGGTTCTTCCAGGATGC
CGCCAATGCCGGCAATCTGCTCGACGGCGGCGACGGCCGCGATACCGTGGATTTC
AGCGGCCCGGGCCGGGGCCTCGACGCCGGCGCAAAGGGCGTATTCCTGAGCTTG
GGCAAGGGGTTCGCCAGCCTGATGGACGAACCCGAAACCAGCAACGTGTTGCGC
AATATCGAGAACGCCGTGGGCAGCGCGCGTGATGACGTGCTGATCGGCGACGCA
GGCGCCAACGTCCTCAATGGCCTGGCGGGCAACGACGTGCTGTCCGGCGGCGCT
GGCGACGATGTGCTGCTGGGCGACGAGGGCTCGGACCTGCTCAGCGGCGATGCG
GGCAACGACGATCTGTTCGGCGGGCAGGGCGATGATACTTATCTGTTCGGGGTCG
GGTACGGGCACGACACGATCTACGAATCGGGCGGCGGCCATGACACCATCCGCA
TCAACGCGGGGGCGGACCAGCTGTGGTTCGCGCGCCAGGGCAACGACCTGGAGA
TCCGCATTCTCGGCACCGACGATGCACTTACCGTGCACGACTGGTATCGCGACGC
CGATCACCGGGTGGAAATCATCCATGCCGCCAACCAGGCGGTAGACCAGGCAGG
CATCGAAAAGCTGGTCGAGGCAATGGCGCAGTATCCGGACCCCGGCGCGGCGGC
GGCTGCCCCGCCGGCGGCGCGCGTGCCGGACACGCTGATGCAGTCCCTGGCTGTC
AACTGGCGCTGA

*Figure S13 continued next page*

**C) Synthetic DLTW11 DNA fragment**

Sequence of the synthetic DNA fragment used for pCACTw11 construction. In black letters are the extra sequence needed for Hifi cloning while the codon-optimized CyaA sequence located between the BstBI and NcoI sites is in magenta. The P454 coding region is in blue. The bold underlined sequences correspond to the BstBI, NheI, KpnI and NcoI restriction sites used for subcloning.

**CCCGGCGTGCCGAGCGGACGTTCGAA**GTTCTCGCCGGATGTACTGGAAACCGT
TCCGGCGTCACCAGGTTTGCGTCGTCCTTCGCTGGGTGCAGTTGAACGTCAGGAT
TCTGGTTATGACTCTCTTGATGGTGTTGGTTCGCGTTCTTTCTCATTAGGTGAGGTT
TCTGACATGGCTGCAGTTGAAGCAGCTGAACTGGAAATGACTCGTCAAGTTTTAC
ACGCTGGTGCACGTCAGGACGATGCTGAGCCAGGTGTTTCTGGT**GCTAGC**GCTC
ATTGGGGTCAGCGTGCTCTGCAAGGTGCTCAGGCTGTTGCAGCTGCTCAGCGTCT
TGTACATGCTATTGCACTGATGACTCAATTCGGGCGCGCCGGTTCTACTAACACT
CCACAGGAAGCTGCTTCTCTTTCTGCTGCAGTTTTCGGTCTGGGTGAAGCTTCTAG
CGCAGTAGCTGAAACTGTTTCTGGTTTTTTCCGTGGTTCTTCACGCTGGGCTGGCG
GTTTCGGCGTGGCTGGTGGCGCAATGGCTCTGGGAGGTGGTATCGCTGCAGCTGT
TGGTGCTGGTATGTCTTTGACTGATGACGCTCCAGCTGGTCAAAAAGCTGCAGCT
GGTGCTGAAATTGCTCTGCAGTTAACAGGT**GGTACC**GTTGAGCTGGCTTCTTCCA
TTGCTTTGGCCTTAGCTGCAGCTCGTGGTGTAACTAGTGGCTTGCAGGTTGCCGGT
GCTTCTGCAGGTGCGGCTGCTGGTGCATTGGCCGCGGCGCTCAGTC**CCATGGAG**
**ATCTACGGCCTGGTGCAG**

**D) Synthetic DLTW16 DNA fragment coding for CyaA$_{Mut1}$.**

Sequence of the synthetic DNA fragment used for pCACTw16 construction. The color code is same as above. Highlighted in yellow are the 6 codons that were changed.

**CCCGGCGTGCCGAGCGGACGTTCGAA**GTTCTCGCCGGATGTACTGGAAACCGT
TCCGGCGTCACCAGGTTTGCGTCGTCCTTCGCTGGGTGCAGTTGAACGTCAGGAT
TCTGGTTATGACTCTCTTGATGGTGTTGGTTCGCGTTCTTTCTCATTAGGTGAGGTT
TCTGACATGGCTGCAGTTGAAGCAGCTGAACTGGAAATGACTCGTCAAGTTTTAC
ACGCTGGTGCACGTCAGGACGATGCTGAGCCAGGTGTTTCTGGT**GCTAGC**GCTC
ATTGGGGTCAGGAAGCTGCACAAGGTGCTCAGGCTGTTGCAGCTGCTCAGGAAG
CAGTATCTGCTGCAGCACTGATGACTCAATTCGGGCGCGCCGGTTCTACTAACAC
TCCACAGGAAGCTGCTTCTCTTTCTGCTGCAGTTTTCGGTCTGGGTGAAGCTTCTA
GCGCAGTAGCTGAAACTGTTTCTGGTTTTTTCCGTGGTTCTTCACGCTGGGCTGGC
GGTTTCGGCGTGGCTGGTGGCGCAATGGCTCTGGGAGGTGGTATCGCTGCAGCTG
TTGGTGCTGGTATGTCTTTGACTGATGACGCTCCAGCTGGTCAAAAAGCTGCAGC
TGGTGCTGAAATTGCTCTGCAGTTAACAGGT**GGTACC**GTTGAGCTGGCTTCTTCC
ATTGCTTTGGCCTTAGCTGCAGCTCGTGGTGTAACTAGTGGCTTGCAGGTTGCCG
GTGCTTCTGCAGGTGCGGCTGCTGGTGCATTGGCCGCGGCGCTCAGTC**CCATGGA**
**GATCTACGGCCTGGTGCAG**

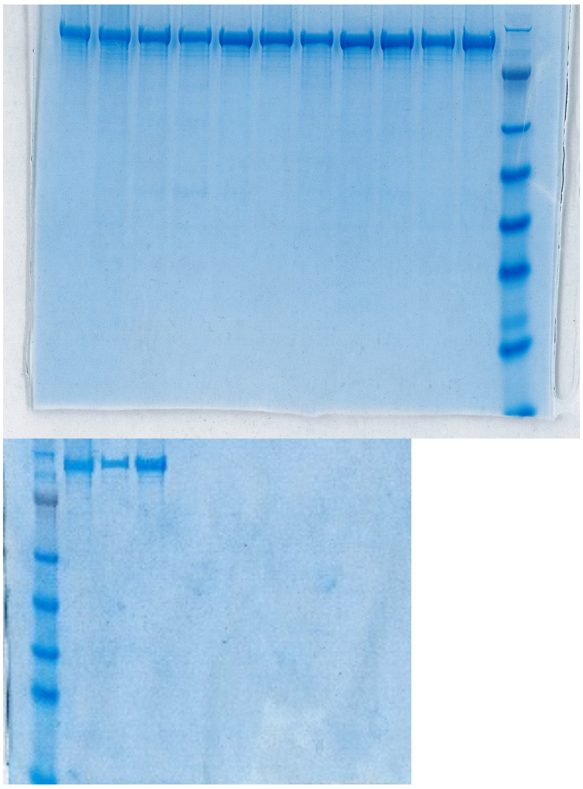

**Figure S14:** SDS-PAGE of purified recombinant CyaA proteins. Two µg of each recombinant CyaA protein are loaded on each line. First (top) SDS-PAGE, from left to right: CyaA WT (batch #0918); CyaA WT (batch #0719); CyaA$_{Mut6}$; CyaA$_{R12E}$; CyaA$_{Mut1}$; CyaA$_{Mut7}$; CyaA$_{R12Q}$; CyaA$_{Mut2}$; CyaA$_{Mut4}$; CyaA$_{R12A}$; CyaA$_{R12K}$ and Ladder SeeBlue Plus. Second (bottom) SDS-PAGE, from left to right: Ladder SeeBlue Plus; CyaA WT (batch #0918); CyaA$_{Mut3}$ and CyaA$_{Mut5}$.

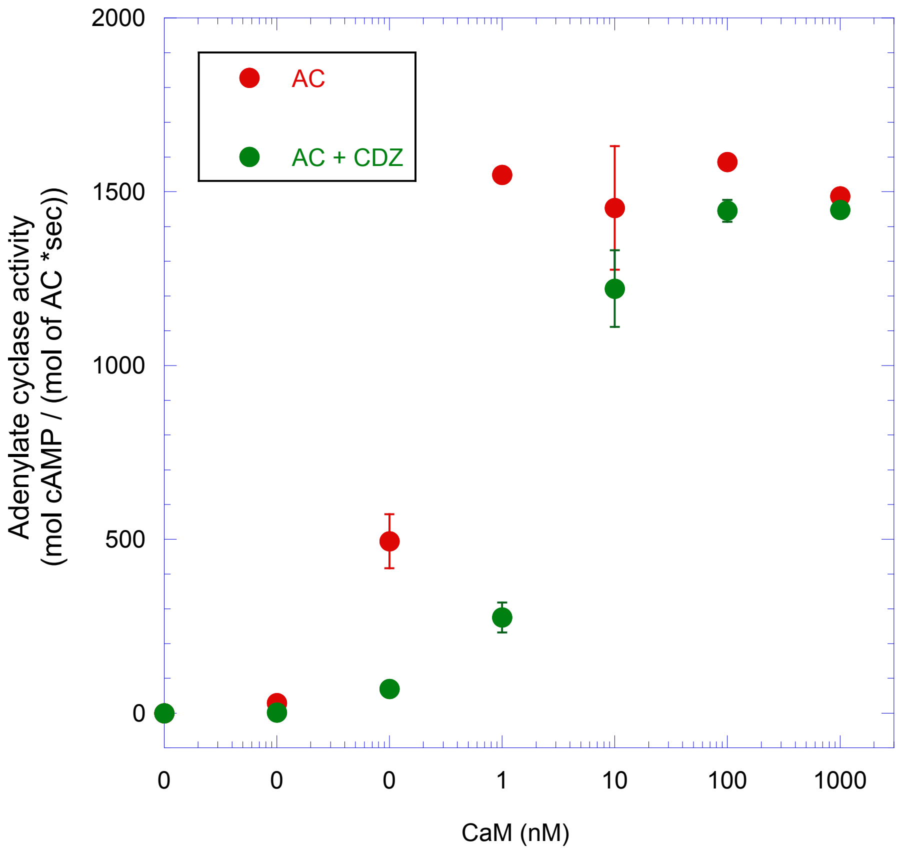


**Figure S15:** CaM-dependent activation of AC with or without calmidazolium (CDZ). The adenylate cyclase activity of AC384GK (encompassing the full catalytic domain of CyaA) was measured in the presence of the indicated concentrations of CaM (red symbols) and in the presence of 10 µM calmidazolium (CDZ, green symbols) as previously described in Vougier *et al.* 2004 at 30 °C, pH 8.0 and 0.1 mM $CaCl_2$. AC384GK was at 0.2 nM. The adenylate cyclase activity is expressed as mol of cAMP per sec per mol of AC384GK.

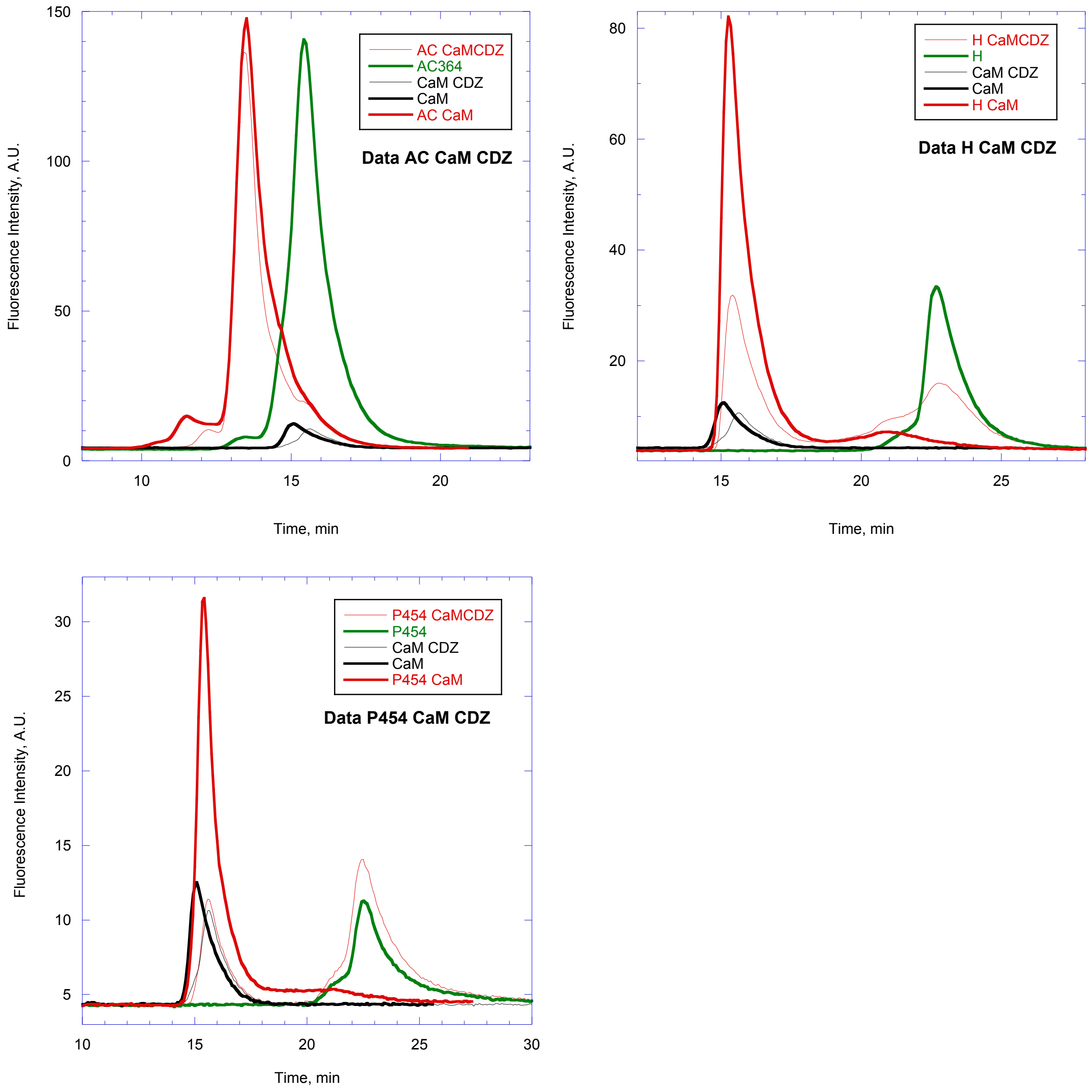


**Figure S16**: Size exclusion chromatography of AC, H and P454 in the absence and presence of CaM and CDZ followed by tryptophan intrinsic fluorescence (excitation: 280 nm; emission: 340 nm). AC, H and P454 contain tryptophan residues. CaM does not contain any tryptophan residue but 8 phenylalanine residues, providing a residual fluorescence signal. The chromatograms of AC, H and P454 are green. CaM is dark and the complexes are red. In the presence of CDZ, the chromatograms are identified by hairline traces (CaM:CDZ is in light dark and complexes:CDZ are in light red). The molecules were mixed at the following finale concentrations: AC, H and P454 at 10 µM, CaM at 15 µM and CDZ at 40 µM. The chromatograms of CaM alone and CaM:CDZ shown in the 3 figures are from the same samples. The SEC experiments were performed on a TSKgel G3000SW 7.5 mm ID * 300 mm. The buffer is 20 mM Hepes, 150 mM NaCl, 2 mM CaCl2, pH 7.4.

13. Sotomayor Perez, A. C.; Karst, J. C.; Davi, M.; Guijarro, J. I.; Ladant, D.; Chenal, A., Characterization of the regions involved in the calcium-induced folding of the

intrinsically disordered RTX motifs from the bordetella pertussis adenylate cyclase toxin. *Journal of molecular biology* **2010,** *397* (2), 534-49.
14. White, S. H.; Wimley, W. C., Membrane protein folding and stability: physical principles. *Annual review of biophysics and biomolecular structure* **1999,** *28*, 319-65.
15. Schuck, P., Size-distribution analysis of macromolecules by sedimentation velocity ultracentrifugation and lamm equation modeling. *Biophys J* **2000,** *78* (3), 1606-19.
16. Micsonai, A.; Wien, F.; Kernya, L.; Lee, Y. H.; Goto, Y.; Refregiers, M.; Kardos, J., Accurate secondary structure prediction and fold recognition for circular dichroism spectroscopy. *Proc Natl Acad Sci U S A* **2015,** *112* (24), E3095-103.
17. Micsonai, A.; Wien, F.; Bulyaki, E.; Kun, J.; Moussong, E.; Lee, Y. H.; Goto, Y.; Refregiers, M.; Kardos, J., BeStSel: a web server for accurate protein secondary structure prediction and fold recognition from the circular dichroism spectra. *Nucleic Acids Res* **2018,** *46* (W1), W315-W322.
18. Karimova, G.; Fayolle, C.; Gmira, S.; Ullmann, A.; Leclerc, C.; Ladant, D., Charge-dependent translocation of Bordetella pertussis adenylate cyclase toxin into eukaryotic cells: implication for the in vivo delivery of CD8(+) T cell epitopes into antigen-presenting cells. *Proc Natl Acad Sci U S A* **1998,** *95* (21), 12532-7.
19. Rogel, A.; Hanski, E., Distinct steps in the penetration of adenylate cyclase toxin of Bordetella pertussis into sheep erythrocytes. Translocation of the toxin across the membrane. *J Biol Chem* **1992,** *267* (31), 22599-605.
20. David, G.; Pérez, J., Combined sampler robot and high-performance liquid chromatography: a fully automated system for biological small-angle X-ray scattering experiments at the Synchrotron SOLEIL SWING beamline. *J. Appl. Cryst.* **2009,** *42*, 892-900.
21. Trewhella, J.; Duff, A. P.; Durand, D.; Gabel, F.; Guss, J. M.; Hendrickson, W. A.; Hura, G. L.; Jacques, D. A.; Kirby, N. M.; Kwan, A. H.; Perez, J.; Pollack, L.; Ryan, T. M.; Sali, A.; Schneidman-Duhovny, D.; Schwede, T.; Svergun, D. I.; Sugiyama, M.; Tainer, J. A.; Vachette, P.; Westbrook, J.; Whitten, A. E., 2017 publication guidelines for structural modelling of small-angle scattering data from biomolecules in solution: an update. *Acta Crystallogr D Struct Biol* **2017,** *73* (Pt 9), 710-728.
22. Franke, D.; Petoukhov, M. V.; Konarev, P. V.; Panjkovich, A.; Tuukkanen, A.; Mertens, H. D. T.; Kikhney, A. G.; Hajizadeh, N. R.; Franklin, J. M.; Jeffries, C. M.; Svergun, D. I., ATSAS 2.8: a comprehensive data analysis suite for small-angle scattering from macromolecular solutions. *J Appl Crystallogr* **2017,** *50* (Pt 4), 1212-1225.
23. Franke, D.; Jeffries, C. M.; Svergun, D. I., Correlation Map, a goodness-of-fit test for one-dimensional X-ray scattering spectra. *Nature methods* **2015,** *12* (5), 419-22.
24. Guinier, A., La diffraction des rayons X aux très petits angles; application à l'étude de phénomènes ultramicroscopiques. *Ann Phys (Paris)* **1939,** *12*, 161–237.
25. Svergun, D. I., Determination of the Regularization Parameter in Indirect - Transform Methods Using Perceptual Criteria. *J. Appl. Cryst.* **1992,** *25*, 495-503.
26. Franke, D.; Svergun, D. I., DAMMIF, a program for rapid ab-initio shape determination in small-angle scattering. *J Appl Crystallogr* **2009,** *42* (Pt 2), 342-346.
27. Svergun, D. I., Restoring low resolution structure of biological macromolecules from solution scattering using simulated annealing. *Biophysical Journal* **1999,** *76* (6), 2879-2886.
28. Evrard, G.; Mareuil, F.; Bontems, F.; Sizun, C.; Perez, J., DADIMODO: a program for refining the structure of multidomain proteins and complexes against small-angle scattering data and NMR-derived restraints. *J Appl Crystallogr* **2011,** *44*, 1264-1271.

29. Weber, P.; Pissis, C.; Navaza, R.; Mechaly, A. E.; Saul, F.; Alzari, P. M.; Haouz, A., High-Throughput Crystallization Pipeline at the Crystallography Core Facility of the Institut Pasteur. *Molecules* **2019,** *24* (24).
30. Vonrhein, C.; Flensburg, C.; Keller, P.; Sharff, A.; Smart, O.; Paciorek, W.; Womack, T.; Bricogne, G., Data processing and analysis with the autoPROC toolbox. *Acta Crystallogr D Biol Crystallogr* **2011,** *67* (Pt 4), 293-302.
31. Mccoy, A. J.; Grosse-Kunstleve, R. W.; Adams, P. D.; Winn, M. D.; Storoni, L. C.; Read, R. J., Phaser crystallographic software. *J Appl Crystallogr* **2007,** *40*, 658-674.
32. Emsley, P.; Cowtan, K., Coot: model-building tools for molecular graphics. *Acta Crystallogr D* **2004,** *60*, 2126-2132.
33. Bricogne, G.; Blanc, E.; Brandl, M.; Flensburg, C.; Keller, P., BUSTER version 2.11.1. *Cambridge, UK: Global Phasing Ltd.* **2011**.
34. Hourdel, V.; Volant, S.; O'Brien, D. P.; Chenal, A.; Chamot-Rooke, J.; Dillies, M. A.; Brier, S., MEMHDX: An interactive tool to expedite the statistical validation and visualization of large HDX-MS datasets. *Bioinformatics (Oxford, England)* **2016**.
35. Brokx, R. D.; Lopez, M. M.; Vogel, H. J.; Makhatadze, G. I., Energetics of target peptide binding by calmodulin reveals different modes of binding. *J Biol Chem* **2001,** *276* (17), 14083-91.
36. Chattopadhyaya, R.; Meador, W. E.; Means, A. R.; Quiocho, F. A., Calmodulin structure refined at 1.7 A resolution. *Journal of molecular biology* **1992,** *228* (4), 1177-92.
